# Social Information use Shapes the Coevolution of Sociality and Virulence

**DOI:** 10.1101/2020.10.02.323451

**Authors:** Ben Ashby, Damien R Farine

## Abstract

Social contacts can facilitate the spread of both survival-related information and infectious diseases, but little is known about how these processes combine to shape host and parasite evolution. Here, we use a theoretical model that captures both infection and information transmission processes to investigate how host sociality (contact effort) and parasite virulence (disease-associated mortality rate) (co)evolve. We show that selection for sociality (and in turn, virulence) depends on both the intrinsic costs and benefits of social information and infection as well as their relative prevalence in the population. Specifically, greater sociality and lower virulence evolve when the risk of infection is either low or high and social information is neither very common nor too rare. Lower sociality and higher virulence evolve when the prevalence patterns are reversed. When infection and social information are both at moderate levels in the population, the direction of selection depends on the relative costs and benefits of being infected or informed. We also show that sociality varies inversely with virulence, and that parasites may be unable to prevent runaway selection for higher contact efforts. Together, these findings provide new insights for our understanding of group living and how apparently opposing ecological processes can influence the evolution of sociality and virulence in a range of ways.

## Introduction

Animals exhibit strong variation in sociality, both within (Watters and Sih 2005; Pike et al. 2008; Croft et al. 2009; Tanner and Jackson 2012; Aplin et al. 2013, 2015b; Weber et al. 2013) and between (Sachser 1986; Barton et al. 1996) populations. Some individuals have relatively few, potentially stable associations (e.g. non-gregarious species or those that form pair bonds) while others have a greater number of interactions that may be more transient (e.g. herding behaviour or roaming individuals) (Couzin 2006; Silk et al. 2014; Farine et al. 2015b). Crucially, what constitutes a good social strategy under one set of ecological conditions may be a poor choice under different circumstances (Krause and Ruxton 2002), hence it is important to understand how changes in the environment shape social behaviour. Depending on the ecological circumstances, sociality may be associated with various costs (e.g. higher risk of infection or greater competition for resources) and benefits (e.g. information exchange, increased mating opportunities, or symbiont transmission), which trade-off to determine whether selection favours increased sociality (Romano et al. 2021). While sociality can take a variety of forms (e.g. contact effort, rate, turnover, degree, or strength), in this study we shall focus on contact effort.

Intuitively, if sociality confers diminishing benefits but accelerating costs, then the population should evolve to an evolutionarily stable level of sociality. This is straightforward when costs and benefits are fixed, so that a given level of sociality always incurs the same cost or provides the same benefit. For example, living in a larger groups may lead to foraging benefits (Cantor et al. 2020) or a reduced risk of predation (Kenward 1978). However, when the costs and benefits are dynamic, for example due to variation in infection prevalence (dynamic cost) and information prevalence (dynamic benefit), feedbacks may exist between sociality and the likelihood of realising an associated cost or benefit (Cantor et al. 2021b). For example, more social members of a population may experience greater exposure to parasites (Lloyd-Smith et al. 2005; Eames and Keeling 2006; Ashby and Gupta 2013) and access to social information (Aplin et al. 2012, 2015a; Snijders et al. 2021) than those that form fewer contacts or have more stable associations (Evans et al. 2020). However, the sociality of the population will affect the prevalence of infection or social information in the population, and therefore the likelihood of incurring a cost or realising a benefit of sociality (Romano et al. 2020). For example, a study on wild house finches found that individuals that visited bird feeders more often (an outcome of information acquisition (Hillemann et al. 2020)) were more likely to be infected by the bacterial pathogen *Mycoplasma gallisepticum* (Adelman et al. 2015). Thus, while we should expect parasitism and social information to influence the evolution of sociality, we must also consider how the (co)evolution of sociality and virulence shapes the dynamics of these cost/benefit processes due to eco-evolutionary feedbacks (Ashby et al. 2019). An outstanding question, which we address in the present study, is how dynamic costs (in terms of parasite transmission) and benefits (in terms of information transmission) mediate the (co)evolution of sociality and virulence.

The evolution of virulence is intrinsic to the information-disease trade-off when being social. Mortality virulence (as opposed to sterility virulence, which we do not consider here) is typically costly to the parasite, as it reduces the average infectious period and hence limits the time available for transmission (note that this is not always the case, for example if host death is required for transmission). Naively, one may therefore assume that selection should favour lower virulence. However, parasites usually need to harm their hosts to grow and reproduce, and so there may be a trade-off between virulence and onwards transmission, leading to selection for an intermediate level of virulence (reviewed in Alizon et al. 2009). As a result, entwined in the evolution of sociality in hosts are virulence-transmission trade-offs, which will not only determine the direction of selection but also the prevalence of infection in the host population.

Classical predictions for parasite evolution suggest that higher levels of population mixing should select for higher virulence when it is positively correlated with transmission (Ewald 1994). Yet this does not necessarily mean that higher levels of sociality will select for higher virulence, as this depends on factors such as population structure (Evans et al. 2020; Cantor et al. 2021b; Romano et al. 2021). Bonds et al. (2005) showed that the coevolution of contact effort with virulence may instead lead to an inverse relationship between these traits. They also showed that sociality (contact rate) is minimised for intermediate infection prevalence, and therefore higher infection prevalence can select for greater sociality. This is because there is little chance of avoiding infection when prevalence is high, and so the marginal costs of greater sociality are low. In a related study, Prado et al. (2009) used a stochastic individual-based network model to show that coevolution between host contact number (degree) and parasite virulence (disease-associated mortality rate) can lead to fluctuating selection in both traits, with the cost of sociality depending both on the virulence of the parasite and its prevalence in the population. However, few theoretical studies have considered how eco-evolutionary feedbacks mediate the evolution of sociality (see Cantor et al. 2021 for a review), and as far as we are aware, these are the only two theoretical studies to date to explore the coevolution of sociality and virulence (Buckingham and Ashby 2022). While these studies demonstrate the importance of parasitism as a dynamic cost in the evolution of sociality, both assume that the benefits of sociality are fixed, so that each additional contact increases host fitness by a predetermined amount. Theoretical studies have yet to consider dynamic rather than fixed benefits of sociality.

One of the most prominent examples of a dynamic benefit of sociality is social information, which can provide individuals with knowledge about ephemeral resources or conditions, such as foraging or nesting sites, or predation risk (Doligez et al. 2002; Danchin et al. 2004; Valone 2007). This information can be transmitted through active signalling (Elgar 1986) or via inadvertent cues (Pöysä 1992; Galef and Giraldeau 2001). Empirical studies from wild songbirds have demonstrated that more social individuals are likely to receive information sooner than poorly connected members of the population (Aplin et al. 2012, 2015a; Snijders et al. 2021). We should therefore expect sociality to increase access to survival-related information, but as is the case with infection prevalence, the prevalence of information will depend on sociality, forming an eco-evolutionary feedback that causes the evolution of sociality to diverge from the fixed-benefit scenario.

While social information and parasite transmission share many similarities, their dynamics may occur over similar or very different timescales, which may affect selection on social strategies. Social information about predation or resource availability is expected to be of high value that rapidly decays over time (between seconds and days), whereas the risk of infection may vary more slowly and over longer periods of time (weeks, months or years). For example, the life cycle of a parasite can de-couple the transmission from social contacts by several weeks (Grear et al. 2013). In general, we should expect social information dynamics to be relatively fast compared to infection dynamics. The success of a particular social strategy is likely to depend on the relationship between these timescales, as well as wider contact patterns in the population, the value of social information, and the prevalence and severity of infection.

Here, we examine how social information mediates the evolution of sociality (contact effort) and virulence (disease-associated mortality rate). We present a mathematical model of social information and epidemiological dynamics, allowing host sociality to evolve with no inherent costs or benefits, and parasite virulence to evolve subject to a trade-off with transmission (i.e. lower virulence implies lower transmission). Evolution is determined purely by balancing access to social information, which reduces the mortality rate, and infectious disease caused by parasites, which increases the mortality rate. We consider two versions of the model, where social information dynamics occur on relatively fast or slow timescales, with fast information dynamics allowing for a separation of timescales with epidemiological dynamics. We show that selection favours increased sociality when social information is neither very common nor too rare and infection prevalence is either high or low. When social information and infection prevalence are both at intermediate levels, selection may favour increased or decreased sociality depending on the relative costs and benefits of acquiring social information and parasites, the relative timescales of the transmission processes, as well as the sociality of other members of the population. As a result, we show how ecological conditions can determine when: (1) the population evolves towards an evolutionarily stable level of sociality; (2) parasites are unable to constrain selection for sociality; or (3) when selection for reduced sociality drives social information and parasites ‘extinct’.

## Model description

### Full model

We model the (co)evolution of sociality (contact effort, *E*) and virulence (disease-associated mortality rate, *α*) due to the transmission of social information and horizontally-transmitted parasites that cause infectious disease (Fig. 1). We assume that the environment is constantly changing (e.g., food sites may be ephemeral, predation risk may vary over time and space) and that individuals can learn about the environment independently and through social cues. For simplicity, individuals either have poor (*P*) or good (*G*) information about the environment. Those with poor information have a baseline mortality rate of *d* (on average), and those with good information have mortality rate *ad*, where 0 ≤ *a* < 1. All individuals continuously learn about the environment at a constant (arbitrary) rate, but the extent to which an individual can learn independently is assumed to be limited, corresponding to a relatively poor information state (*P*). An individual in a poor information state can only improve to a good information state (*G*) through social cues from an individual who is better informed about the environment, which they do with probability *τ* given social contact. Individuals are poor at integrating information, such that multiple individuals with poor information cannot come together to produce good information–good information can only be socially learnt. For example, the location of a highly camouflaged predator may (almost never) be individually discovered, but is readily socially learnt, and that information is only valid while the individual is foraging near the predator. Since the environment is constantly changing, and since individual priorities may change (e.g., an individual may seek shelter once satiated, seek a different type of food, or simply leave the area), socially learned information loses value to an individual over time (but may still be valuable to others). For simplicity, individuals in the good information state revert to the poor state of information at a constant rate, *σ*.

**Figure 1.**
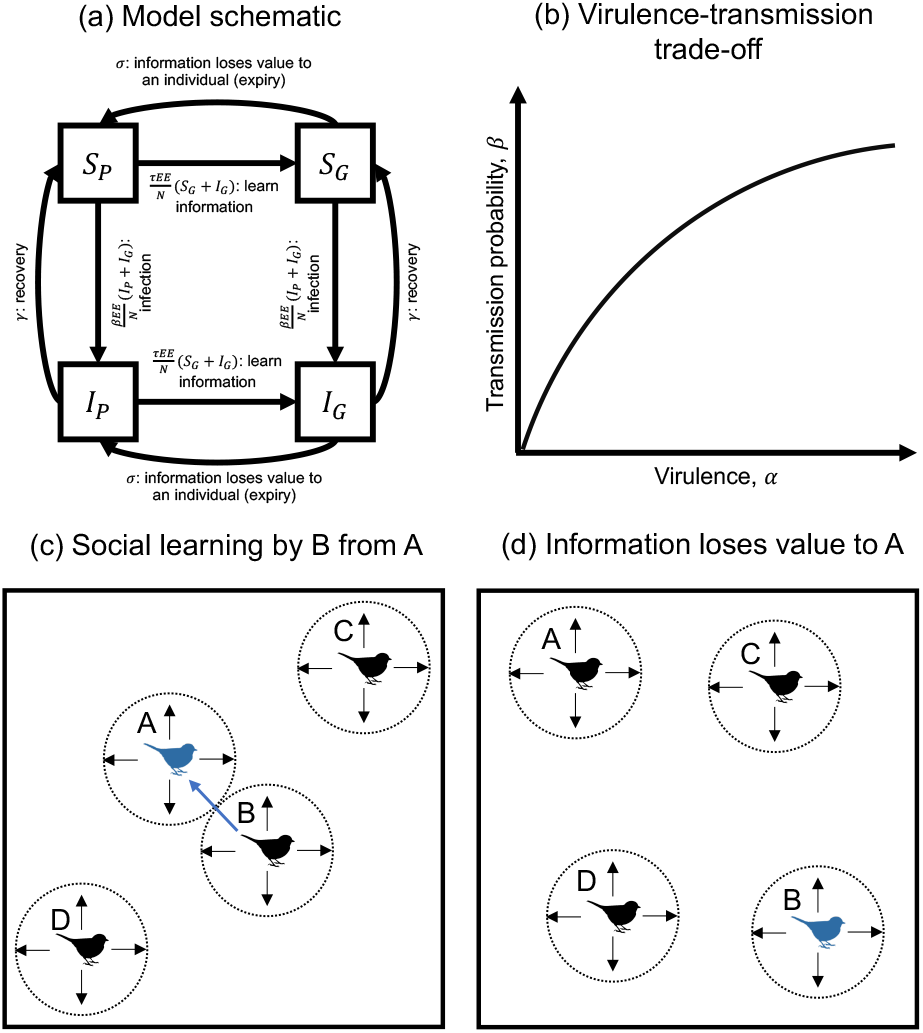
Model schematic and illustration of social information dynamics. (a) Transition diagram with percapita rates of transition between states, as described in the main text. (b) Illustration of the virulence-transmission trade-off (*β*’(*α*) > 0, *β*”(*α*) < 0). (c-d) Illustration of social information dynamics. (c) An individual, B, in the “poor” information state (black) learns socially transmitted information about the environment from individual A, who is better informed (i.e., in the “good” information state, blue). (d) Information loses value to individual A (expires), but remains valuable to individual B (e.g., individual A may be satiated or may leave the area).

In addition to their information status, individuals are classed as either susceptible (*S*) or infectious (*I*). Infected hosts transmit parasites to susceptible social contacts with probability *β* per social contact. Infected hosts recover at rate *γ*, at which point they become susceptible again. Infection leads to an increase in the host mortality rate due to virulence, *α*. We assume that the transmission probability (*β)* and virulence (*α*) are evolving traits subject to a trade-off such that *β* = *β*(*α*). While increased transmission is beneficial for the parasite, increased virulence is costly as it results in higher host mortality and therefore reduces the average infectious period. In the absence of a trade-off, the parasite would therefore maximise the infectious period by minimising virulence. We assume that new transmission stages are produced by infecting host cells or tissues, and so more transmissible parasites are also more virulent (*β*’(*α*) > 0). We also assume that the parasite eventually tends towards perfect transmission as virulence increases (*β*(*α*) → 1 as *α* → ∞, with *β*”(*α*) < 0; Fig. 1b). This means that there are diminishing returns on virulence, which ensures that there is always an optimal level of virulence, *α*\*.

We model social contacts following Bonds et al. (2005). Social contacts are assumed to be ephemeral, random, and occur based on contact ‘effort’. Each individual, *i*, has a contact effort, *E_i_*, which is assumed to be a heritable quantitative trait. An individual’s contact frequency, 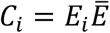, is the product of its own contact effort and the average contact effort of the population 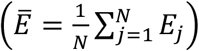. We assume that the only benefits and costs associated with contact effort (sociality) are access to survival-related information and the risk of infection, respectively.

In a monomorphic population with *E_i_* = *E* there are four classes (*S_P_,S_G_, I_P_, I_G_*), where *S_i_* and *I_i_* refer to susceptible and infected individuals in information state *i* ∈ {*P, G*} and the total population size is *N* = *S_P_* + *S_G_* + *I_P_* + *I_G_*. The per-capita birth rate is equal to (*b* – *qN*), where *b* is the maximum birth rate and *q* is the strength of density-dependence (resource competition). The per-capita information transmission rate is equal to the product of the information transmission probability per contact, *τ*, the population contact frequency, *E*^2^, and the proportion of individuals in state 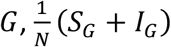. Similarly, the parasite transmission rate is equal to the product of the parasite transmission probability per contact *β*, the population contact frequency, *E*^2^, and the proportion of individuals who are infected, 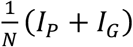. The population dynamics are described by the following set of ordinary differential equations (ODEs):

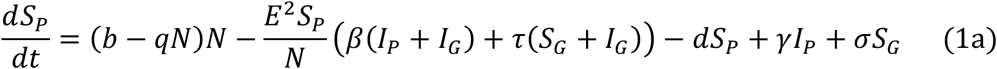

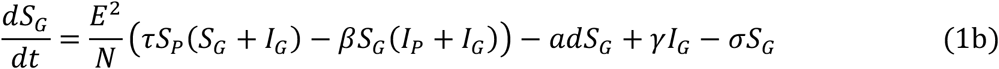

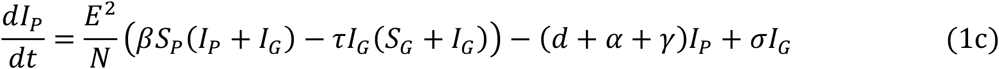

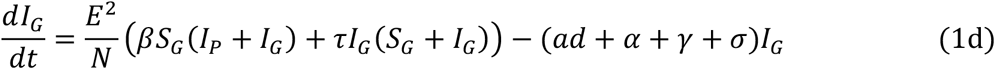

### Fast social information dynamics approximation

When the dynamics of social information are much faster than demographic (*τE*^2^, *σ* ≫ *b, d*) and epidemiological processes (*τE*^2^ ≫ *β, σ* ≫ *γ*), the proportion of individuals in the good information state for a given value of *E, G_E_*, rapidly reaches equilibrium. Letting 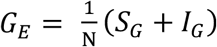, we see that:

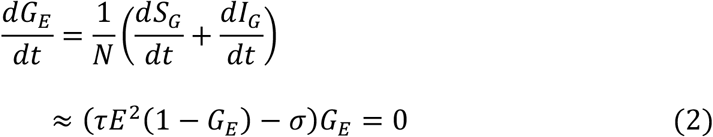

Thus, either no individuals are in information state *G*, or 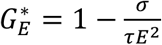 when *σ* < *τE*^2^. We can therefore make the following approximation to the full model when social information dynamics are relatively fast:

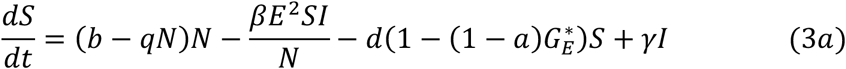

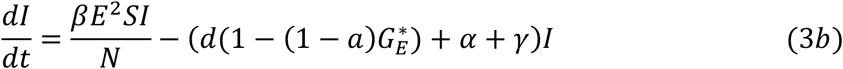

where *N* = *S* + *I* and 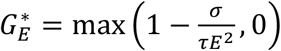 is the proportion of individuals in information state *G*. Note that 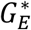 is an increasing, concave downward function of *E*, which means that there are diminishing effects of sociality on mortality.

### Analysis

We use evolutionary invasion analysis (adaptive dynamics; Geritz et al. 1998) to determine how host contact effort (*E*) and parasite virulence (*α*) (co)evolve. Briefly, this method assumes that mutations (1) are sufficiently rare, so that we can separate ecological and evolutionary timescales (i.e. the ecological dynamics reach equilibrium before the next mutation occurs); and (2) have small phenotypic effects, so that mutant traits are similar to resident traits. For the full model (equation 1), we derive host fitness analytically (see *Supplementary Material* for details), but solve the system numerically, as it is not possible to find an expression for the coexistence equilibrium where parasites and social information are both present. For the fast social information approximation (equation 3), it is possible to find an expression for the coexistence equilibrium, and in turn, expressions for fitness and fitness gradients. However, we omit these expressions as they are lengthy and provide no analytical insights, and so we present numerical results instead. We illustrate the different dynamics using evolutionary simulations. Briefly, every *T* time units of the deterministic ODE solver, a rare mutant is introduced which is phenotypically similar to a randomly chosen member of the population. Phenotypes which fall below an arbitrary frequency, *ϵ*, are removed from the population, and the process is repeated. The simulations relax the adaptive dynamics assumptions of continuous traits and a complete separation of ecological and evolutionary time scales (source code available in the *Supplementary Material* and at github.com/ecoevogroup/Ashby_and_Farine_2020).

### Ecological dynamics

In the full model, the host population has a trivial equilibrium at 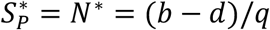 in the absence of parasites or social information, which exists provided *b* > *d* (hereafter we assume this condition is always satisfied). This equilibrium is stable if:

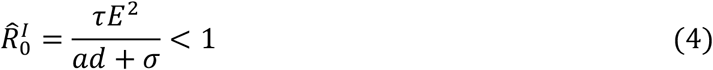

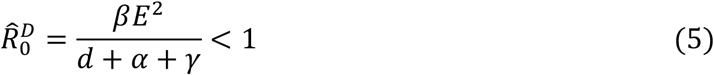

where 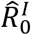 and 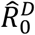 are the basic reproductive ratios for social information and infection at the trivial equilibrium, respectively. The basic reproductive ratios are the average number of transmission events per informed or infected individual in an otherwise uninformed or susceptible population. If 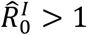 then social information can spread in the absence of parasites and if 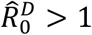 then infection can spread in the absence of social information.

The full model also has three non-trivial equilibria: (i) information-only (viable if 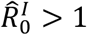), (ii) infection-only (viable if 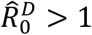), and (iii) both information and infection present (coexistence equilibrium). The latter is viable if 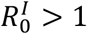 at the infection-only equilibrium and 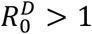 at the information-only equilibrium, where 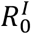 and 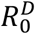 are the full basic reproductive ratios for information and infection:

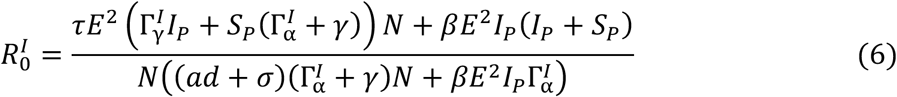

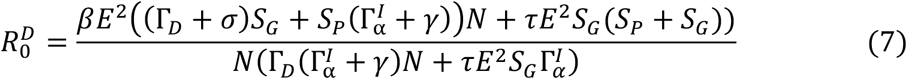

with 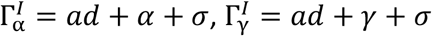 and Γ_*D*_ = *d* + *α* + *γ* (see *Supplementary Material* for derivation). Note that in the special case where there is either no infection or no social information, equations 6-7 reduce to equations 4-5. Intuitively, for the coexistence equilibrium to be stable, we require 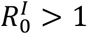 at the infection-only equilibrium and 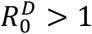 at the information-only equilibrium (i.e. social information and infection can always spread when rare). While it is not possible to obtain an analytical expression for the coexistence equilibrium, a numerical parameter sweep indicates that it is always stable when it is viable.

The fast social information approximation has two equilibria. There is an infection-free equilibrium at 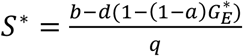 and a coexistence equilibrium at:

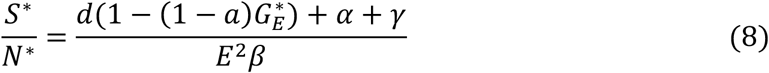

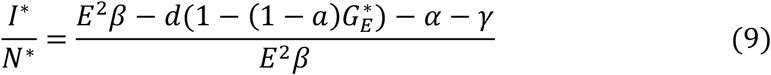

where

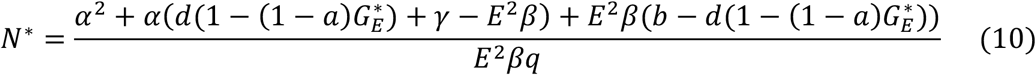

The corresponding basic reproductive ratios for the fast social information approximation are:

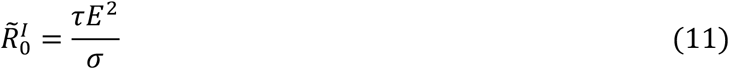

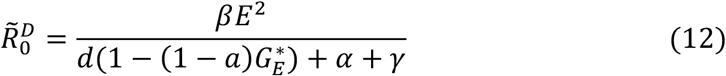

(see *Supplementary Material*). The coexistence equilibrium is stable when 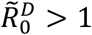 and is unstable when 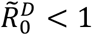 (the opposite is true for the infection-free equilibrium). Note that when 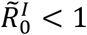 we must have 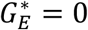 (social information transmission cannot be sustained), and hence 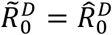.

The basic reproductive ratios for the full model and the approximation are very similar provided social information dynamics are sufficiently fast (*τ* ≫ *β*; Fig. 2a-b). When information dynamics occur on slower timescales, then the approximation diverges from the full model (Fig. 2a-b). Likewise, as the information dynamics become increasingly fast relative to the epidemiological dynamics, the equilibrium population distributions in the full model approach that of the approximation (Fig. 2c-d).

**Figure 2.**
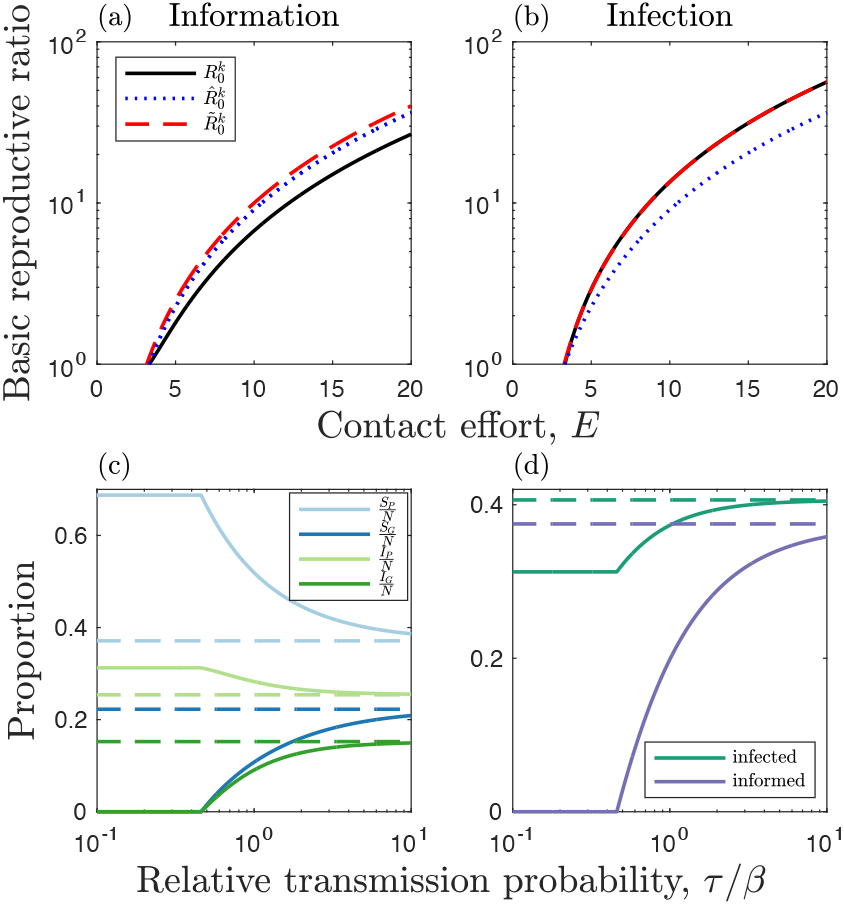
Ecological dynamics. (a)-(b) Basic reproductive ratios for (a) social information (*k* = *I*) and (b) infection (*k* = *D*) as a function of the contact effort, *E*. Curves correspond to the basic reproductive ratio in the full model, 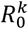 (black, solid), the full model without infection or social information present, 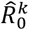 (blue, dotted), and the approximation for fast social information dynamics, 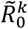 (red, dashed). (c)-(d) Comparison of the non-trivial equilibria in the full model (solid) and the approximation for fast social information (dashed). All parameters except for *τ* and *σ* are fixed, with *σ/τ* = 10 held constant. The full model approaches the approximation when social information dynamics are fast relative to other processes. (c) Proportion of individuals in each class. (d) Proportion of individuals that are infected (green) or are in the good information state (purple). Fixed parameters (where applicable): *a* = 0.2, *b* = 1, *d* = 0.5, *E* = 4, *q* = 10^-3^, *α* = 0.4, *β* = 0.1, *γ* = 0.2, *σ* = 1, *τ* = 0.1.

### Evolution of sociality

We explore the evolution of host sociality in terms of the contact effort, *E*. We first investigate how the fitness gradient varies for a fixed value of *E* to illustrate how social information and infection prevalence influence the direction of selection. We then focus our analysis on the extent to which social information reduces mortality (*a*) (note that smaller values of *a* correspond to greater benefits of social information), the cost of being infected (*α*), the rate at which social information loses value to an individual (expires) relative to its transmission probability (*σ/τ*), and the relative transmission probabilities of social information and infection (*τ/β*).

To explore host evolution, we consider the invasion of rare host mutant with contact effort *E_m_* ≈ *E* into an established resident population at equilibrium (denoted by asterisks). In the full model, the invasion dynamics of the rare host mutant are given by:

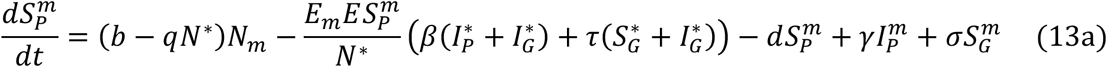

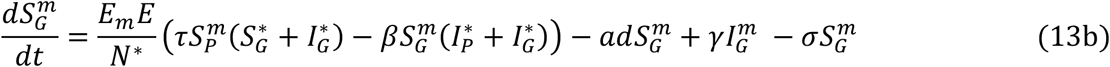

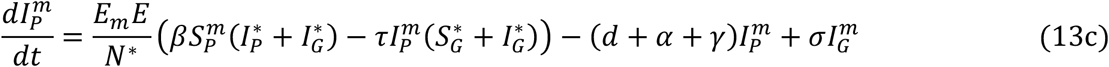

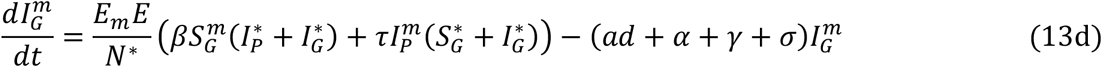

For the fast social information approximation, we let 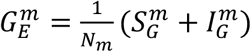 so that:

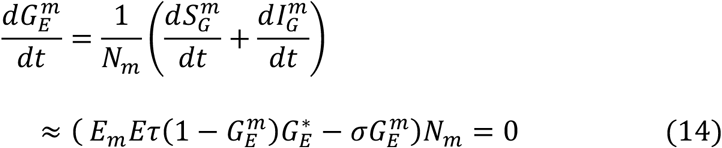

Rearranging and substituting 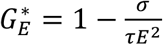 gives:

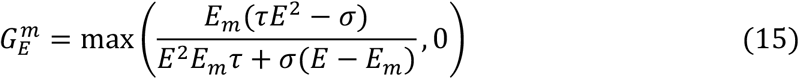

We can therefore approximate the invasion dynamics of a rare mutant when social information dynamics are fast by:

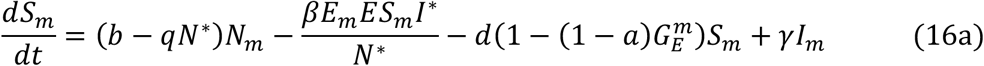

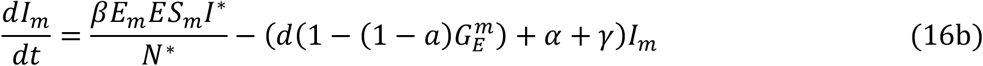

In the *Supplementary Material*, we show the derivation for invasion fitness in both versions of the model.

### Host fitness gradient

Fig. 3 shows how the host fitness gradient varies with social information and infection prevalence when the benefits of sociality are relatively high (Fig. 3a) or low (Fig. 3b) compared to the costs. Social information dynamics are assumed to be fast, but the overall patterns hold for slow information dynamics. Note that here *E* is fixed, with *σ* and *β* modulating social information and infection prevalence, but when *E* evolves (see *Host evolutionary dynamics*) it will feedback to affect prevalence and hence selection for sociality. Nevertheless, considering how the fitness gradient varies when *E* is fixed allows us to qualitatively see how changes in social information and infection prevalence affect the fitness gradient, all else being equal.

**Figure 3.**
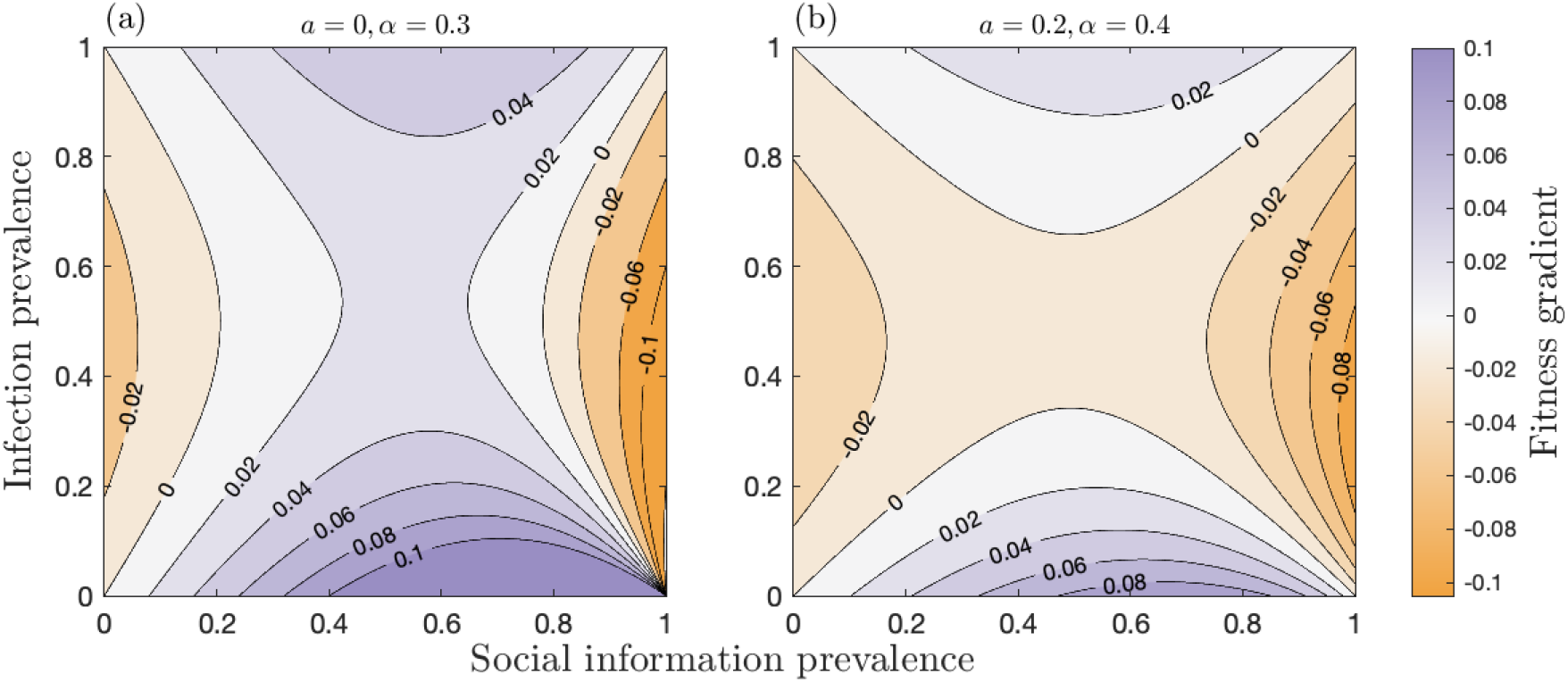
Effects of social information and infection prevalence on the host fitness gradient. Contour plots of the host fitness gradient, in the fast social information approximation as a function of social information and infection prevalence with: (a) higher (*a* = 0, *α* = 0.3) and (b) lower (*a* = 0.2, *α* = 0.4) benefits of sociality relative to costs. All parameters except for *σ* and *β* are fixed within each panel, with values of *σ* and *β* chosen so that social information and infection prevalence vary accordingly. In both cases, the host fitness gradient is positive (bluer shading) when social information prevalence is intermediate and there is either a high or low level of infection, and is negative (redder shading) when social information prevalence is either at a high or low level and infection is at intermediate prevalence. Fixed parameters as in Fig. 2, except *β* = 0.2.

The fitness gradient is always positive when social information prevalence is intermediate and infection is either rare or common, regardless of whether the benefits of sociality are relatively high or low compared to the costs. Intuitively, increased sociality is likely to be beneficial when infection prevalence is low, but only if social information is neither too rare nor very common. This is because greater sociality does not significantly increase the rate at which individuals obtain social information if it is readily available or unlikely to become accessible. Somewhat counter-intuitively, increased sociality may also be beneficial when infection prevalence is high (as shown in Bonds *et al*., 2005). This is because most individuals are likely to be infected anyway, but increased sociality will improve access to social information with little increase to the risk of infection. Conversely, the fitness gradient is always negative when social information is either rare or very common and infection is at intermediate prevalence. This pattern mirrors the one described above, but with the roles of social information and infection switched due to their opposing effects on the fitness gradient. Intuitively the fitness gradient is negative when social information prevalence is low and infection is common in the population, as greater sociality increases the risk of infection but does not substantially increase the likelihood of obtaining social information. But perhaps counter-intuitively the fitness gradient is also negative when social information is widespread and infection is at moderate levels. This is because most individuals are well-informed, and so increased sociality will only marginally improve access to social information but may substantially increase the risk of infection.

When both social information and infection are at intermediate prevalence, the sign of the fitness gradient depends on the relative costs and benefits of being infected or informed. When awareness of social information is relatively more valuable than the cost of being infected, the fitness gradient is positive whenever social information prevalence is at an intermediate level regardless of infection prevalence (Fig. 3a). Infection may therefore not always be sufficient to curtail the evolution of sociality. In contrast, when awareness of social information is less valuable, and the cost of being infected is relatively high, the fitness gradient is negative whenever infection prevalence is at an intermediate level regardless of social information prevalence (Fig. 3b). Our qualitative analysis of the fitness gradient suggests that the host population may experience different evolutionary outcomes depending on the initial conditions: if infection prevalence is sufficiently low the population may evolve towards a stable level of sociality, but if infection is initially very common then the population may experience runaway selection for greater sociality.

### Host evolutionary dynamics

We focus on how the host contact effort, *E*, evolves depending on the rate at which social information loses value to an individual (the expiry rate), *σ*, relative to the information transmission probability, *τ*, which we consider for fixed costs of infection (virulence, *α*) and benefits of social information (*a*) (Fig. 4, S1). The shaded regions in Fig. 4 and S1 indicate the direction of selection for the contact effort. If the initial contact effort is too low, then neither infection nor social information is viable (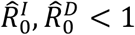; grey regions in Fig. 4, S1) and so no adaptive evolution occurs. If 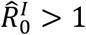 and/or 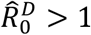, adaptive evolution will occur, causing the contact effort to increase or decrease until the contact effort either: (i) evolves to a *continuously stable strategy* (CSS; an ‘evolutionary endpoint’); (ii) decreases until neither infection nor social information is viable 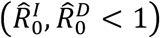, resulting in a semi-stable contact effort (selection prevents an increase in contact effort, but drift may cause it to decrease); or (iii) increases until a physiological limit is reached. The first outcome (CSS) generally occurs for sufficiently low values of *σ/τ* (social information expires at a relative slow rate, all else being equal), provided the initial contact effort is neither too low nor too high. This ensures that social information prevalence is neither too rare nor too common, resulting in evolution towards an optimal contact effort (Fig. 5a-b). If it exists, the CSS peaks for intermediate values of *σ/τ*, which will typically correspond with moderate levels of information prevalence. Social information expires more quickly as *σ/τ* increases, and so the overall prevalence of social information in the population falls, which increases selection for sociality. However, as sociality increases so too does infection prevalence, which reduces selection for sociality.

**Figure 4.**
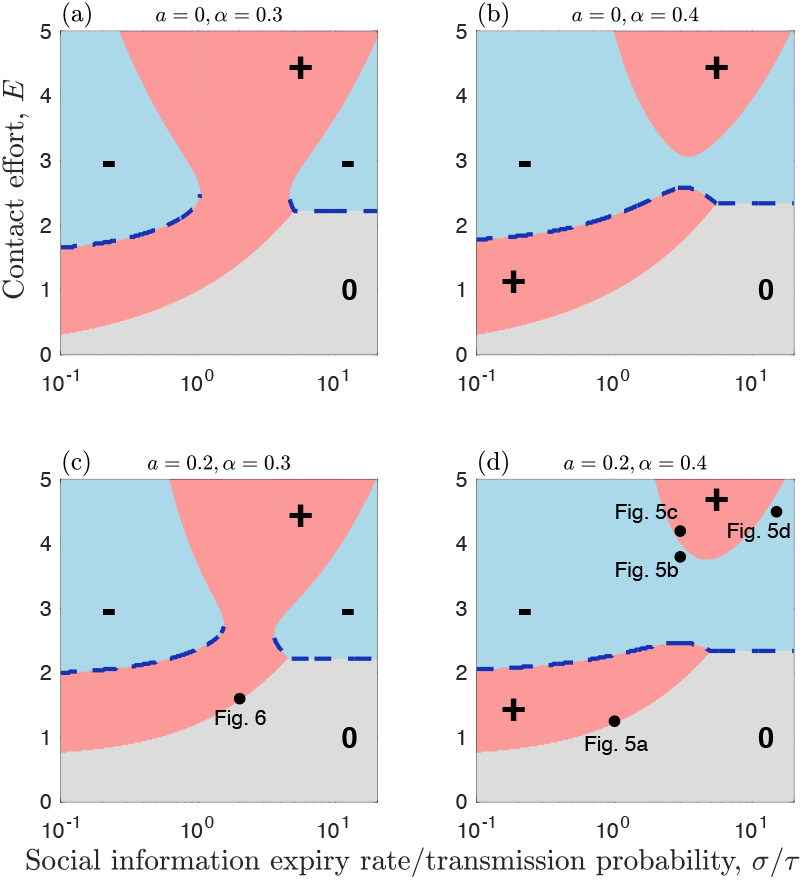
The evolution of sociality arising from social information and parasite transmission. Evolution of contact effort, *E*, as a function of the social information expiry rate divided by the transmission probability, *σ/τ*, and for different values of the benefits of social information, *a*, and virulence, *α*. Note that smaller values of *a* correspond to greater benefits of social information. Results are shown for relatively slow social information dynamics, but are broadly similar for fast information dynamics (Fig. S1). Pink (+) and blue (-) regions indicate when the fitness gradient is positive (*E* will increase) and negative (*E* will decrease), respectively. Grey (0) regions indicate when there is no selection as social information and infection are both absent from the population. Blue dashed curves indicate stable or semi-stable levels of contact effort. Dots in (d) and (e) correspond to initial conditions for simulations in Fig. 6 and 5, respectively. Fixed parameters as in Fig. 2, except *β* = 0.2.

**Figure 5.**
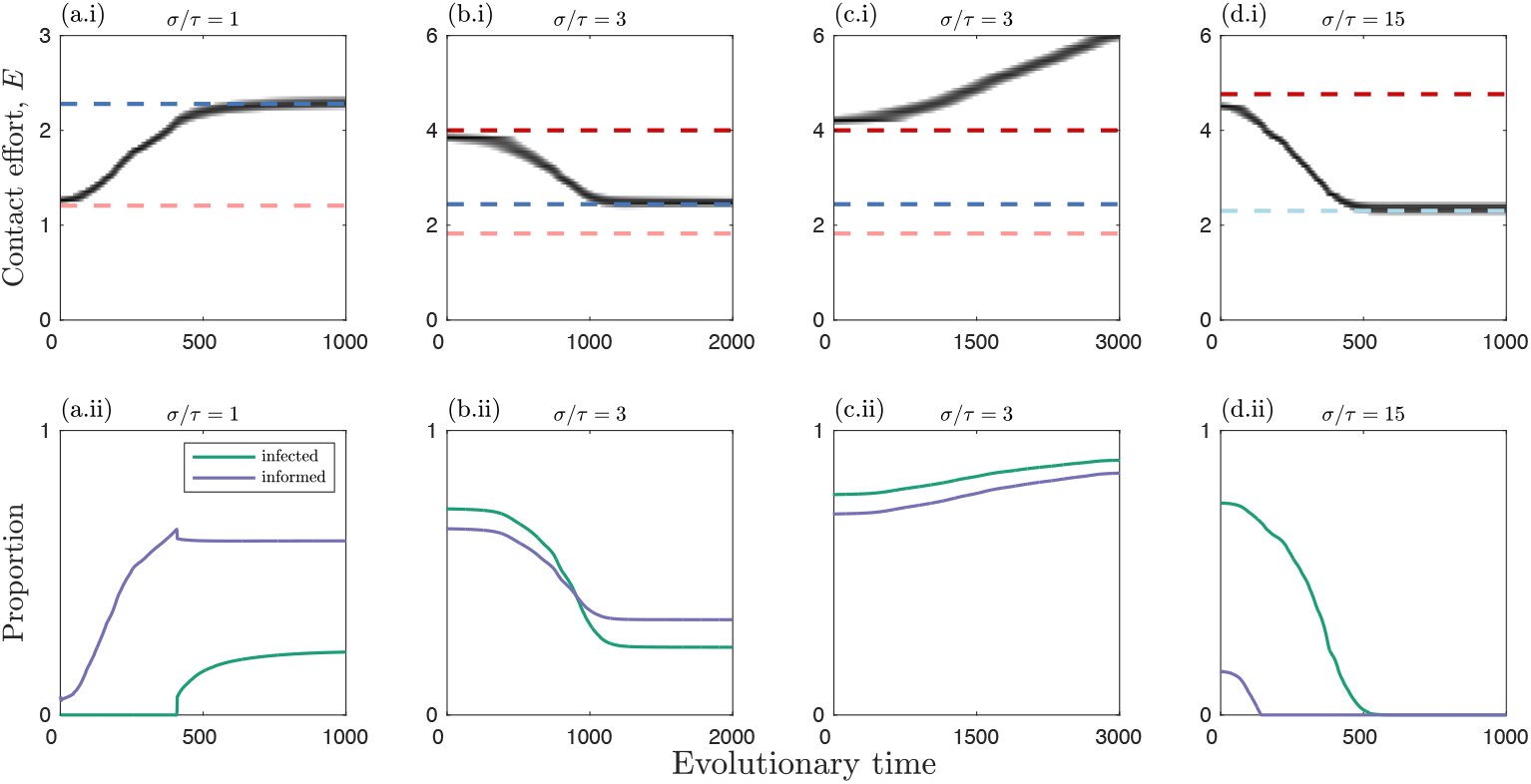
Simulation trajectories when social information and epidemiological dynamics occur on similar timescales. Trajectories of simulations corresponding to crosses in Fig. 4e. Top row: evolutionary trajectories (solid black); singular strategies: repellers (red) and continuously stable strategies (CSSs; dark blue); and thresholds below which social information and information transmission is viable (pink and light blue, respectively). Above the pink dashed line, sociality increases, and above the light blue line sociality decreases. Bottom row: proportion of individuals that are infected (green) or are in the good information state (purple) in each simulation. Fixed parameters as in Fig. 2, except *τ* = *β* = 0.2.

The second outcome (semi-stable contact effort) occurs for sufficiently high values of *σ/τ* (social information expires at a relative fast rate, all else being equal). As information is relatively scarce, this causes the population to evolve a lower contact effort until both 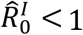 and 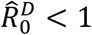, at which point the fitness gradient is zero and no further adaptive evolution occurs (hence it is semi-stable, as it could still be subject to drift; Fig. 5d). For intermediate values of *σ/τ*, the contact effort may evolve to a stable evolutionary endpoint (CSS) or increase through runaway selection until a physiological limit is reached (Fig. 5c). The outcome depends on the precise costs of infection (*α*), the benefits if information (*a*), and the initial contact effort. Higher costs of infection (larger *α*) and smaller benefits of information (lower *a*) generally have a stabilising effect, whereas lower costs of infection and larger benefits of information promote runaway selection at intermediate *σ/τ*. Thus, if the costs of being more social are relatively low, the population may experience runaway selection, but at higher costs, sociality is curtailed at a stable intermediate level. Moreover, the higher the initial contact effort, the more likely the population will experience runaway selection. Runaway selection can occur, for example, when most individuals in the population are infected, and so there is little cost to increased sociality. Runaway selection for sociality is unrealistic as contact efforts would be constrained by other factors (e.g. other costs or physical constraints). Instead, runaway selection for sociality in the model should be interpreted as infection being insufficient to constrain sociality.

The results are broadly the same whether social information and epidemiological dynamics occur on similar timescales (*σ* ≈ *γ*; Fig. 4) or when the former occurs on a much faster timescale (*σ* ≫ *γ*; Fig. S1). However, there are a few notable differences. First, the viability threshold for social information transmission is lower when the dynamics are rapid 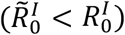, which means the population can evolve over a broader range of initial contact efforts. Second, faster information dynamics generally make runaway selection less likely. This means that when social information dynamics are relatively slow, there may be runaway selection for greater sociality, but when the social information dynamics are relatively fast, sociality may evolve to a stable level (Fig. 6). The timescale of social information dynamics can therefore play an important role in determining the evolution of sociality.

**Figure 6.**
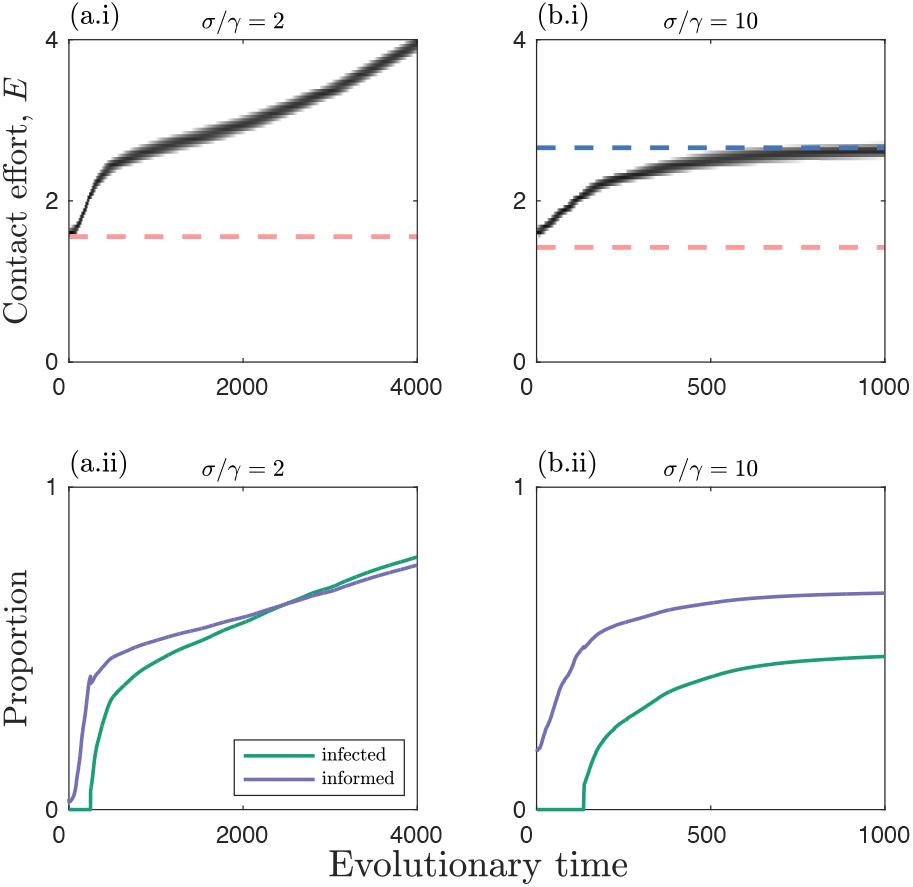
Simulation trajectories when social information dynamics occurs on similar (a) or faster (b) timescales than epidemiological dynamics. Simulations corresponding to the asterisk in Fig. 4d. Top row: evolutionary trajectories (solid black); continuously stable strategies (CSSs; dark blue); and viability threshold for social information (pink). Bottom row: proportion of individuals that are infected (green) or informed by social information (purple) in each simulation. Fixed parameters as in Fig. 2, except *α* = 0.3, *β* = 0.2, *σ/τ* = 2.

### Evolution of virulence

To explore parasite evolution, we consider the invasion of rare mutant with virulence *α_M_* ≈ *α* into an established resident population (at equilibrium, denoted by asterisks). For simplicity we assume that social information dynamics are fast by using the approximation to the full model. The invasion dynamics of the rare parasite mutant are given by:

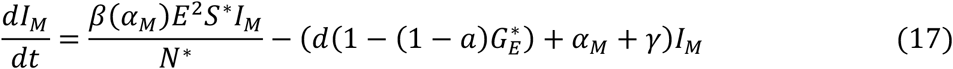

In the *Supplementary Material*, we derive the invasion fitness and show that evolution maximises the basic reproductive ratio, 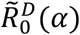:

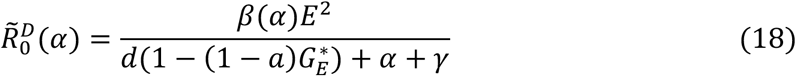

We now consider how the evolution of sociality affects the evolution of parasite virulence.

The optimal level of virulence, *α* *, occurs when:

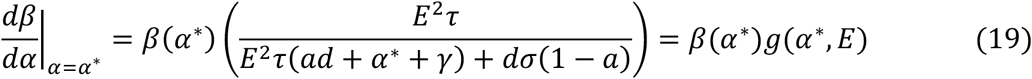

where *g*(*α*\*, *E*) is used for notational convenience and is equal to the term in parentheses. To determine what happens to the optimal level of virulence when sociality changes, consider 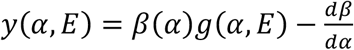 and suppose *α* = *α*\*, so that *y*(*α*\*, *E*) = 0. Note that *g*(*α, E*) is a decreasing function of *α* and is an increasing function of *E* provided 0 ≤ *a* < 1. Hence, *y*(*α, E*) is also an increasing function of *E*. Now consider the derivative of *y* with respect to *α* at *α* = *α*\*:

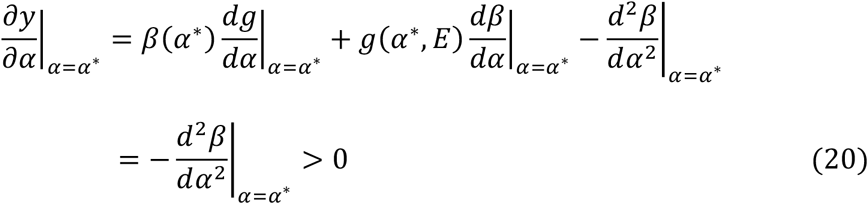

where 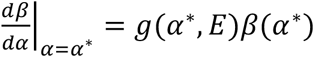 from equation 19 and 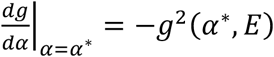. This means that if sociality increases, the optimal level of virulence must decrease (Fig. 7). This may appear to contradict classical predictions on population mixing and the evolution of virulence (Ewald 1994), but the difference occurs due to the effects of social information on the mortality rate of the host, which affects the infectious period. Note that if the host mortality rate is independent of sociality (*a* = 1), then it follows that the optimal level of virulence occurs when:

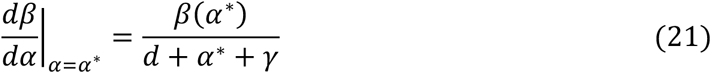

which does not depend on the contact effort, *E*. Thus, the reason optimal virulence is inversely related to sociality in the present model is due to effects of sociality on the mortality rate of the host (and hence the infectious period).

**Figure 7.**
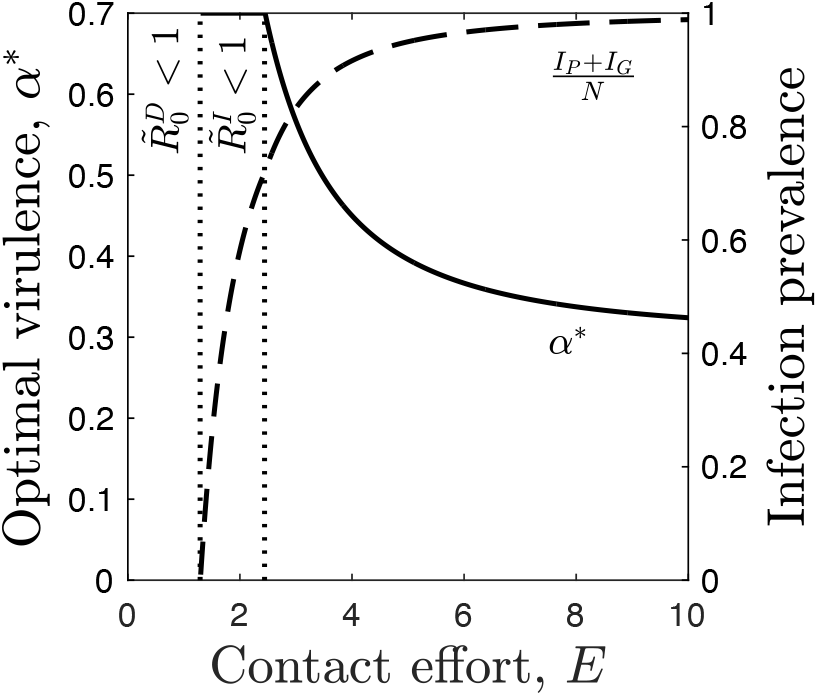
The evolution of virulence in the fast social information approximation. Optimal virulence, *α*\*, decreases with the contact effort (solid) and infection prevalence (dashed) increases. Vertical dotted lines indicate where parasite 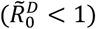 and 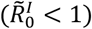 social information transmission are not viable. Other parameters as in Fig. 2, with 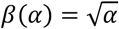 and 0 ≤ *α* ≤ 1.

### Coevolution of sociality and virulence

We combine our results from the previous sections to analyse the effects of social information on the coevolution of sociality and virulence. For tractability, we make three simplifying assumptions: (1) social information dynamics are fast (i.e. we use the approximation to the full model); (2) the parasite evolves much faster than the host; and (3) the relationship between transmission and virulence is 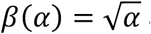 and 0 ≤ *α* ≤ 1. Simulations indicate that relaxing these assumptions does not qualitatively affect our results. The second assumption allows us to substitute *α* = *α*\* into the host fitness gradient, where:

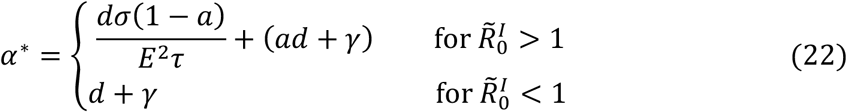

The range of qualitative outcomes for coevolution are the same as for one-sided adaptation in the host (Fig. 8). The host and parasite may tend to a joint stable level of sociality and virulence, respectively, or they may experience runaway selection for higher sociality and lower virulence, or infection or social information may be unviable. However, note that in the case of runaway selection virulence asymptotes to a lower bound of *α*\* = *ad* + *γ* (from equation 22). Under coevolution, the evolutionary stable level of virulence increases with the ratio of the information expiry rate to transmissibility (*σ/τ*), until social information dynamics are no longer viable 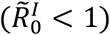.

**Figure 8.**
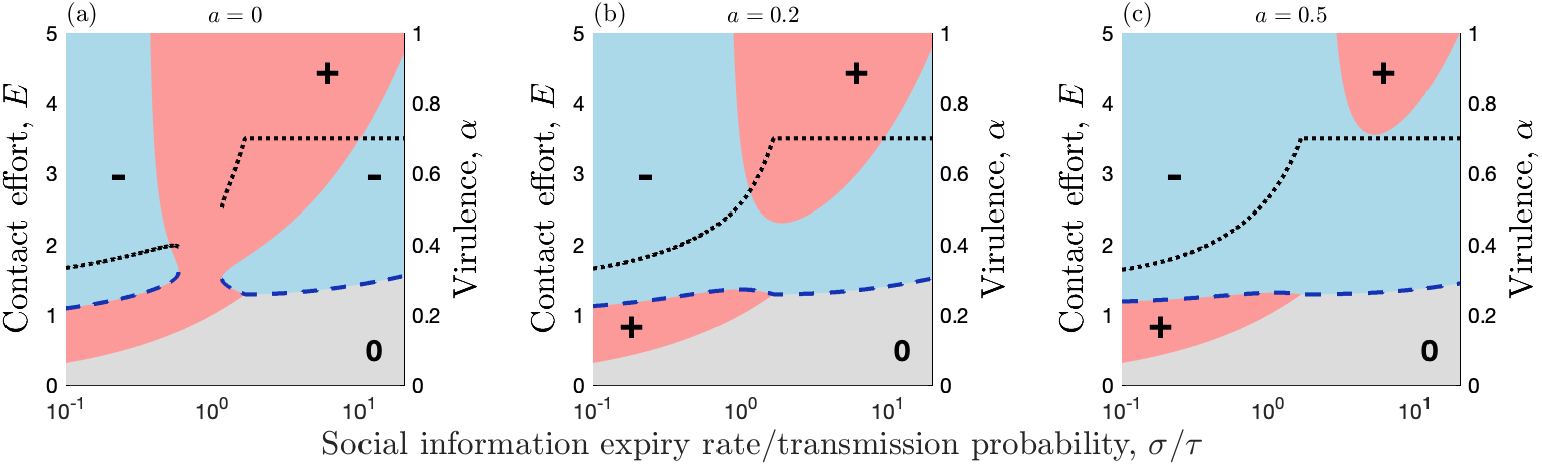
Coevolution arising from social information and parasite transmission dynamics. Coevolution of host sociality (contact effort, *E*) and parasite virulence (*α*) as a function of the social information expiry rate divided by the transmission probability, *σ/τ*, and for different values of the benefits of social information, *a* (fast social information approximation). Note that smaller values of *a* correspond to greater benefits of social information. Pink (+) and blue (-) regions indicate when the fitness gradient is positive (*E* will increase) and negative (*E* will decrease), respectively. Grey (0) regions indicate when there is no selection as social information and information are both absent from the population. Blue dashed curves indicate stable or semistable levels of contact effort and black dotted curves indicate the corresponding stable levels of virulence. Other parameters as in Fig. 2, with *τ* = 1, 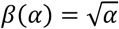 and 0 ≤ *α* ≤ 1.

**Table 1 –.** Summary of key results.

| Scenario | Variable | Description of results | Figure(s) |
| --- | --- | --- | --- |
| Host evolution | Infection prevalence | Selection against contact effort at high/low infection prevalence | Fig. 3 |
|  | Social information prevalence | Selection for higher contact effort at high/low social information prevalence | Fig. 3 |
|  | Social information expiry rate | Slow expiry: contact effort either evolves to a stable level or increases without bound; fast expiry: contact effort either increases without bound or decreases until parasites are driven extinct | Fig. 4 |
|  | Social information timescale | Faster social information dynamics generally have a stabilising effect (prevent runaway selection for contact effort) | Fig. 4, 6 |
| Parasite evolution | Contact effort | Virulence decreases with higher contact efforts, all else being equal | Fig. 7 |
| Coevolution | Social information expiry rate | Virulence increases with the social information expiry rate until social information is no longer viable | Fig. 8 |

## Discussion

Sociality entails both risks and benefits, but there is currently little understanding of how dynamic ecological processes influence the evolution of sociality. Here, we explored a model of social information and parasite transmission to determine their effects on the evolution of host sociality and parasite virulence. Although previous studies have examined the role of parasitism in the evolution of sociality, ours is the first that captures both dynamic costs and benefits of different social tendencies. This means that the fitness benefits of sociality, in terms of increased access to social information, and fitness costs, in terms of increased exposure to infection, are not imposed on the system, but instead are emergent properties that are subject to eco-evolutionary feedbacks. Hence, changes in sociality at the individual level affect the properties of transmission at the population level, and these population-level properties, in turn, affect both the costs and benefits of sociality at the level of the individual.

Our model shows how contrasting transmission processes can trade-off to mediate the evolution of sociality and virulence. Crucially, we have shown that selection for sociality (and hence virulence) depends non-monotonically on the prevalence of both social information and infection (Fig. 3). Specifically, selection usually favours greater sociality (and lower virulence) when social information is neither too common nor too rare and infection risk is either low or high. Conversely, selection against sociality (and for greater virulence) typically occurs when social information is either rare or very common and infection is at moderate prevalence. The fact that these two processes mirror each other is not surprising: the transmission processes are fundamentally the same in the model, but social information is beneficial and infection is costly.

Naively, one might assume that selection for sociality should increase with social information prevalence and decrease with infection prevalence. But when social information prevalence is high, there is little benefit to increasing contact effort because most individuals are already informed; similarly, when infection prevalence is high, a higher contact effort carries little extra cost because most individuals are already infected. Slightly higher or lower sociality therefore makes little difference to the rate of exposure to social information or infection when prevalence is high. The effects of infection prevalence on the evolution of sociality have previously been observed by Bonds et al. (2005), and occurs across a variety of behavioural and physiological defences against parasitism, including resistance (Haldane 1949), recovery (van Baalen 1998), mate choice (Ashby and Boots 2015) and sexual reproduction (Ashby and King 2015). However, the effects of social information prevalence on the evolution of sociality have not previously been documented.

The patterns described above for high/low prevalence are robust, but the picture is more complicated when social information and infection are both at intermediate prevalence. Here, it is possible for evolutionary trajectories to exist where the sign of the fitness gradient is invariant to infection or social information prevalence, leading to runaway selection for higher sociality/lower virulence or lower sociality/higher virulence (Fig. 4, 8). However, while our model allowed for runaway selection for sociality, in reality one would expect other factors to eventually constrain evolution (e.g., physiological constraints, additional costs). We omitted additional constraints from our model so that we could determine when infection alone is sufficient to constrain the evolution of sociality. The results for “runaway selection” should therefore be interpreted in this light.

In previous studies (Bonds et al. 2005; Prado et al. 2009), the fitness benefits of sociality were fixed rather than dynamic. While some modelling studies have explored how social information and infectious disease affect sociality, this has been in the form of host plasticity during an epidemic (Funk et al. 2010), rather than an evolutionary response, or where the outcomes are compared across populations with different fixed social strategies (Udiani and Fefferman 2020). Here, the two processes operate as independent forces of selection. Our findings are closely linked to those of Bonds et al. (2005), where sociality and virulence are shown to coevolve to stable levels. In contrast, Prado et al. (2009) found that coevolutionary cycling (fluctuating selection) may occur between sociality and virulence. It is not clear why Prado et al. (2009) found cycles, yet Bonds et al. (2005) and the present study have not, but part of the reason may be due to model structure. Prado et al. (2009) modelled epidemiological dynamics using a stochastic individual-based network, whereas Bonds et al. (2005) and the present study employed deterministic models with randomly mixing populations. A priority for future research in this area is therefore to establish precisely when coevolutionary cycling of sociality and virulence occurs, whether this is driven by population structure and/or stochasticity, and how it relates to other mechanisms that generate coevolutionary cycling (Ashby and Boots 2017).

An important consideration when modelling the benefits of sociality dynamically in terms of social information, as opposed to fixed benefits (Bonds et al. 2005; Prado et al. 2009), is the timescale of information dynamics relative to other processes. We have shown that for the most part, evolutionary outcomes are qualitatively and quantitatively similar whether social information dynamics are relatively fast or on a similar timescale to demographic and epidemiological processes. However, this is not always the case. In general, faster social information dynamics tend to have a stabilising effect (Fig. 6), reducing parameter ranges for runaway selection. Hence, while one can often approximate social information dynamics by assuming the population rapidly reaches a stable distribution, this assumption does not always hold and can fundamentally change evolutionary outcomes.

Our model also suggests that the initial conditions can determine the long-term evolutionary outcome (stable level of sociality and virulence, or runaway selection for increased sociality and decreased virulence; Fig. 4, 8). The possibility of more than one outcome suggests that different populations might evolve contrasting social strategies for coping with parasitism, and that variable ecological conditions can lead to sudden shifts in sociality and virulence. How evolutionary outcomes play out in a given population could be affected by the structure of the social contacts. Our model assumes a well-mixed population, which can represent dynamics in one small patch. However, at a broader scale, it is now clear that almost all animal populations exhibit some structure (Cantor et al. 2021b), and that population structure can shape both transmission dynamics and parasite evolution (Eames and Keeling 2006; Lion and Boots 2010; Ashby and Gupta 2013).

Social structure can arise due to individual preferences for certain relationships (Alberts 2019) or because the environment induces limitations on who can come into contact with whom (He et al. 2019; Silk et al. 2019; Albery et al. 2021). For example, recent work has suggested that—all else being equal—some landscapes may be more predisposed to facilitating parasite transmission than other landscapes (He et al. 2021). Social organisation can also play a part. For example, cooperative breeding can limit host dispersal (Armansin et al. 2020), affecting transmission processes and, in turn, selection for sociality and virulence. Furthermore, different dispersal strategies may evolve depending on the infection status of the host (Iritani 2015). Some mammal (Grueter et al. 2020) and birds (Papageorgiou et al. 2019; Camerlenghi et al. 2022) also exhibit multiple levels of social connections - named multilevel societies - with social contacts being expressed within and across social units.

A number of studies have also shown that within-population variation in sociality can influence the spread of social information (Aplin et al. 2012; Allen et al. 2013; Farine et al. 2015a; Snijders et al. 2021). For example, the social network attributes of wild great tits (*Parus major*) can be used to predict the discovery rate of novel food patches (Aplin et al. 2012). As in previous theoretical studies (Bonds et al. 2005; Prado et al. 2009), we did not observe coexistence of different social strategies. However, several empirical studies have highlighted the potential importance of maintaining mixed strategies in social groups or local populations. For example, Aplin et al. (2014) suggested that variation in sociality could increase group-level efficiency in exploiting patches. Similarly, in ants, mixed colonies (in terms of aggressive type) have more efficient task allocation and are more successful at capturing prey (Modlmeier et al. 2012), with task specialisation also buffering colonies from the effects of parasite exposure (Scharf et al. 2012).

One potential reason for the lack of mixed social strategies in current theoretical studies is that they do not include assortative mixing (Farine 2014). Assortative mixing has been shown to facilitate the coexistence of multiple parasites (Eames and Keeling 2006), so it is possible that mixed social strategies could also evolve in the presence of non-random mixing - a process that could be either actively generated through social preferences or passively through exogenous drivers of social structure. It is also possible that mixed social strategies could evolve in fluctuating environments, especially if individuals value information differently (e.g. due to personality traits, Aplin et al. 2014), or if individuals vary in their propensity to form social relationships (e.g. due to developmental conditions, Boogert et al. 2014; Brandl et al. 2019). A challenge for future theoretical research is to establish whether selection arising from infection and information-use dynamics can drive the evolution of mixed social strategies. Our results suggest that a number of potential mechanisms (e.g. variation in the perceived value of information) could facilitate this process.

Our model, along with many simulation studies of dynamics on social networks, further assumes individuals do not vary their contact effort depending on their infection or information status. In reality, infected individuals may become less social when they are sick or may be avoided by those who are healthy (reviewed in Romano et al. 2020; Stockmaier et al. 2021), and an individual may become more or less social depending on their information state (Kulahci and Quinn 2019) or starvation risk (Gareta García et al. 2021). Future theoretical work should therefore consider the implications of plastic sociality for host-parasite coevolution and how this may affect our predictions for how variance in information awareness affects selection for sociality.

One of the key predictions from this study and previous work (Bonds et al. 2005) is that sociality should vary inversely with mortality virulence, which may appear at odds with classical predictions for virulence evolution (Ewald 1994). The contradiction is resolved by noticing that: (1) our model considers contact rates in a randomly mixing population rather than contact number in a structured population; and (2) higher contact rates do not directly select for lower virulence, but do so indirectly through the spread of social information, which reduces host mortality and increases the infectious period. Hence, greater sociality as modelled here is not equivalent to population mixing in a structured model, and the effects on virulence evolution are mediated via host mortality rather than contact rates. Based on these insights, we therefore predict that sociality should covary inversely with mortality virulence in populations where social or spatial structuring is weak but should covary positively with mortality virulence when structuring is strong. Future empirical studies may be able to test this prediction directly by manipulating population structure or through comparative data (e.g. Sah et al. 2018).

While social structure, variation in social strategies, and social contact dynamics might be important, studies simulating disease transmission suggest that the importance of social structure in shaping parasite transmission is likely to be over-stated (Sah et al. 2017). Recent studies of how information is accumulated and integrated into cultural traits in structured populations have also suggested that transmission properties, rather than contact structure, have the most significant impact on cultural evolution in networks (Cantor et al. 2021a). For example, individuals can use different learning rules, such as conformist transmission [e.g. having a disproportionate preference for copying a more common behaviour (Aplin et al. 2015a)] or require passing a certain threshold number of informed contacts before becoming informed (Rosenthal et al. 2015). While real transmission processes for social information will rarely mirror those of parasites, such findings are highly relevant to the spread of infectious diseases (Sah et al. 2017; Evans et al. 2020). For example complex parasitic life cycles can impact how social contacts translate to transmission events (Grear et al. 2013; Farine 2017). Transmission properties will therefore modify how transmission pathways—for both disease and information—emerge from social contacts.

Fully testing the predictions of our model will require tracking infection and information states. Empirically determining the prevalence of infection in a population is generally straightforward, but it is likely to be more challenging to determine the prevalence of information. In some systems, it may be possible to focus on variance in information awareness or infection status. Our model predicts that when most individuals are either informed or uninformed (low variance), selection is unlikely to favour increased sociality, but when the numbers of informed and uninformed individuals are more balanced (high variance), selection may favour increased sociality (and in turn, lower virulence). Similarly, one can reframe the predictions based on infection prevalence in terms of variance in infection status.

The dynamic benefit of sociality in our model was motivated by studies of social information in animals, but our model is also applicable to other dynamic benefits such as symbiont transmission. If symbionts are horizontally transmitted and provide a benefit to their hosts in terms of reduced mortality (Brownlie and Johnson 2009; Ford and King 2016; Ashby and King 2017; Rafaluk-mohr et al. 2018), then we should expect our key results to hold. From an empirical perspective, it may be easier to test our predictions about dynamic costs and benefits using experimental evolution of bacteria and plasmids. Plasmids can increase or decrease bacterial fitness and can be transmitted through conjugation, with bacteria able to evolve higher or lower conjugation rates (akin to contact effort) depending on environmental conditions (Harrison and Brockhurst 2012). Bacteria-plasmid systems may therefore prove to be a more tractable target for directly testing some of our predictions, especially those involving coevolution.

In summary, our study suggests that not only does the sociality of the population affect social information and epidemiological dynamics, but also that these processes are likely to influence the evolution of sociality and virulence. Crucially, we predict that: (1) selection for sociality and virulence will vary non-monotonically with both social information and infection prevalence; and (2) if social information increases host survival, then increased sociality may correlate with decreased virulence. The context-dependent relationships that we have shown to exist between the ecology of information/epidemiological dynamics and the evolution of sociality/virulence highlights the need to understand host and parasite evolution in the context of multiple ecological processes and that simulations of disease spread on networks are likely to produce limited insights on the costs and benefits of sociality.

## Acknowledgements

We thank Ben Sheldon for comments on an earlier version of the manuscript. BA was supported by the Natural Environment Research Council (grant numbers NE/N014979/1 and NE/V003909/1). DRF was funded by a grant from the European Research Council (ERC) under the European Union’s Horizon 2020 research and innovation programme (grant agreement No. 850859) and an Eccellenza Professorship Grant of the Swiss National Science Foundation (Grant Number PCEFP3_187058), and received additional support from the Deutsche Forschungsgemeinschaft (DFG, German Research Foundation) under Germany’s Excellence Strategy - EXC 2117 - 422037984.

## Data availability

Source code is available in the *Supplementary Material* and at the following Github repository: https://github.com/ecoevogroup/Ashby_and_Farine_2020.

## References

Adelman, J. S., S. C. Moyers, D. R. Farine, and D. M. Hawley. 2015. Feeder use predicts both acquisition and transmission of a contagious pathogen in a North American songbird. Proc. Biol. Sci. 282: 20151429.

Alberts, S. C. 2019. Social influences on survival and reproduction: Insights from a long-term study of wild baboons. J. Anim. Ecol. 88:47–66.

Albery, G. F., L. Kirkpatrick, J. A. Firth, and S. Bansal. 2021. Unifying spatial and social network analysis in disease ecology. J. Anim. Ecol. 90:45–61.

Alizon, S., A. Hurford, N. Mideo, and M. Van Baalen. 2009. Virulence evolution and the trade-off hypothesis: History, current state of affairs and the future. J. Evol. Biol. 22:245–259.

Allen, J., M. Weinrich, W. Hoppitt, and L. Rendell. 2013. Network-Based Diffusion Analysis Reveals Cultural Transmission of Lobtail Feeding in Humpback Whales. Science. 340:485–488.

Aplin, L. M., D. R. Farine, R. P. Mann, and B. C. Sheldon. 2014. Individual-level personality influences social foraging and collective behaviour in wild birds. Proc. R. Soc. B 281:20141016.

Aplin, L. M., D. R. Farine, J. Morand-Ferron, A. Cockburn, A. Thornton, and B. C. Sheldon. 2015a. Experimentally induced innovations lead to persistent culture via conformity in wild birds. Nature 518:538–541.

Aplin, L. M., D. R. Farine, J. Morand-Ferron, E. F. Cole, A. Cockburn, and B. C. Sheldon. 2013. Individual personalities predict social behaviour in wild networks of great tits (Parus major). Ecol. Lett. 16:1365–1372.

Aplin, L. M., D. R. Farine, J. Morand-Ferron, and B. C. Sheldon. 2012. Social networks predict patch discovery in a wild population of songbirds. Proc. R. Soc. B 279:4199–4205.

Aplin, L. M., J. A. Firth, D. R. Farine, B. Voelkl, R. A. Crates, A. Culina, C. J. Garroway, C. A. Hinde, L. R. Kidd, I. Psorakis, N. D. Milligan, R. Radersma, B. L. Verhelst, and B. C. Sheldon. 2015b. Consistent individual differences in the social phenotypes of wild great tits, Parus major. Anim. Behav. 108:117–127.

Armansin, N. C., A. J. Stow, M. Cantor, S. T. Leu, J. A. Klarevas-Irby, A. A. Chariton, and D. R. Farine. 2020. Social Barriers in Ecological Landscapes: The Social Resistance Hypothesis. Trends Ecol. Evol. 35:137–148.

Ashby, B., and M. Boots. 2015. Coevolution of parasite virulence and host mating strategies. Proc. Natl. Acad. Sci. 112:13290–13295.

Ashby, B., and M. Boots. 2017. Multi-mode fluctuating selection in host-parasite coevolution. Ecol. Lett. 20:357–365.

Ashby, B., and S. Gupta. 2013. Sexually transmitted infections in polygamous mating systems. Philos. Trans. R. Soc. Lond. B. Biol. Sci. 368:20120048.

Ashby, B., R. Iritani, A. Best, A. White, and M. Boots. 2019. Understanding the role of eco-evolutionary feedbacks in host-parasite coevolution. J. Theor. Biol. 464:115–125.

Ashby, B., and K. C. King. 2015. Diversity and the maintenance of sex by parasites. J. Evol. Biol. 28:511–520.

Ashby, B., and K. C. King. 2017. Friendly foes: The evolution of host protection by a parasite. Evol. Lett. 1:211–221.

Barton, R. A., R. W. Byrne, and A. Whiten. 1996. Ecology, feeding competition and social structure in baboons. Behav. Ecol. Sociobiol. 38:321–329.

Bonds, M. H., D. C. Keenan, A. J. Leidner, and P. Rohani. 2005. Higher disease prevalence can induce greater sociality: a game theoretic coevolutionary model. Evolution. 59:1859–1866.

Boogert, N. J., D. R. Farine, and K. A. Spencer. 2014. Developmental stress predicts social network position. Biol. Lett. 10:20140561.

Brandl, H. B., D. R. Farine, C. Funghi, W. Schuett, and S. C. Griffith. 2019. Early-life social environment predicts social network position in wild zebra finches. Proc. R. Soc. B Biol. Sci. 286.

Brownlie, J. C., and K. N. Johnson. 2009. Symbiont-mediated protection in insect hosts. Trends Microbiol. 17:348–354.

Buckingham, L. J., and B. Ashby. 2022. Coevolutionary theory of hosts and parasites. J. Evol. Biol. 205–224.

Camerlenghi, E., A. McQueen, K. Delhey, C. N. Cook, S. A. Kingma, D. R. Farine, and A. Peters. 2022. Cooperative breeding and the emergence of multilevel societies in birds. Ecol. Lett. 1–12.

Cantor, M., L. M. Aplin, and D. R. Farine. 2020. A primer on the relationship between group size and group performance. Anim. Behav. 166:139–146.

Cantor, M., M. Chimento, S. Q. Smeele, P. He, D. Papageorgiou, L. M. Aplin, and D. R. Farine. 2021a. Social network architecture and the tempo of cumulative cultural evolution. Proc. R. Soc. B Biol. Sci. 288.

Cantor, M., A. A. Maldonado-Chaparro, K. B. Beck, H. B. Brandl, G. G. Carter, P. He, F. Hillemann, J. A. Klarevas-Irby, M. Ogino, D. Papageorgiou, L. Prox, and D. R. Farine. 2021b. The importance of individual-to-society feedbacks in animal ecology and evolution. J. Anim. Ecol. 90:27–44.

Couzin, I. D. 2006. Behavioral ecology: Social organization in fission-fusion societies. Curr. Biol. 16:R169–R171.

Croft, D. P., J. Krause, S. K. Darden, I. W. Ramnarine, J. J. Faria, and R. James. 2009. Behavioural trait assortment in a social network: patterns and implications. Behav. Ecol. Sociobiol. 63:1495–1503.

Danchin, E., L.-A. Giraldeau, T. J. Valone, and R. H. Wagner. 2004. Public information: from nosy neighbors to cultural evolution. Science. 305:487–491.

Doligez, B., E. Danchin, and J. Clobert. 2002. Public information and breeding habitat selection in a wild bird population. Science. 297:1168–1170.

Eames, K. T. D., and M. J. Keeling. 2006. Coexistence and specialization of pathogen strains on contact networks. Am. Nat. 168:230–241.

Elgar, M. A. 1986. The establishment of foraging flocks in house sparrows: risk of predation and daily temperature. Behav. Ecol. Sociobiol. 19:433–438.

Evans, J. C., M. J. Silk, N. J. Boogert, and D. J. Hodgson. 2020. Infected or informed? Social structure and the simultaneous transmission of information and infectious disease. Oikos 129:1271–1288.

Ewald, P. W. 1994. Evolution of infectious disease. Oxford Univ Press, Oxford, UK.

Farine, D. 2017. The dynamics of transmission and the dynamics of networks. J. Anim. Ecol. 86:415–418.

Farine, D. R. 2014. Measuring phenotypic assortment in animal social networks: Weighted associations are more robust than binary edges. Anim. Behav. 89:141–153.

Farine, D. R., L. M. Aplin, B. C. Sheldon, and W. Hoppitt. 2015a. Interspecific social networks promote information transmission in wild songbirds. Proc. Biol. Sci. 282:20142804.

Farine, D. R., J. A. Firth, L. M. Aplin, R. a. Crates, A. Culina, C. J. Garroway, C. a. Hinde, L. R. Kidd, N. D. Milligan, I. Psorakis, R. Radersma, B. Verhelst, B. B. Voelkl, B. C. Sheldon, L. M. Aplina, R. a. Crates, A. Culina, C. J. Garroway, C. a. Hinde, L. R. Kidd, N. D. Milligan, I. Psorakis, R. Radersma, B. Verhelst, B. B. Voelkl, and B. C. Sheldon. 2015b. The role of social and ecological processes in structuring animal populations: a case study from automated tracking of wild birds. R. Soc. Open Sci. 2:150057.

Ford, S. A., and K. C. King. 2016. Harnessing the Power of Defensive Microbes: Evolutionary Implications in Nature and Disease Control. PLOS Pathog. 12:e1005465.

Funk, S., M. Salathé, and V. A. A. Jansen. 2010. Modelling the influence of human behaviour on the spread of infectious diseases: a review. J. R. Soc. Interface 7:1247–1256.

Galef, B. G., and L.-A. Giraldeau. 2001. Social influences on foraging in vertebrates: causal mechanisms and adaptive functions. Anim. Behav. 61:3–15.

Gareta García, M., D. R. Farine, C. Brachotte, C. Borgeaud, and R. Bshary. 2021. Wild female vervet monkeys change grooming patterns and partners when freed from feeding constraints. Anim. Behav. 181:117–136.

Geritz, S. A. H., E. Kisdi, G. Meszena, and J. A. J. Metz. 1998. Evolutionarily singular strategies and the adaptive growth and branching of the evolutionary tree. Evol. Ecol. 12:35–37.

Grear, D. A., L. T. Luong, P. J. Hudson, D. A. Grear, L. T. Luong, and P. J. Hudson. 2013. Network transmission inference: Host behavior and parasite life cycle make social networks meaningful in disease ecology. Ecol. Appl. 23:1906–1914.

Grueter, C. C., X. Qi, D. Zinner, T. Bergman, M. Li, Z. Xiang, P. Zhu, A. B. Migliano, A. Miller, M. Krützen, J. Fischer, D. I. Rubenstein, T. N. C. Vidya, B. Li, M. Cantor, and L. Swedell. 2020. Multilevel Organisation of Animal Sociality. Trends Ecol. Evol. 35:834–847.

Haldane, J. B. S. 1949. Disease and evolution. La Ric. Sci. 19:68–76.

Harrison, E., and M. A. Brockhurst. 2012. Plasmid-mediated horizontal gene transfer is a coevolutionary process. Trends Microbiol. 20:262–267.

He, P., A. A. Maldonado-Chaparro, and D. R. Farine. 2019. The role of habitat configuration in shaping social structure: a gap in studies of animal social complexity. Behav. Ecol. Sociobiol. 73:9.

He, P., P.-O. Montiglio, M. Somveille, M. Cantor, and D. R. Farine. 2021. The role of habitat configuration in shaping animal population processes: a framework to generate quantitative predictions. Oecologia 2020.07.30.228205.

Hillemann, F., E. F. Cole, B. C. Sheldon, and D. R. Farine. 2020. Information use in foraging flocks of songbirds: no evidence for social transmission of patch quality. Anim. Behav. 165:35–41.

Iritani, R. 2015. How parasite-mediated costs drive the evolution of disease state-dependent dispersal. Ecol. Complex. 21:1–13.

Kenward, R. E. 1978. Hawks and Doves: Factors Affecting Success and Selection in Goshawk Attacks on Woodpigeons. J. Anim. Ecol. 47:449–460.

Krause, J., and G. D. Ruxton. 2002. Living in groups. Oxford University Press, Oxford, UK.

Kulahci, I. G., and J. L. Quinn. 2019. Dynamic Relationships between Information Transmission and Social Connections. Trends Ecol. Evol. 34:545–554.

Lion, S., and M. Boots. 2010. Are parasites “‘prudent’“ in space? Ecol. Lett. 13:1245–1255.

Lloyd-Smith, J. O., S. J. Schreiber, P. E. Kopp, and W. M. Getz. 2005. Superspreading and the effect of individual variation on disease emergence. Nature 438:355–339.

Modlmeier, A. P., J. E. Liebmann, and S. Foitzik. 2012. Diverse societies are more productive: a lesson from ants. Proc. R. Soc. B 279:2142–2150.

Papageorgiou, D., C. Christensen, G. E. C. Gall, J. A. Klarevas-Irby, B. Nyaguthii, I. D. Couzin, and D. R. Farine. 2019. The multilevel society of a small-brained bird. Curr. Biol. 29:R1120–R1121.

Pike, T. W., M. Samanta, J. Lindström, and N. J. Royle. 2008. Behavioural phenotype affects social interactions in an animal network. Proc. R. Soc. B 275:2515–2520.

Pöysä, H. 1992. Group foraging in patchy environments: the importance of coarse-level local enhancement. Ornis Scand. 23:159–166.

Prado, F., A. Sheih, J. D. West, and B. Kerr. 2009. Coevolutionary cycling of host sociality and pathogen virulence in contact networks. J. Theor. Biol. 261:561–569.

Rafaluk-mohr, C., B. Ashby, D. A. Dahan, and K. C. King. 2018. Mutual fitness benefits arise during coevolution in a nematode-defensive microbe model. Evol. Lett. 2:246–256.

Romano, V., A. J. J. MacIntosh, and C. Sueur. 2020. Stemming the Flow: Information, Infection, and Social Evolution. Trends Ecol. Evol. 35:849–853.

Romano, V., C. Sueur, and A. J. J. MacIntosh. 2021. The tradeoff between information and pathogen transmission in animal societies. Oikos 1–11.

Rosenthal, S. B., C. R. Twomey, A. T. Hartnett, H. S. Wu, and I. D. Couzin. 2015. Revealing the hidden networks of interaction in mobile animal groups allows prediction of complex behavioral contagion. Proc. Natl. Acad. Sci. U. S. A. 112:4690–4695.

Sachser, N. 1986. Different forms of social organization at high and low population densities in guinea pigs. Behaviour 97:253–272.

Sah, P., S. T. Leu, P. C. Cross, P. J. Hudson, and S. Bansal. 2017. Unraveling the disease consequences and mechanisms of modular structure in animal social networks. Proc. Natl. Acad. Sci. U. S. A. 114:4165–4170.

Sah, P., J. Mann, and S. Bansal. 2018. Disease implications of animal social network structure: A synthesis across social systems. J. Anim. Ecol. 546–558.

Scharf, I., A. P. Modlmeier, S. Beros, and S. Foitzik. 2012. Ant societies buffer individual-level effects of parasite infections. Am. Nat. 180:671–683.

Silk, M. J., D. P. Croft, T. Tregenza, and S. Bearhop. 2014. The importance of fission-fusion social group dynamics in birds. Ibis. 156:701–715.

Silk, M. J., D. J. Hodgson, C. Rozins, D. P. Croft, R. J. Delahay, M. Boots, and R. A. McDonald. 2019. Integrating social behaviour, demography and disease dynamics in network models: Applications to disease management in eclining wildlife populations. Philos. Trans. R. Soc. B Biol. Sci. 374.

Snijders, L., S. Krause, A. N. Tump, M. Breuker, C. Ortiz, S. Rizzi, I. W. Ramnarine, J. Krause, and R. H. J. M. Kurvers. 2021. Causal evidence for the adaptive benefits of social foraging in the wild. Commun. Biol. 4:94.

Stockmaier, S., N. Stroeymeyt, E. C. Shattuck, D. M. Hawley, L. A. Meyers, and D. I. Bolnick. 2021. Infectious diseases and social distancing in nature. Science. 371:eabc8881.

Tanner, C. J., and A. L. Jackson. 2012. Social structure emerges via the interaction between local ecology and individual behaviour. J. Anim. Ecol. 81:260–277.

Udiani, O., and N. H. Fefferman. 2020. How disease constrains the evolution of social systems. Proceedings. Biol. Sci. 287:20201284.

Valone, T. J. 2007. From eavesdropping on performance to copying the behavior of others: a review of public information use. Behav. Ecol. Sociobiol. 62:1–14.

van Baalen, M. 1998. Coevolution of recovery ability and virulence. Proc. R. Soc. B 265:317–325.

Watters, J., and A. Sih. 2005. The mix matters: behavioural types and group dynamics in water striders. Behaviour 142:1417–1431.

Weber, N., S. P. Carter, S. R. X. Dall, R. J. Delahay, J. L. McDonald, S. Bearhop, and R. A. McDonald. 2013. Badger social networks correlate with tuberculosis infection. Curr. Biol. 23:R915–R916.

