## Supplementary material for "Social Information use Shapes the Coevolution of Sociality and Virulence"

### A. FULL MODEL

##### *A.1: BASIC REPRODUCTIVE RATIO: SOCIAL INFORMATION*

To determine the viability of social information transmission, consider the dynamics when social information is rare:

$$\frac{dS_G}{dt} = \frac{E^2}{N}(\tau S_P(S_G + I_G) - \beta S_G I_P) - adS_G + \gamma I_G - \sigma S_G \quad \text{(S1a)}$$

$$\frac{dI_G}{dt} = \frac{E^2}{N}\big(\beta S_G I_P + \tau I_P(S_G + I_G)\big) - (ad + \alpha + \gamma + \sigma)I_G \quad \text{(S1b)}$$

One can use the next-generation method (Hurford et al. 2010) to calculate the basic reproductive ratio for social information, $R_0^I$, which first requires deriving the Jacobian of this system:

$$J = \begin{pmatrix} \frac{E^2}{N}(\tau S_P - \beta I_P) - ad - \sigma & \frac{E^2 \tau S_P}{N} + \gamma \\ \frac{E^2}{N} I_P(\beta + \tau) & \frac{E^2 \tau I_P}{N} - (ad + \alpha + \gamma + \sigma) \end{pmatrix} \quad (S2)$$

Next, one must separate the Jacobian into components, $F$ and $V$ such that $J = F - V$ with:

$$F = \frac{E^2 \tau}{N}\begin{pmatrix} S_P & S_P \\ I_P & I_P \end{pmatrix} \quad (S3)$$

$$V = \begin{pmatrix} \frac{E^2 \beta I_P}{N} + ad + \sigma & -\gamma \\ -\frac{E^2 \beta I_P}{N} & ad + \alpha + \gamma + \sigma \end{pmatrix} \quad (S4)$$

The next-generation matrix is given by $N_G = FV^{-1}$, and the basic reproductive ratio for social information is equal to the largest eigenvalue:

$$R_0^I = \frac{\tau E^2 \left(\Gamma_\gamma^I I_P + S_P(\Gamma_\alpha^I + \gamma)\right) N + \beta E^2 I_P (I_P + S_P)}{N\big((ad + \sigma)(\Gamma_\alpha^I + \gamma)N + \beta E^2 I_P \Gamma_\alpha^I\big)} \quad \text{(S5)}$$

where $\Gamma_\alpha^I = ad + \alpha + \sigma$ and $\Gamma_\gamma^I = ad + \gamma + \sigma$.

##### *A.2: BASIC REPRODUCTIVE RATIO: DISEASE*

We now follow the same method described above, but for disease. First consider the dynamics when disease is rare:

$$\frac{dI_P}{dt} = \frac{E^2}{N}(\beta S_P(I_P + I_G) - \tau I_P S_G) - (d + \alpha + \gamma)I_P + \sigma I_G \quad \text{(S6a)}$$

$$\frac{dI_A}{dt} = \frac{E^2}{N}(\beta S_G(I_P + I_G) + \tau I_P S_G) - (ad + \alpha + \gamma + \sigma)I_G \quad \text{(S6b)}$$

The Jacobian of this system is:

$$J = \begin{pmatrix} \frac{E^2}{N}(\beta S_P - \tau S_G) - (d + \alpha + \gamma) & \frac{E^2 \beta S_P}{N} + \sigma \\ \frac{E^2}{N} S_G(\beta + \tau) & \frac{E^2 \beta S_G}{N} - (ad + \alpha + \gamma + \sigma) \end{pmatrix} \quad (S7)$$

Separating the Jacobian into components, $F$ and $V$ such that $J = F - V$:

$$F = \frac{E^2 \beta}{N}\begin{pmatrix} S_P & S_P \\ S_G & S_G \end{pmatrix} \quad (S8)$$

$$V = \begin{pmatrix} \frac{E^2 \tau S_G}{N} + d + \alpha + \gamma & -\sigma \\ -\frac{E^2 \tau S_G}{N} & ad + \alpha + \gamma + \sigma \end{pmatrix} \quad (S9)$$

The next-generation matrix is again given by $N_G = FV^{-1}$, and the basic reproductive ratio for disease is equal to the largest eigenvalue:

$$R_0^D = \frac{\beta E^2\big((\Gamma_D + \sigma)S_G + S_P(\Gamma_\alpha^I + \gamma)\big)N + \tau E^2 S_G(S_P + S_G))}{N(\Gamma_D(\Gamma_\alpha^I + \gamma)N + \tau E^2 S_G \Gamma_\alpha^I)} \quad \text{(S10)}$$

where $\Gamma_D = d + \alpha + \gamma$.

##### *A.3: HOST FITNESS*

Consider the dynamics of a rare host mutant with contact effort $c_m \approx c$ into an established resident population at equilibrium (denoted by asterisks):

$$\frac{dS_P^m}{dt} = (b - qN^*)N_m - \frac{E_m E S_P^m}{N^*}\left(\beta(I_P^* + I_G^*) + \tau(S_G^* + I_G^*)\right) - dS_P^m + \gamma I_P^m + \sigma S_G^m \quad (S11\text{a})$$

$$\frac{dS_G^m}{dt} = \frac{E_m E}{N^*}\left(\tau S_P^m(S_G^* + I_G^*) - \beta S_G^m(I_P^* + I_G^*)\right) - adS_G^m + \gamma I_G^m - \sigma S_G^m \quad (S11\text{b})$$

$$\frac{dI_P^m}{dt} = \frac{E_m E}{N^*}\left(\beta S_P^m(I_P^* + I_G^*) - \tau I_P^m(S_G^* + I_G^*)\right) - (d + \alpha + \gamma)I_P^m + \sigma I_G^m \quad (S11\text{c})$$

$$\frac{dI_G^m}{dt} = \frac{E_m E}{N^*}\left(\beta S_G^m(I_P^* + I_G^*) + \tau I_P^m(S_G^* + I_G^*)\right) - (ad + \alpha + \gamma + \sigma)I_G^m \quad (S11\text{d})$$

The Jacobian of this system is:

$$J = \begin{pmatrix} b - qN^* - \frac{C_m}{N}(\beta I^* + \tau G^*) - d & b - qN^* + \sigma & b - qN^* + \gamma & b - qN^* \\ \frac{C_m}{N}\tau G^* & -\frac{C_m}{N}\beta I^* + ad + \sigma & 0 & \gamma \\ \frac{C_m}{N}\beta I^* & 0 & -\frac{C_m}{N}\tau G^* - (d + \alpha + \gamma) & \sigma \\ 0 & \frac{C_m}{N}\beta I^* & \frac{C_m}{N}\tau G^* & -(ad + \alpha + \gamma + \sigma) \end{pmatrix} \quad (S12)$$

where $C_m = E_m E$, $G^* = S_G^* + I_G^*$ and $I^* = I_P^* + I_G^*$ for notational convenience. Separating the Jacobian into components, $F$ and $V$ such that $J = F - V$:

$$F = \begin{pmatrix} b - qN^* & b - qN^* & b - qN^* & b - qN^* \\ 0 & 0 & 0 & 0 \\ 0 & 0 & 0 & 0 \\ 0 & 0 & 0 & 0 \end{pmatrix} \quad (S13)$$

$$V = \begin{pmatrix} \frac{C_m}{N}(\beta I^* + \tau G^*) + d & -\sigma & -\gamma & 0 \\ -\frac{C_m}{N}\tau G^* & \frac{C_m}{N}\beta I^* + ad + \sigma & 0 & -\gamma \\ -\frac{C_m}{N}\beta I^* & 0 & \frac{C_m}{N}\tau A^* + d + \alpha + \gamma & -\sigma \\ 0 & -\frac{C_m}{N}\beta I^* & -\frac{C_m}{N}\tau G^* & ad + \alpha + \gamma + \sigma \end{pmatrix} \quad (S14)$$

The next-generation matrix is given by $N_G = FV^{-1}$, and the invasion fitness, $w_H(E_m, E)$, is equal to the largest eigenvalue of this matrix, which we omit here as it is a lengthy expression that offers no analytical insights. The host fitness gradient is then given by $f_H(E) = \left.\frac{\partial w_H}{\partial E_m}\right|_{E_m=E}$, and singular strategies, $E^*$, occur when $f_H(E^*) = 0$. Again, we omit the fitness gradient here as it is a lengthy expression. The singular strategy is evolutionarily stable if $\left.\frac{\partial^2 w_H}{\partial E_m^2}\right|_{E_m=E=E^*} < 0$ (giving a continuously stable strategy), otherwise it is evolutionarily unstable.

#### B. Fast social information approximation

##### B.1: Basic reproductive ratio: social information

In the fast social information approximation, the proportion of the population in the good information state $G$ for a given value of $E$, $G_E$, rapidly reaches equilibrium. Letting $G_E = \frac{1}{N}(S_G + I_G)$, we see in the main text that:

$$\frac{dG_E}{dt} \approx (\tau E^2(1 - G_E) - \sigma)G_E = 0 \qquad (S15)$$

Thus, either no individuals are in the good information state, or $G_E^* = 1 - \frac{\sigma}{\tau E^2}$ when $\sigma < \tau E^2$. The basic reproductive ratio for social information is therefore:

$$\tilde{R}_0^I = \frac{\tau E^2}{\sigma} \qquad (S16)$$

as social information can only spread when $\tilde{R}_0^I > 1$.

##### B.2: Basic reproductive ratio: disease

In the fast approximation model,

$$\frac{dI}{dt} = \frac{\beta E^2 SI}{N} - (d(1 - (1 - a)G_E^*) + \alpha + \gamma)I \qquad (S17)$$

When disease is rare ($I \approx 0, S \approx N$), we have:

$$\frac{1}{I}\frac{dI}{dt} \approx \beta E^2 - (d(1-(1-a)G_E^*) + \alpha + \gamma) \qquad (S18)$$

Rearranging $\frac{1}{I}\frac{dI}{dt} > 0$, we see that the basic reproductive ratio for disease is:

$$\tilde{R}_0^D = \frac{\beta E^2}{d(1-(1-a)G_E^*) + \alpha + \gamma} \qquad (S19)$$

##### *B.3: Host fitness*

We can approximate the invasion dynamics of a rare host mutant when social information dynamics are fast by:

$$\frac{dS_m}{dt} = (b - qN^*)N_m - \frac{\beta E_m E S_m I^*}{N^*} - d(1-(1-a)G_E^m)S_m + \gamma I_m \qquad (S20a)$$

$$\frac{dI_m}{dt} = \frac{\beta E_m E S_m I^*}{N^*} - (d(1-(1-a)G_E^m) + \alpha + \gamma)I_m \qquad (S20b)$$

where $G_E^m = \max\left(\frac{E_m(\tau E^2 - \sigma)}{E^2 E_m \tau + \sigma(E - E_m)}, 0\right)$. The Jacobian of this system is:

$$J = \begin{pmatrix} b - qN^* - \frac{\beta E_m E I^*}{N^*} - d(1-(1-a)G_E^m) & b - qN^* + \gamma \\ \frac{\beta E_m E I^*}{N^*} & -(d(1-(1-a)G_E^m) + \alpha + \gamma) \end{pmatrix} \qquad (S21)$$

Separating the Jacobian into components, $F$ and $V$ such that $J = F - V$:

$$F = \begin{pmatrix} b - qN^* & b - qN^* \\ 0 & 0 \end{pmatrix} \qquad (S22)$$

$$V = \begin{pmatrix} \frac{\beta E_m E I^*}{N^*} + d(1-(1-a)G_E^m) & -\gamma \\ -\frac{\beta E_m E I^*}{N^*} & d(1-(1-a)G_E^m) + \alpha + \gamma \end{pmatrix} \qquad (S23)$$

The next-generation matrix is again given by $N_G = FV^{-1}$, and the invasion fitness, $w_H(E_m, E)$, is equal to the largest eigenvalue of this matrix, which we omit here as it is a lengthy expression which offers no analytical insights. The process for finding the fitness gradient, singular strategies, and evolutionary stability are as described above.

##### *B.4: PARASITE FITNESS*

In the fast social information approximation, the invasion dynamics of the rare parasite mutant $(M)$ with traits $\alpha_M$ and $\beta(\alpha_M)$, are given by:

$$\frac{dI_M}{dt} = \frac{\beta(\alpha_M)E^2S^*I_M}{N^*} - (d(1-(1-a)G_E^*) + \alpha_M + \gamma)I_M \qquad (S24)$$

Parasite invasion fitness is then simply equal to the per-capita growth rate when rare, $w_P(\alpha_M, \alpha) = \frac{1}{I_M}\frac{dI_M}{dt}$. The fitness gradient, $f_P(\alpha) = \left.\frac{\partial w_p}{\partial \alpha_M}\right|_{\alpha_M=\alpha}$, is then:

$$f_P(\alpha) = \frac{d\beta}{d\alpha}\left(\frac{\tau E^2(ad + \alpha + \gamma) + d\sigma(1-a)}{\beta(\alpha)\tau E^2}\right) - 1 \qquad (S25)$$

A singular strategy, $\alpha^*$, occurs when $f_P(\alpha^*) = 0$, which requires:

$$\frac{d\beta}{d\alpha} = \frac{\beta(\alpha)\tau E^2}{\tau E^2(ad + \alpha + \gamma) + d\sigma(1-a)} \qquad (S26)$$

Now consider the basic reproductive ratio from equation (S19), with $\beta = \beta(\alpha)$:

$$\tilde{R}_0^D(\alpha) = \frac{\beta(\alpha)E^2}{d(1-(1-a)G_E^*) + \alpha + \gamma} \qquad (S27)$$

where $G_E^* = 1 - \frac{\sigma}{\tau E^2}$ when $\sigma < \tau E^2$ and is 0 otherwise. Taking the derivative of $\tilde{R}_0^D(\alpha)$, setting this to 0 and rearranging, we find that:

$$\frac{d\beta}{d\alpha} = \frac{\beta(\alpha)\tau E^2}{\tau E^2(ad+\alpha+\gamma)+d\sigma(1-a)} \qquad (S28)$$

which is identical to equation (S27). Hence, evolution will maximise $\tilde{R}_0^D(\alpha)$.

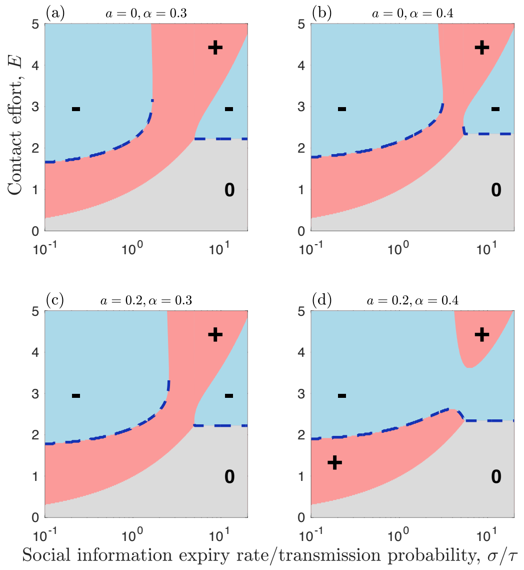


**Figure S1. The evolution of sociality arising from social information and disease transmission (fast information approximation).** Evolution of contact effort, $E$, as a function of the social information expiry rate divided by the transmission probability, $\sigma/\tau$, and for different values of the benefits of social information, $a$, and costs of disease, $\alpha$. Pink (+) and blue (-) regions indicate when the fitness gradient is positive ($E$ will increase) and negative ($E$ will decrease), respectively. Grey (0) regions indicate when there is no selection as social information and disease are both absent from the population. Blue dashed curves indicate stable or semi-stable levels of contact effort. Fixed parameters as in Fig. 2, except $\beta = 0.2$.

#### References

Hurford, A., D. Cownden, and T. Day. 2010. Next-generation tools for evolutionary invasion analyses. J. R. Soc. Interface 7:561–571.
