## Supplementary material for "Social Information use Shapes the Coevolution of Sociality and Virulence": Source code: fig1.pptx

### Slide 1
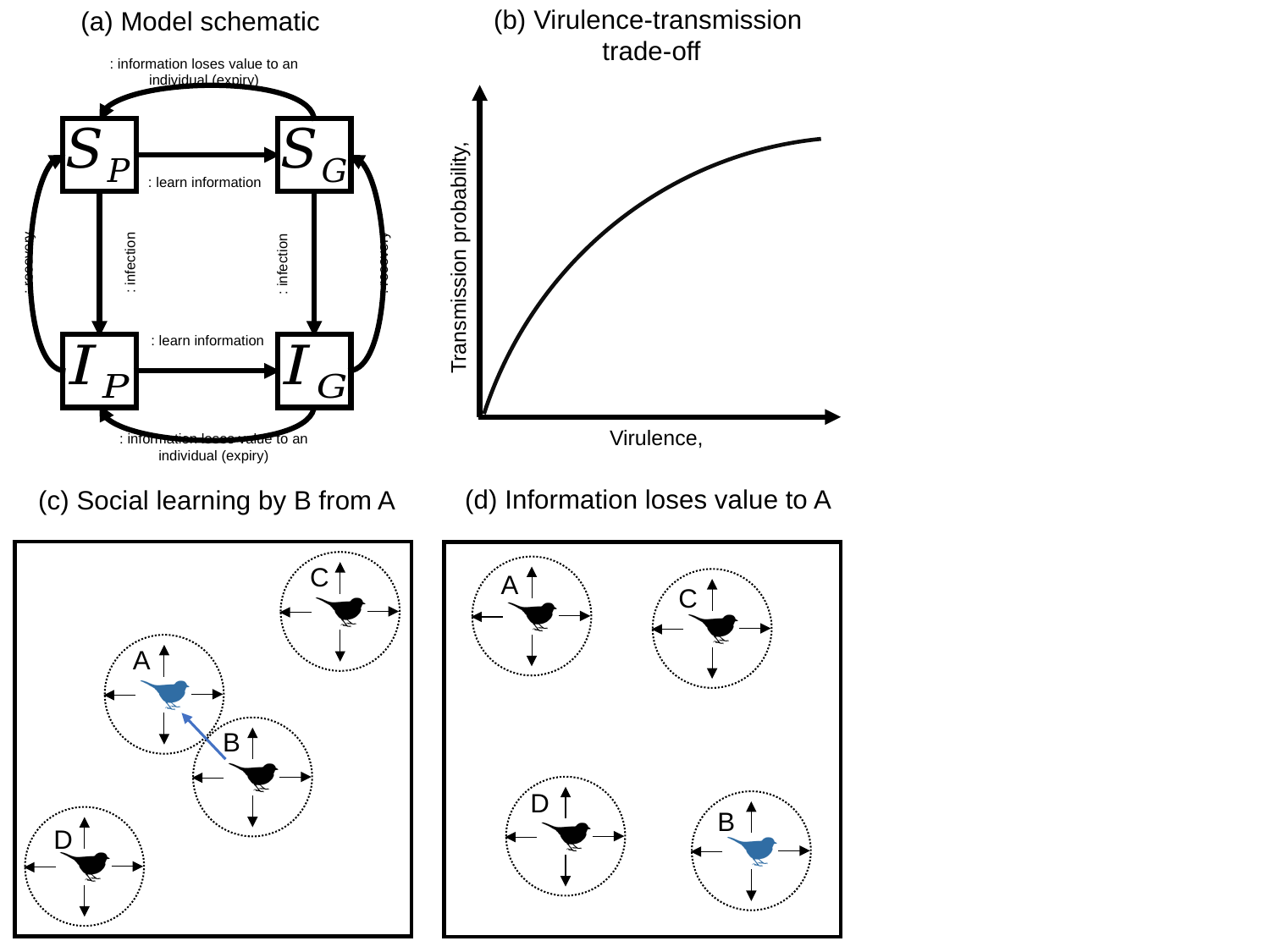

(b) Virulence-transmission
trade-off
(a) Model schematic
(d) Information loses value to A
(c) Social learning by B from A
C
A
C
A
B
D
B
D
