## Supplementary material for "Social Information use Shapes the Coevolution of Sociality and Virulence": Source code: fig2.m

```
function fig2

a = 0.2;
b = 1;
d = 0.5;
q = 1e-3;
alpha = 0.4;
beta = 0.1;
gamma = 0.2;
sigma = 1;
betamult = 1;
tau = betamult*beta;
sigtau = sigma/tau;

E = linspace(1,20,1001);
ymax = 100;
E(E==0)=[];

figure(2)
clf
set(gcf,'color','w')
set(gcf,'PaperUnits','centimeters')
xSize = 8; ySize = 8;
xLeft = (21-xSize)/2; yTop = (30-ySize)/2;
set(gcf,'PaperPosition',[xLeft yTop xSize ySize])
set(gcf,'Position',[10 100 xSize*50 ySize*50])

labs1 = {'(a)','(b)'};

%%% Approximation
GEE = max(0,1-(sigma./(tau.*E.^2)));
R0_I_approx = (tau.*E.^2)/sigma;
R0_D_approx = (beta*E.^2)./(d*(1-(1-a)*GEE)+alpha+gamma);

%%% Full model
% Starting from disease-free and information-free states
R0_I_DF = (tau.*E.^2)/(a*d+sigma);
R0_D_IF = (beta.*E.^2)/(d+alpha+gamma);

% Starting from disease or information present
SP = (d + alpha + gamma).*(alpha.^2 + (-beta.*E.^2 + d + gamma).*alpha + E.^2.*beta.*(b - d))./(beta.^2.*E.^4.*q);
IP = max(0,(alpha.^2 + (-beta.*E.^2 + d + gamma).*alpha + E.^2.*beta.*(b - d)).*(beta.*E.^2 - alpha - d - gamma)./(beta.^2.*E.^4.*q));
list = SP<=0 | IP<=0;
SP(list) = (b-d)/q;
IP(list) = 0;
N = SP+IP;
R0_I = tau.*E.^2.*(((a.*d + gamma + sigma).*IP + SP.*(a.*d + alpha + gamma + sigma)).*N + E.^2.*IP.*beta.*(IP + SP))./(N.*((a.*d + sigma).*(a.*d + alpha + gamma + sigma).*N + E.^2.*IP.*beta.*(a.*d + alpha + sigma)));
R0_I(IP==0) = R0_I_DF(IP==0);

SP = (a.*d + sigma).*((a.^2 - a).*d.^2 + ((-E.^2.*tau + sigma).*a - sigma).*d + b.*E.^2.*tau)./(E.^4.*q.*tau.^2);
SG = max(0,-(-E.^2.*tau + a.*d + sigma).*((a.^2 - a).*d.^2 + ((-E.^2.*tau + sigma).*a - sigma).*d + b.*E.^2.*tau)./(E.^4.*q.*tau.^2));
list = SP<=0 | SG<=0;
SP(list) = (b-d)/q;
SG(list) = 0;
N = SP+SG;
R0_D = beta.*(((sigma + d + alpha + gamma).*SG + SP.*(a.*d + alpha + gamma + sigma)).*N + E.^2.*SG.*tau.*(SP + SG)).*E.^2./(((d + alpha + gamma).*(a.*d + alpha + gamma + sigma).*N + E.^2.*SG.*tau.*(a.*d + alpha + gamma)).*N);
R0_D(SG==0) = R0_D_IF(SG==0);

subplot(2,2,1)
hold on
plot(E,R0_I,'k','linewidth',1.5)
plot(E,R0_I_DF,'b:','linewidth',1.5)
plot(E,R0_I_approx,'r--','linewidth',1.5)
plot(E,R0_I_DF,'b:','linewidth',1.5)
box on
set(gca,'yscale','log')
set(gca,'fontsize',10)
ylim([1,ymax])
text(0,140,labs1{1},'interpreter','latex','fontsize',12)
ylabel('Basic reproductive ratio','interpreter','latex','fontsize',16);
title('Information','interpreter','latex','fontsize',14);

L = legend('$R_0^k$','$\hat{R}_0^k$','$\tilde{R}_0^k$','fontsize',8);
set(L,'interpreter','latex')
temp = get(L,'position');
temp(1) = 0.15;
temp(2) = 0.82;
set(L,'position',temp)
x1=xlabel('Contact effort, $E$','interpreter','latex','fontsize',16);
temp=get(x1,'position');
temp(1) = temp(1) + 13;
set(x1,'position',temp);

subplot(2,2,2)
hold on
plot(E,R0_D,'k','linewidth',1.5)
plot(E,R0_D_approx,'r--','linewidth',1.5)
plot(E,R0_D_IF,'b:','linewidth',1.5)
box on
set(gca,'fontsize',10)
set(gca,'yscale','log')
ylim([1,ymax])
text(0,140,labs1{2},'interpreter','latex','fontsize',12)
title('Infection','interpreter','latex','fontsize',14);

%%%%%%%%%%%%%%%%%%%%%%%%%%%%%%%%%%%%%%%%%%%%%%%%%%%%%%
%%%%%%%%%%%%%%%%%%%%%%%%%%%%%%%%%%%%%%%%%%%%%%%%%%%%%%

% Parameters
t_max = 1e4;
E = 4;
eqtol = 1e-3;
init_pop1 = ((b-d)/(4*q)*ones(1,4));
init_pop2 = ((b-d)/(2*q)*ones(1,2));
RES = 1001;

TAU = beta*logspace(-1,1,RES);
SIGMA = sigtau*TAU;
OUT1 = zeros(RES,4);
OUT2 = zeros(RES,4);

for i=1:RES
    tau=TAU(i);
    sigma=SIGMA(i);
    
    GE =  max(0,1 - sigma/(E^2*tau));
    
    [~,x1] = sociality_slowinfo_mex(t_max,a,b,E,d,q,alpha,beta,gamma,sigma,tau,eqtol,init_pop1);
    [~,x2] = sociality_fastinfo_mex(t_max,a,b,E,d,q,alpha,beta,gamma,sigma,tau,eqtol,init_pop2);
    
    OUT1(i,:) = x1(end,:)./sum(x1(end,:));
    OUT2(i,:) = [x2(end,1)*[1-GE,GE],x2(end,2)*[1-GE,GE]]./sum(x2(end,:));
end

DISPREV1 = sum(OUT1(:,3:4),2);
DISPREV2 = sum(OUT2(:,3:4),2);
INFOPREV1 = sum(OUT1(:,[2,4]),2);
INFOPREV2 = sum(OUT2(:,[2,4]),2);

cols = [166,206,227
    31,120,180
    178,223,138
    51,160,44]/255;

cols2 = [27,158,119
    117,112,179]/255;

labs2 = {'(c)','(d)'};
subplot(2,2,3)
hold on
for i=1:4
    plot(TAU/beta,OUT1(:,i),'color',cols(i,:),'linewidth',1.5)
end
for i=1:4
    plot(TAU/beta,OUT2(:,i),'--','color',cols(i,:),'linewidth',1.5)
end
set(gca,'ytick',0:0.2:1)
set(gca,'xscale','log')
set(gca,'fontsize',10)
ylim([0,0.7])
box on
ylabel('Proportion','interpreter','latex','fontsize',16)
text(TAU(1)/beta,1.07*0.7,labs2{1},'interpreter','latex','fontsize',12)
% text(TAU(end)/beta,OUT2(end,1)+0.02,'$\frac{S_P}{N}$','interpreter','latex','fontsize',9)
% text(TAU(end)/beta,OUT2(end,2)-0.02,'$\frac{S_G}{N}$','interpreter','latex','fontsize',9)
% text(TAU(end)/beta,OUT2(end,3)+0.03,'$\frac{I_P}{N}$','interpreter','latex','fontsize',9)
% text(TAU(end)/beta,OUT2(end,4)-0.03,'$\frac{I_G}{N}$','interpreter','latex','fontsize',9)
L=legend('$\frac{S_P}{N}$','$\frac{S_G}{N}$','$\frac{I_P}{N}$','$\frac{I_G}{N}$','location','northeast','interpreter','latex','fontsize',8);
temp=get(L,'position');
temp(1) = temp(1)+0.005;
temp(2) = temp(2)+0.01;
set(L,'position',temp);
x1 = xlabel('Relative transmission probability, $\tau/\beta$','interpreter','latex','fontsize',16);
temp=get(x1,'position');
temp(1) = temp(1) + 17;
set(x1,'position',temp);

subplot(2,2,4)
hold on
plot(TAU/beta,DISPREV1,'color',cols2(1,:),'linewidth',1.5)
plot(TAU/beta,INFOPREV1,'color',cols2(2,:),'linewidth',1.5)
plot(TAU/beta,DISPREV2,'--','color',cols2(1,:),'linewidth',1.5)
plot(TAU/beta,INFOPREV2,'--','color',cols2(2,:),'linewidth',1.5)
set(gca,'ytick',0:0.2:1)
set(gca,'xscale','log')
set(gca,'fontsize',10)
ylim([0,0.42])
box on
text(TAU(1)/beta,1.07*0.42,labs2{2},'interpreter','latex','fontsize',12)
% text(TAU(end)/beta,DISPREV2(end),'$\frac{I_P+I_G}{N}$','interpreter','latex','fontsize',10)
% text(TAU(end)/beta,INFOPREV2(end)-0.02,'$\frac{S_G+I_G}{N}$','interpreter','latex','fontsize',10)
legend('infected','informed','location','southeast','interpreter','latex','fontsize',8)

% save2pdf('fig2.pdf');
```
