## Supplementary material for "Social Information use Shapes the Coevolution of Sociality and Virulence": Source code: fig3.m

```
function fig3

A0 = [0,0.2];
ALPHA0 = [0.3,0.4];
% A0 = [0,0.2];
% ALPHA0 = [0.05,0.1];

d = 0.5;
beta = 0.2;
gamma = 0.2;
betamult = 1;
tau = betamult*beta;

E = 4;
xres = 1001;
yres = 1000;

infoprev = linspace(0,1,xres);
disprev = linspace(0,1,yres);

figure(3)
clf
set(gcf,'color','w')
set(gcf,'PaperUnits','centimeters')
xSize = 12; ySize = 5;
xLeft = (21-xSize)/2; yTop = (30-ySize)/2;
set(gcf,'PaperPosition',[xLeft yTop xSize ySize])
set(gcf,'Position',[10 100 xSize*50 ySize*50])

labs = {'(a)','(b)'};
for sub=1:2
    a = A0(sub);
    alpha = ALPHA0(sub);
    sigtau = E^2*(1-infoprev);
    sigma = sigtau*tau;
    [sigma0,disprev0] = meshgrid(sigma,disprev);
    beta = -(a*E^2*d*tau + alpha*E^2*tau + E^2*gamma*tau - a*d*sigma0 + d*sigma0)./(E^4*tau*(disprev0 - 1));
    
    w_1=(-beta.*tau.^2.*(alpha.*(a.*d + alpha + gamma).*tau + sigma0.*d.*beta.*(a - 1)).*E.^8 + (alpha.*(a.*d + alpha + gamma).^2.*tau.^2 + 2.*d.*beta.*alpha.*sigma0.*(a - 1).*tau + d.*beta.^2.*sigma0.^2.*(a - 1)).*tau.*E.^6 - 3.*d.*(a - 1).*((a.*d + alpha + gamma).*tau + beta.*sigma0./3).*alpha.*tau.*sigma0.*E.^4 + 3.*d.*(a - 1).*(alpha./3 + (a - 2./3).*d + gamma./3).*alpha.*tau.*sigma0.^2.*E.^2 - d.^2.*alpha.*sigma0.^3.*(a - 1).^2)./(beta.*(tau.*beta.*(a.*d + alpha).*E.^4 + (-alpha.*(a.*d + alpha + gamma).*tau - sigma0.*d.*beta.*(a - 1)).*E.^2 + sigma0.*d.*alpha.*(a - 1)).*E.^7.*tau.^2);
    
    subplot(1,2,sub)
    if(sub==2)
        v = -0.1:0.02:0.1;
    else
        v = [-0.2,-0.1,-0.06,-0.02,0:0.02:0.1];
    end
    [c0,h]=contourf(infoprev,disprev,w_1,v);
    set(gca,'fontsize',10)
    clabel(c0,h,'fontsize',8,'LabelSpacing',500,'color','k')
    set(h,'color','k')
    temp = get(gca,'position');
    temp(1) = temp(1)-0.03*sub;
    temp(2) = temp(2)+0.03;
    temp(4) = temp(4)-0.03;
    set(gca,'position',temp);    
    set(gca,'clim',[-0.105,0.1])
    drawnow
    
    if(sub==2)
        cols = [241,163,64
            247,247,247
            153,142,195]/255;
        [cmap]=buildcmap2(cols,256);
        colormap(cmap)
        C=colorbar;
        temp=get(C,'position');
        temp(1) = temp(1) + 0.1;
        set(C,'position',temp)
        set(C,'ytick',-0.1:0.02:0.1)
        set(C,'fontsize',8)
        ylabel(C,'Fitness gradient','interpreter','latex','fontsize',14)
    end
    if(sub==1)
        x1=xlabel('Social information prevalence','interpreter','latex','fontsize',14);
        temp=get(x1,'position');
        temp(1)=temp(1)+0.6;
        temp(2)=temp(2)+0.01;
        set(x1,'position',temp);
        ylabel('Infection prevalence','interpreter','latex','fontsize',14)
    end
    text(0,1.05,labs{sub},'interpreter','latex','fontsize',12)
    title(strcat('$a=',num2str(a),', \alpha=',num2str(alpha),'$'),'interpreter','latex','fontsize',10);
end

% save2pdf('fig3.pdf');
```
