## Supplementary material for "Social Information use Shapes the Coevolution of Sociality and Virulence": Source code: fig6.m

```
function fig6

% Parameter values
t_max = 100;
b = 1;
Emin = 0;
Emax = 4;
Estart = 1.6;
d = 0.5;
q = 1e-3;
beta = 0.2;
gamma = 0.2;
res0 = 101;
res1 = 1001;
nevol = [4000,1000]+1;
plotflag=0;
a=0.2;
alpha=0.3;
sigtau = 2;
taubeta = [1,5];

cols = [251,154,153
    171,217,233
    200,10,10
    69,117,180
    0,0,0
    0,0,0
    0,0,0
    0,0,0
    0,0,0]/255;

cols2 = [27,158,119
117,112,179]/255;

labs1 = {'(a.i)','(b.i)','(c.i)','(d.i)'};
labs2 = {'(a.ii)','(b.ii)','(c.ii)','(d.ii)'};

figure(6)
clf
set(gcf,'color','w')
set(gcf,'PaperUnits','centimeters')
xSize = 9; ySize = 8;
xLeft = (21-xSize)/2; yTop = (30-ySize)/2;
set(gcf,'PaperPosition',[xLeft yTop xSize ySize])
set(gcf,'Position',[10 100 xSize*50 ySize*50])

for sub=1:2
    tau = taubeta(sub)*beta;
    sigma = sigtau*tau;
    
    [singstrat,outcome] = singstrat_slowinfo(t_max,a,b,Emin,Emax,d,q,alpha,beta,gamma,sigma,tau,res1);
    [SOCIALITY,DISPREV,INFOPREV] = sociality_slowinfo_simulation(t_max,a,b,Emin,Emax,Estart,d,q,alpha,beta,gamma,sigma,tau,res0,nevol(sub),plotflag);
    
    a0=2;
    SOCIALITY0=log10(SOCIALITY);
    SOCIALITY0(SOCIALITY0<-a0)=-a0;
    SOCIALITY0=(SOCIALITY0+a0)/a0;
    
    subplot(2,2,sub)
    imagesc(SOCIALITY0');set(gca,'ydir','normal')
    map=colormap('gray');
    map=flipud(map);
    colormap(map);
    hold on
    
    for k=1:length(singstrat)
        plot([1,length(DISPREV)],res0*singstrat(k)*[1,1]/Emax,'--','color',cols(outcome(k),:),'linewidth',2)
    end
    set(gca,'fontsize',10)
    if(sub==1)        
        ylabel('Contact effort, $E$','interpreter','latex','fontsize',14)
    end
    ylim([1,res0])
    set(gca,'xtick',linspace(1,nevol(sub),3),'xticklabel',linspace(0,nevol(sub)-1,3))
    set(gca,'ytick',linspace(1,res0,3),'yticklabel',linspace(Emin,Emax,3))
    text(0,res0*1.07,labs1{sub},'interpreter','latex','fontsize',12)
    title(strcat('$\sigma/\gamma=',num2str(sigma/gamma),'$'),'interpreter','latex','fontsize',10)
    drawnow
    
    subplot(2,2,sub+2)
    hold on
    plot(1:length(DISPREV),DISPREV,'color',cols2(1,:),'linewidth',1.5)
    plot(1:length(DISPREV),INFOPREV,'color',cols2(2,:),'linewidth',1.5);
    ylim([0,1])
    set(gca,'ytick',[0,1])
    set(gca,'xtick',linspace(0,nevol(sub),3),'xticklabel',linspace(0,nevol(sub)-1,3))
    if(sub==1)
        legend('infected','informed','location','southeast','interpreter','latex','fontsize',8)
    end
    box on
    set(gca,'fontsize',10)
    if(sub==1)
        ylabel('Proportion','interpreter','latex','fontsize',14)
    end
    if(sub==1)                
        x1=xlabel('Evolutionary time','interpreter','latex','fontsize',16);
        temp=get(x1,'position');
        temp(1)=temp(1)+2700;
        temp(2)=temp(2)+0.02;
        set(x1,'position',temp);
    end
    text(0,1.07,labs2{sub},'interpreter','latex','fontsize',12)
    title(strcat('$\sigma/\gamma=',num2str(sigma/gamma),'$'),'interpreter','latex','fontsize',10)
    
    drawnow
end
% save2pdf('fig6.pdf')
```
