## Supplementary material for "Social Information use Shapes the Coevolution of Sociality and Virulence": Source code: fig7.m

```
function fig7

a = 0.2;
d = 0.5;
gamma = 0.2;
sigtau = 6;
tau = 0.2;
sigma = sigtau.*tau;
kappa = 1;

E = linspace(0,10,1001);
E(E==0)=[];

figure(7)
clf
set(gcf,'color','w')
set(gcf,'PaperUnits','centimeters')
xSize = 5; ySize = 4.5;
xLeft = (21-xSize)/2; yTop = (30-ySize)/2;
set(gcf,'PaperPosition',[xLeft yTop xSize ySize])
set(gcf,'Position',[10 100 xSize*50 ySize*50])

alpha = (a.*E.^2.*d.*tau + E.^2.*gamma.*tau - a.*d.*sigma + d.*sigma)./(E.^2.*tau);
beta = kappa*sqrt(alpha);
disprev = (E.^4.*tau.*beta - tau.*(a.*d + alpha + gamma).*E.^2 + d.*sigma.*(-1 + a))./(E.^4.*tau.*beta);
AE = max(0,1-(sigma./(tau.*E.^2)));

alpha(AE==0)=gamma+d;
beta(AE==0)=kappa*(sqrt(alpha(AE==0)));
disprev(AE==0)=(E(AE==0).^2.*kappa.*sqrt(alpha(AE==0)) - gamma - alpha(AE==0) - d)./(kappa.*E(AE==0).^2.*sqrt(alpha(AE==0)));

R0_D_approx = (beta.*E.^2)./(d*(1-(1-a)*AE)+alpha+gamma);

alpha(R0_D_approx<1)=NaN;
disprev(R0_D_approx<1)=NaN;

yyaxis left
plot(E,alpha,'k','linewidth',1.5)
hold on
plot(E(find(isnan(alpha),1,'last'))*[1,1],[0,ceil(10*max(alpha))/10],'k:','linewidth',1.5)
plot(E(find(AE==0,1,'last'))*[1,1],[0,ceil(10*max(alpha))/10],'k:','linewidth',1.5)
set(gca,'fontsize',10)
ylabel('Optimal virulence, $\alpha^*$','interpreter','latex','fontsize',16);
text(E(end)*0.75,alpha(end)*0.9,'$\alpha^*$','interpreter','latex','fontsize',12);
text(0.7,max(alpha)*0.75,'$\tilde{R}_0^D<1$','interpreter','latex','fontsize',12,'rotation',90);
text(1.9,max(alpha)*0.75,'$\tilde{R}_0^I<1$','interpreter','latex','fontsize',12,'rotation',90);
set(gca,'ycolor','k')

yyaxis right
plot(E,disprev,'k--','linewidth',1.5)
text(E(end)*0.75,disprev(end)*0.9,'$\frac{I_P+I_G}{N}$','interpreter','latex','fontsize',12);
set(gca,'ycolor','k')

xlabel('Contact effort, $E$','interpreter','latex','fontsize',16);
ylabel('Infection prevalence','interpreter','latex','fontsize',16);
box on

% save2pdf('fig7.pdf');
```
