## Supplementary material for "Social Information use Shapes the Coevolution of Sociality and Virulence": Source code: fig8.m

```
function fig8

if(~exist('fig8.mat','file'))
    fig8_data;
end
load('fig8.mat')
gamma = 0.2;

PARAM = SIGTAU;
cols = [171,217,233
    251,154,153
    20,50,180
    180,50,20
    220,220,220]/255;

figure(8)
clf
set(gcf,'color','w')
set(gcf,'PaperUnits','centimeters')
xSize = 18; ySize = 4.5;
xLeft = (21-xSize)/2; yTop = (30-ySize)/2;
set(gcf,'PaperPosition',[xLeft yTop xSize ySize])
set(gcf,'Position',[10 100 xSize*50 ySize*50])

labs = {'(a)','(b)','(c)'};
ymax = 5;
for i2=1:length(A)
    subplot(1,length(A),i2)
    singstrat = permute(SINGSTRAT_SIGTAU_FAST(:,i2,:),[1,3,2]);
    outcome = permute(OUTCOME_SIGTAU_FAST(:,i2,:),[1,3,2]);
    
    % rebuild arrays
    outcome2 = NaN*zeros(xres,4);
    singstrat2 = NaN*zeros(xres,4);
    for j=1:4
        list = find(sum(outcome==j,2));
        outcome2(list,j)=j;
        for k=1:length(list)
            singstrat2(list(k),j) = singstrat(list(k),outcome(list(k),:)==j);
        end
    end
    
    % Fill in direction of selection
    
    % Fill background
    patch([SIGTAU(1);SIGTAU(end);SIGTAU(end);SIGTAU(1)],[SIGTAU(1),SIGTAU(1),SIGTAU(end),SIGTAU(end)],cols(2,:),'linestyle','none')
    
    %         % fill +ve between outcome2==1 and outcome2==4 and above outcome2==3
    %
    %         pos1_lower = singstrat2(~isnan(singstrat2(:,1)),1);
    %         pos1_upper = min(ymax+1,singstrat2(:,4));
    %
    %         patch([SIGTAU(1:length(pos1_lower))';SIGTAU(length(pos1_lower):-1:1)'],[pos1_upper(1:length(pos1_lower));flipud(pos1_lower)],cols(2,:),'linestyle','none');
    %
    %         pos2_lower = neg_upper;
    %         pos2_upper = ymax+0*pos2_lower;
    %
    %         patch([SIGTAU';flipud(SIGTAU')],[pos2_lower;pos2_upper],cols(2,:),'linestyle','none')
    
    
    % fill -ve between outcome2==1 and outcome2==3
    
    % patch([(75:101)';(101:-1:75)'],[singstrat2(75:end,2);flipud(singstrat2(75:end,3))],'r')
    
    neg_upper = singstrat2(:,3); neg_upper(isnan(neg_upper))=ymax;
    neg_upper = min(ymax,neg_upper);
    
    neg_lower = sum([singstrat2(:,2),singstrat2(:,4)],2,'omitnan');
    neg_lower(neg_lower==0)=ymax+1;
    
    patch([SIGTAU';flipud(SIGTAU')],[neg_lower;flipud(neg_upper)],cols(1,:),'linestyle','none')
    
    % fill neutral between outcome2==1 and
    
    neutral_lower = zeros(length(SIGTAU),1);
    neutral_upper = min([singstrat2(:,1),singstrat2(:,2)],[],2,'omitnan');
    
    patch([SIGTAU';flipud(SIGTAU')],[neutral_lower;flipud(neutral_upper)],cols(5,:),'linestyle','none')
    
    xlim([SIGTAU(1),SIGTAU(end)])
    ylim([0,ymax])
    
    set(gca,'xtick',[0.1,1,10])
    set(gca,'xscale','log')
    box on
    set(gca,'fontsize',10)
    
    ALPHA = (singstrat2.^2.*(a.*d + gamma).*tau - d.*repmat(SIGTAU',[1,4]).*(-1 + a))./(singstrat2.^2.*tau);
    ALPHA(:,[1,3,5:9])=0;
    ALPHA(~isnan(ALPHA(:,2)),2) = gamma+d;
    ALPHA(isnan(ALPHA))=0;
    ALPHA=sum(ALPHA,2);
    ALPHA(ALPHA==0)=NaN;
    BETA = kappa*sqrt(ALPHA);
    
    yyaxis left
    hold on
%     plot(SIGTAU,sqrt(SIGTAU),'k--','linewidth',1) % Fast information threshold (no disease)
%     plot(SIGTAU,sqrt((d+ALPHA+gamma)./BETA),'k:','linewidth',1.5) % Disease threshold (no information)

    neg_lower(neg_lower>ymax)=NaN;
    plot(SIGTAU,neg_lower,'--','color',cols(3,:),'linewidth',2)
    
    title(strcat('$a=',num2str(A(i2)),'$'),'interpreter','latex','fontsize',12);
    text(SIGTAU(1),ymax*1.05,labs{i2},'fontsize',12,'interpreter','latex')
    
    if(i2==1)
        text(5,4.5,'+','fontsize',20,'fontweight','bold')
        text(0.2,3,'-','fontsize',20,'fontweight','bold')
        text(11,3,'-','fontsize',20,'fontweight','bold')
        text(11,0.5,'0','fontsize',14,'fontweight','bold')
    else
        text(5,4.5,'+','fontsize',20,'fontweight','bold')
        text(0.15,0.9,'+','fontsize',20,'fontweight','bold')
        text(0.2,3,'-','fontsize',20,'fontweight','bold')
        text(11,0.5,'0','fontsize',14,'fontweight','bold')
    end

    yyaxis right
    plot(SIGTAU,ALPHA,'k:','linewidth',2)
    set(gca,'ycolor','k')
    set(gca,'fontsize',10)
    ylim([0,1])
    
end

for i=1:3
    subplot(1,3,i)
    yyaxis left
    ylabel('Contact effort, $E$','interpreter','latex','fontsize',16);
    yyaxis right
    ylabel('Virulence, $\alpha$','interpreter','latex','fontsize',16);
end

subplot(1,3,2)
xlabel({'Social information expiry rate/transmission probability, $\sigma/\tau$'},'interpreter','latex','fontsize',16);

% save2pdf('fig8.pdf')

function fig8_data

% Fixed and default parameter values
Emin = 0;
Emax = 5;
d = 0.5;
kappa = 1;
gamma = 0.2;
xres = 10001;
res1 = 1001;
tau = 1;

% Variables
SIGTAU = logspace(-1,log10(20),xres);
A = [0,0.2,0.5];

% OUTPUTS
SINGSTRAT_SIGTAU_FAST1 = NaN*zeros(xres,length(A));
SINGSTRAT_SIGTAU_FAST2 = NaN*zeros(xres,length(A));
SINGSTRAT_SIGTAU_FAST3 = NaN*zeros(xres,length(A));
SINGSTRAT_SIGTAU_FAST4 = NaN*zeros(xres,length(A));
SINGSTRAT_SIGTAU_FAST5 = NaN*zeros(xres,length(A));
OUTCOME_SIGTAU_FAST1 = NaN*zeros(xres,length(A));
OUTCOME_SIGTAU_FAST2 = NaN*zeros(xres,length(A));
OUTCOME_SIGTAU_FAST3 = NaN*zeros(xres,length(A));
OUTCOME_SIGTAU_FAST4 = NaN*zeros(xres,length(A));
OUTCOME_SIGTAU_FAST5 = NaN*zeros(xres,length(A));
NUM_OUTCOMES_SIGTAU_FAST = NaN*zeros(xres,length(A));

% Fast info version depends on SIGTAU, so can fix tau
COUNT = 0;
TOTAL = length(A)
for i2=1:length(A)
    % Sweep over SIGTAU
    tic;
    parfor i1=1:length(SIGTAU)
        [singstrat,outcome] = singstrat_fastinfo_coevo_approx(A(i2),Emin,Emax,d,kappa,gamma,SIGTAU(i1)*tau,tau,res1);

        NUM_OUTCOMES_SIGTAU_FAST(i1,i2) = length(singstrat);
        SINGSTRAT_SIGTAU_FAST1(i1,i2) = singstrat(1);
        OUTCOME_SIGTAU_FAST1(i1,i2) = outcome(1);
        if(NUM_OUTCOMES_SIGTAU_FAST(i1,i2)>1)
            SINGSTRAT_SIGTAU_FAST2(i1,i2) = singstrat(2);
            OUTCOME_SIGTAU_FAST2(i1,i2) = outcome(2);
            if(NUM_OUTCOMES_SIGTAU_FAST(i1,i2)>2)
                SINGSTRAT_SIGTAU_FAST3(i1,i2) = singstrat(3);
                OUTCOME_SIGTAU_FAST3(i1,i2) = outcome(3);
                if(NUM_OUTCOMES_SIGTAU_FAST(i1,i2)>3)
                    SINGSTRAT_SIGTAU_FAST4(i1,i2) = singstrat(4);
                    OUTCOME_SIGTAU_FAST4(i1,i2) = outcome(4);
                    if(NUM_OUTCOMES_SIGTAU_FAST(i1,i2)>4)
                        SINGSTRAT_SIGTAU_FAST5(i1,i2) = singstrat(5);
                        OUTCOME_SIGTAU_FAST5(i1,i2) = outcome(5);
                    end
                end
            end
        end
    end
    toc;
    COUNT=COUNT+1;
    PROGRESS = COUNT/TOTAL
end

% Combine results
SINGSTRAT_SIGTAU_FAST = cat(3,SINGSTRAT_SIGTAU_FAST1,SINGSTRAT_SIGTAU_FAST2,SINGSTRAT_SIGTAU_FAST3,SINGSTRAT_SIGTAU_FAST4,SINGSTRAT_SIGTAU_FAST5);
OUTCOME_SIGTAU_FAST = cat(3,OUTCOME_SIGTAU_FAST1,OUTCOME_SIGTAU_FAST2,OUTCOME_SIGTAU_FAST3,OUTCOME_SIGTAU_FAST4,OUTCOME_SIGTAU_FAST5);

clear i1 i2 i3 i4 singstrat outcome ans COUNT TOTAL PROGRESS
clear SINGSTRAT_SIGTAU_FAST1 SINGSTRAT_SIGTAU_FAST2 SINGSTRAT_SIGTAU_FAST3 SINGSTRAT_SIGTAU_FAST4 SINGSTRAT_SIGTAU_FAST5 OUTCOME_SIGTAU_FAST1 OUTCOME_SIGTAU_FAST2 OUTCOME_SIGTAU_FAST3 OUTCOME_SIGTAU_FAST4 OUTCOME_SIGTAU_FAST5 SINGSTRAT_SIGTAU_SLOW1 SINGSTRAT_SIGTAU_SLOW2 SINGSTRAT_SIGTAU_SLOW3 SINGSTRAT_SIGTAU_SLOW4 SINGSTRAT_SIGTAU_SLOW5 OUTCOME_SIGTAU_SLOW1 OUTCOME_SIGTAU_SLOW2 OUTCOME_SIGTAU_SLOW3 OUTCOME_SIGTAU_SLOW4 OUTCOME_SIGTAU_SLOW5
save('fig8.mat')
```
