## Supplementary material for "Social Information use Shapes the Coevolution of Sociality and Virulence": Source code: figS1.m

```
function figS1

if(~exist('figS1.mat','file'))
    figS1_data;
end
load('figS1.mat')
% beta=0.2;
% gamma=0.2;
% tau=0.2;
ymax = 5;

cols = [171,217,233
    251,154,153
    20,50,180
    180,50,20
    220,220,220]/255;

figure(9)
clf
set(gcf,'color','w')
set(gcf,'PaperUnits','centimeters')
xSize = 9; ySize = 9;
xLeft = (21-xSize)/2; yTop = (30-ySize)/2;
set(gcf,'PaperPosition',[xLeft yTop xSize ySize])
set(gcf,'Position',[10 100 xSize*50 ySize*50])
colormap([1,1,1;0.5,0.5,0.5;0.75,0.75,0.75])

labs = {'(a)','(b)','(c)','(d)'};
sub=0;
for i3=1:length(A)
    for i2=1:length(ALPHA)
        
        sub=sub+1;
        subplot(length(A),length(ALPHA),sub)
        
        singstrat = permute(SINGSTRAT_SIGTAU_FAST(:,i2,i3,:),[1,4,3,2]);
        outcome = permute(OUTCOME_SIGTAU_FAST(:,i2,i3,:),[1,4,3,2]);
        
        % rebuild arrays
        outcome2 = NaN*zeros(xres,4);
        singstrat2 = NaN*zeros(xres,4);
        for j=1:4
            list = find(sum(outcome==j,2));
            outcome2(list,j)=j;
            for k=1:length(list)
                singstrat2(list(k),j) = singstrat(list(k),outcome(list(k),:)==j);
            end
        end
        
        % Fill in direction of selection
        
        % Fill background
        patch([SIGTAU(1);SIGTAU(end);SIGTAU(end);SIGTAU(1)],[SIGTAU(1),SIGTAU(1),SIGTAU(end),SIGTAU(end)],cols(2,:),'linestyle','none')
        
%         % fill +ve between outcome2==1 and outcome2==4 and above outcome2==3
%         
%         pos1_lower = singstrat2(~isnan(singstrat2(:,1)),1);
%         pos1_upper = min(ymax+1,singstrat2(:,4));
%         
%         patch([SIGTAU(1:length(pos1_lower))';SIGTAU(length(pos1_lower):-1:1)'],[pos1_upper(1:length(pos1_lower));flipud(pos1_lower)],cols(2,:),'linestyle','none');
%         
%         pos2_lower = neg_upper;
%         pos2_upper = ymax+0*pos2_lower;
%         
%         patch([SIGTAU';flipud(SIGTAU')],[pos2_lower;pos2_upper],cols(2,:),'linestyle','none')
        
        
        % fill -ve between outcome2==1 and outcome2==3
        
        % patch([(75:101)';(101:-1:75)'],[singstrat2(75:end,2);flipud(singstrat2(75:end,3))],'r')
        
        neg_upper = singstrat2(:,3); neg_upper(isnan(neg_upper))=ymax;
        neg_upper = min(ymax,neg_upper);
        
        neg_lower = sum([singstrat2(:,2),singstrat2(:,4)],2,'omitnan');
        neg_lower(neg_lower==0)=ymax+1;
        
        patch([SIGTAU';flipud(SIGTAU')],[neg_lower;flipud(neg_upper)],cols(1,:),'linestyle','none')
        
        % fill neutral between outcome2==1 and
        
        neutral_lower = zeros(length(SIGTAU),1);
        neutral_upper = min([singstrat2(:,1),singstrat2(:,2)],[],2,'omitnan');
        
        patch([SIGTAU';flipud(SIGTAU')],[neutral_lower;flipud(neutral_upper)],cols(5,:),'linestyle','none')
        
        xlim([SIGTAU(1),SIGTAU(end)])
        ylim([0,ymax])

        hold on
        neg_lower(neg_lower>ymax)=NaN;
        plot(SIGTAU,neg_lower,'--','color',cols(3,:),'linewidth',2)
        
        set(gca,'xtick',[0.1,1,10])
        set(gca,'xscale','log')
        box on
        set(gca,'fontsize',10)
        text(SIGTAU(1),ymax*1.05,labs{sub},'fontsize',12,'interpreter','latex')
        if(sub==3)
            x1=xlabel('Social information expiry rate/transmission probability, $\sigma/\tau$','interpreter','latex','fontsize',14);
            y1=ylabel('Contact effort, $E$','interpreter','latex','fontsize',14);
            temp=get(x1,'position');
            temp(1) = temp(1) + 35;
            set(x1,'position',temp);
            temp=get(y1,'position');
            temp(2) = temp(2) + 3.5;
            set(y1,'position',temp);
        end
        title(strcat('$a=',num2str(A(i3)),', \alpha=',num2str(ALPHA(i2)),'$'),'interpreter','latex','fontsize',10);
        
        if(sub==4)
            text(7,4.5,'+','fontsize',20,'fontweight','bold')
            text(0.15,1.4,'+','fontsize',20,'fontweight','bold')
            text(0.2,3,'-','fontsize',20,'fontweight','bold')
            text(11,1,'0','fontsize',14,'fontweight','bold')
        else
            text(7,4.5,'+','fontsize',20,'fontweight','bold')
            text(0.2,3,'-','fontsize',20,'fontweight','bold')
            text(11,3,'-','fontsize',20,'fontweight','bold')
            text(11,1,'0','fontsize',14,'fontweight','bold')
        end
    end
end

% save2pdf('figS1.pdf')


function figS1_data

% Fixed and default parameter values
t_max = 1000;
b = 1;
Emin = 0;
Emax = 20;
d = 0.5;
q = 1e-3;
beta = 0.2;
gamma = 0.2;
tau = beta;
xres = 1001;
res1 = 1001;

% Variables
SIGTAU = logspace(-1,log10(20),xres);
A = [0,0.2];
ALPHA = [0.3,0.4];

% OUTPUTS
SINGSTRAT_SIGTAU_FAST1 = NaN*zeros(xres,length(ALPHA),length(A));
SINGSTRAT_SIGTAU_FAST2 = NaN*zeros(xres,length(ALPHA),length(A));
SINGSTRAT_SIGTAU_FAST3 = NaN*zeros(xres,length(ALPHA),length(A));
SINGSTRAT_SIGTAU_FAST4 = NaN*zeros(xres,length(ALPHA),length(A));
SINGSTRAT_SIGTAU_FAST5 = NaN*zeros(xres,length(ALPHA),length(A));
OUTCOME_SIGTAU_FAST1 = NaN*zeros(xres,length(ALPHA),length(A));
OUTCOME_SIGTAU_FAST2 = NaN*zeros(xres,length(ALPHA),length(A));
OUTCOME_SIGTAU_FAST3 = NaN*zeros(xres,length(ALPHA),length(A));
OUTCOME_SIGTAU_FAST4 = NaN*zeros(xres,length(ALPHA),length(A));
OUTCOME_SIGTAU_FAST5 = NaN*zeros(xres,length(ALPHA),length(A));
NUM_OUTCOMES_SIGTAU_FAST = NaN*zeros(xres,length(ALPHA),length(A));

% Fast info version depends on SIGTAU, so can fix tau
COUNT = 0;
TOTAL = length(A)*length(ALPHA);
for i3=1:length(A)
    for i2=1:length(ALPHA)
        % Sweep over SIGTAU        
        tic;
        parfor i1=1:length(SIGTAU)
            [singstrat,outcome] = singstrat_fastinfo(A(i3),Emin,Emax,d,ALPHA(i2),beta,gamma,SIGTAU(i1)*tau,tau,res1);
            
            NUM_OUTCOMES_SIGTAU_FAST(i1,i2,i3) = length(singstrat);
            SINGSTRAT_SIGTAU_FAST1(i1,i2,i3) = singstrat(1);
            OUTCOME_SIGTAU_FAST1(i1,i2,i3) = outcome(1);
            if(NUM_OUTCOMES_SIGTAU_FAST(i1,i2,i3)>1)
                SINGSTRAT_SIGTAU_FAST2(i1,i2,i3) = singstrat(2);
                OUTCOME_SIGTAU_FAST2(i1,i2,i3) = outcome(2);
                if(NUM_OUTCOMES_SIGTAU_FAST(i1,i2,i3)>2)
                    SINGSTRAT_SIGTAU_FAST3(i1,i2,i3) = singstrat(3);
                    OUTCOME_SIGTAU_FAST3(i1,i2,i3) = outcome(3);
                    if(NUM_OUTCOMES_SIGTAU_FAST(i1,i2,i3)>3)
                        SINGSTRAT_SIGTAU_FAST4(i1,i2,i3) = singstrat(4);
                        OUTCOME_SIGTAU_FAST4(i1,i2,i3) = outcome(4);
                        if(NUM_OUTCOMES_SIGTAU_FAST(i1,i2,i3)>4)
                            SINGSTRAT_SIGTAU_FAST5(i1,i2,i3) = singstrat(5);
                            OUTCOME_SIGTAU_FAST5(i1,i2,i3) = outcome(5);
                        end
                    end
                end
            end
        end
        toc;
        COUNT=COUNT+1;
        PROGRESS = COUNT/TOTAL
    end
end

% Combine results
SINGSTRAT_SIGTAU_FAST = cat(4,SINGSTRAT_SIGTAU_FAST1,SINGSTRAT_SIGTAU_FAST2,SINGSTRAT_SIGTAU_FAST3,SINGSTRAT_SIGTAU_FAST4,SINGSTRAT_SIGTAU_FAST5);
OUTCOME_SIGTAU_FAST = cat(4,OUTCOME_SIGTAU_FAST1,OUTCOME_SIGTAU_FAST2,OUTCOME_SIGTAU_FAST3,OUTCOME_SIGTAU_FAST4,OUTCOME_SIGTAU_FAST5);

clear i1 i2 i3 i4 singstrat outcome tau ans COUNT TOTAL PROGRESS
clear SINGSTRAT_SIGTAU_FAST1 SINGSTRAT_SIGTAU_FAST2 SINGSTRAT_SIGTAU_FAST3 SINGSTRAT_SIGTAU_FAST4 SINGSTRAT_SIGTAU_FAST5 OUTCOME_SIGTAU_FAST1 OUTCOME_SIGTAU_FAST2 OUTCOME_SIGTAU_FAST3 OUTCOME_SIGTAU_FAST4 OUTCOME_SIGTAU_FAST5
save('figS1.mat')
```
