## Supplementary material for "Social Information use Shapes the Coevolution of Sociality and Virulence": Source code: fitness_grad_sign_fastinfo.m

```
function [w_1,ES] = fitness_grad_sign_fastinfo(a,E,d,alpha,beta,gamma,sigma,tau)

% This function returns the parameterised fitness gradient over E array

% Assume disease and information present to start
w_1 = sign((-(alpha.*(a.*d + alpha + gamma).*tau + sigma.*d.*beta.*(-1 + a)).*beta.*tau.^2.*E.^8 + (alpha.*(a.*d + alpha + gamma).^2.*tau.^2 + 2.*d.*beta.*alpha.*sigma.*(-1 + a).*tau + d.*beta.^2.*sigma.^2.*(-1 + a)).*tau.*E.^6 - 3.*d.*sigma.*(-1 + a).*((a.*d + alpha + gamma).*tau + beta.*sigma./3).*alpha.*tau.*E.^4 + 3.*d.*sigma.^2.*(-1 + a).*(alpha./3 + (a - 2./3).*d + gamma./3).*alpha.*tau.*E.^2 - d.^2.*alpha.*sigma.^3.*(-1 + a).^2)./(beta.*E.^7.*(tau.*beta.*(a.*d + alpha).*E.^4 + (-alpha.*(a.*d + alpha + gamma).*tau - sigma.*d.*beta.*(-1 + a)).*E.^2 + sigma.*d.*alpha.*(-1 + a)).*tau.^2));
ES = sign((2.*(a.*d + alpha).*(alpha.*(a.*d + alpha + gamma).*tau + sigma.*d.*beta.*(-1 + a)).*beta.^2.*tau.^5.*E.^16 - 4.*(a.*d + alpha).*beta.*(alpha.*(a.*d + alpha + gamma).^2.*tau.^2 + (3.*d.*beta.*alpha.*sigma.*(-1 + a).*tau)./2 + d.*beta.^2.*sigma.^2.*(-1 + a)).*tau.^4.*E.^14 + 2.*(a.*d + alpha).*(alpha.*(a.*d + alpha + gamma).^3.*tau.^3 + 8.*d.*beta.*alpha.*sigma.*(-1 + a).*(a.*d + alpha + gamma).*tau.^2 + 3.*d.*beta.^2.*alpha.*sigma.^2.*(-1 + a).*tau + d.*beta.^3.*sigma.^3.*(-1 + a)).*tau.^3.*E.^12 - 12.*d.*((a.*d + alpha + gamma./6).*(a.*d + alpha + gamma).^2.*tau.^2 + 2.*sigma.*beta.*(alpha.^2./3 + (((5.*a)./4 - 2./3).*d + gamma./6).*alpha + (a.^2 - 3./4.*a).*d.^2 + gamma.*(a - 2./3).*d./4 - gamma.^2./6).*tau + sigma.^2.*(2.*alpha + (a + 1).*d + gamma).*beta.^2./6).*sigma.*(-1 + a).*alpha.*tau.^3.*E.^10 + 30.*d.*sigma.^2.*(-1 + a).*((a.*d + alpha + gamma).*(alpha.^2./5 + (((17.*a)./15 - 4./5).*d + gamma./5).*alpha + d.*((a.^2 - 13./15.*a).*d + (7.*gamma.*(a - 5./7))./15)).*tau + (8.*sigma.*beta.*(alpha.^2./4 + (((3.*a)./4 - 3./8).*d + gamma./4).*alpha + d.*((a.^2 - 5./4.*a + 3./8).*d - gamma.*(a - 3./2)./4)))./15).*alpha.*tau.^3.*E.^8 - 40.*d.*sigma.^3.*(-1 + a).*alpha.*((alpha.^3./20 + ((a./2 - 2./5).*d + gamma./10).*alpha.^2 + ((7./5.*a.^2 - 19./10.*a + 11./20).*d.^2 + (3.*gamma.*(-5./6 + a).*d)./5 + gamma.^2./20).*alpha + d.*(-1 + a).*((a.^2 - 3./5.*a).*d.^2 + (9.*gamma.*(a - 4./9).*d)./10 + gamma.^2./10)).*tau + d.*sigma.*beta.*(-1 + a).*(alpha./2 + (a - 1./2).*d - gamma./2)./10).*tau.^2.*E.^6 + 30.*d.^2.*sigma.^4.*(-1 + a).^2.*((2.*alpha.^2)./15 + (((4.*a)./5 - 3./5).*d + (2.*gamma)./15).*alpha + d.*((a.^2 - 6./5.*a + 4./15).*d + (7.*gamma.*(a - 6./7))./15)).*alpha.*tau.^2.*E.^4 - 12.*d.^3.*sigma.^5.*(-1 + a).^3.*(alpha./3 + (a - 2./3).*d + gamma./6).*alpha.*tau.*E.^2 + 2.*d.^4.*alpha.*sigma.^6.*(-1 + a).^4)./(beta.*E.^12.*(tau.*beta.*(a.*d + alpha).*E.^4 + (-alpha.*(a.*d + alpha + gamma).*tau - sigma.*d.*beta.*(-1 + a)).*E.^2 + sigma.*d.*alpha.*(-1 + a)).^2.*tau.^4));

% Special cases
GEE = max(0,1-(sigma./(tau.*E.*E)));
R0_I=(tau.*E.^2)/sigma;
R0_D=(beta.*E.^2)./(d.*(1-(1-a).*GEE)+alpha+gamma);

disonly = R0_I<1 & R0_D>1;
infoonly = R0_I>1 & R0_D<1;
neither = R0_I<=1 & R0_D<=1;

if(alpha>0)
    w_1(disonly) = -1;
    ES(disonly) = -1;
else
    w_1(disonly) = 0;
    ES(disonly) = 0;
end

if(a<1)
    w_1(infoonly) = 1;
    ES(infoonly) = 1;
else
    w_1(infoonly) = 0;
    ES(infoonly) = 0;
end 

w_1(neither) = 0;
ES(neither) = 0;
```
