## Supplementary material for "Social Information use Shapes the Coevolution of Sociality and Virulence": Source code: fitness_grad_sign_fastinfo_coevo_approx.m

```
function [w_1,ES] = fitness_grad_sign_fastinfo_coevo_approx(a,E,d,kappa,gamma,sigma,tau)

% This function returns the parameterised fitness gradient over E array

% Assume disease and information present to start
alpha = (E.^2.*(a.*d + gamma).*tau - d.*sigma.*(-1 + a))./(E.^2.*tau);
beta = kappa.*sqrt(alpha);
w_1 = sign((-alpha.^(3./2).*E.^2.*tau + (-E.^2.*(a.*d + gamma).*tau + d.*sigma.*(-1 + a)).*sqrt(alpha) + E.^2.*kappa.*(tau.*(a.*d + alpha).*E.^2 - d.*sigma.*(-1 + a))).*(-alpha.^(3./2).*E.^8.*kappa.*tau.^3 - (tau.^2.*(a.*d + gamma).*E.^4 - 2.*d.*tau.*sigma.*(-1 + a).*E.^2 + d.*sigma.^2.*(-1 + a)).*tau.*E.^4.*kappa.*sqrt(alpha) - d.*tau.^2.*kappa.^2.*sigma.*(-1 + a).*E.^8 + tau.*((a.*d + alpha + gamma).^2.*tau.^2 + d.*kappa.^2.*sigma.^2.*(-1 + a)).*E.^6 - 3.*d.*tau.^2.*sigma.*(-1 + a).*(a.*d + alpha + gamma).*E.^4 + 3.*sigma.^2.*tau.*((a - 2./3).*d + alpha./3 + gamma./3).*(-1 + a).*d.*E.^2 - d.^2.*sigma.^3.*(-1 + a).^2).*alpha./(tau.^2.*E.^7.*kappa.*(alpha.^(3./2).*E.^4.*kappa.*tau + E.^2.*kappa.*d.*(a.*E.^2.*tau - sigma.*(-1 + a)).*sqrt(alpha) - (E.^2.*(a.*d + alpha + gamma).*tau - d.*sigma.*(-1 + a)).*alpha).^2));
ES = sign(2.*(-alpha.^(3./2).*E.^2.*tau + (-E.^2.*(a.*d + gamma).*tau + d.*sigma.*(-1 + a)).*sqrt(alpha) + E.^2.*kappa.*(tau.*(a.*d + alpha).*E.^2 - d.*sigma.*(-1 + a))).*alpha.*(-6.*(-d.*tau.^3.*kappa.^2.*sigma.*(-1 + a).*E.^10./6 + tau.^2.*((a.*d + gamma./3).*(a.*d + gamma).*tau.^2 + d.*kappa.^2.*sigma.^2.*(-1 + a)./3).*E.^8 - (8.*sigma.*((a.*d + gamma./2).*tau.^2 + kappa.^2.*sigma.^2./16).*tau.*(-1 + a).*d.*E.^6)./3 + (5.*sigma.^2.*((a - 8./15).*d + (2.*gamma)./15).*tau.^2.*(-1 + a).*d.*E.^4)./2 - sigma.^3.*((a - 1./2).*d + gamma./3).*tau.*(-1 + a).*d.*E.^2 + d.^2.*sigma.^4.*(-1 + a).^2./6).*tau.^2.*E.^6.*kappa.*alpha.^(3./2) - 6.*tau.^3.*E.^8.*kappa.*(tau.^3.*(a.*d + (2.*gamma)./3).*E.^6 - (4.*d.*tau.^2.*sigma.*(-1 + a).*E.^4)./3 + (2.*d.*tau.*sigma.^2.*(-1 + a).*E.^2)./3 - d.*sigma.^3.*(-1 + a)./3).*alpha.^(5./2) - 2.*alpha.^(7./2).*E.^14.*kappa.*tau.^6 - 2.*tau.^2.*E.^6.*kappa.*(-a.*d.*tau.^3.*kappa.^2.*sigma.*(-1 + a).*E.^10./2 + a.*tau.^2.*((a.*d + gamma).^2.*tau.^2 + d.*kappa.^2.*sigma.^2.*(-1 + a)).*E.^8 - 4.*a.*sigma.*tau.*(tau.^2.*(a.*d + gamma) + kappa.^2.*sigma.^2./8).*(-1 + a).*d.*E.^6 + 6.*sigma.^2.*tau.^2.*(-1 + a).*((a.^2 - 3./4.*a).*d.^2 + gamma.*(a - 2./3).*d./4 - gamma.^2./6).*E.^4 - 4.*((a.^2 - 5./4.*a + 3./8).*d - gamma.*(a - 3./2)./4).*sigma.^3.*tau.*(-1 + a).*d.*E.^2 + sigma.^4.*(-1 + a).^2.*d.*((a - 1./2).*d - gamma./2)).*d.*sqrt(alpha) + tau.^6.*alpha.*kappa.^2.*(a.*d + alpha).*(a.*d + alpha + gamma).*E.^16 - 3.*d.*tau.^5.*alpha.*kappa.^2.*sigma.*(-1 + a).*(a.*d + alpha).*E.^14 + ((a.*d + alpha + gamma).^3.*tau.^2 + 3.*d.*alpha.*kappa.^2.*sigma.^2.*(-1 + a)).*tau.^4.*(a.*d + alpha).*E.^12 - 6.*sigma.*tau.^3.*((a.*d + alpha + gamma).^2.*(a.*d + alpha + gamma./6).*tau.^2 + sigma.^2.*kappa.^2.*alpha.*((a + 1).*d + 2.*alpha + gamma)./6).*(-1 + a).*d.*E.^10 + 15.*sigma.^2.*(a.*d + alpha + gamma).*tau.^4.*((a.^2 - 13./15.*a).*d.^2 + (((17.*alpha)./15 + (7.*gamma)./15).*a - (4.*alpha)./5 - gamma./3).*d + alpha.*(alpha + gamma)./5).*(-1 + a).*d.*E.^8 - 20.*sigma.^3.*tau.^3.*((a.^3 - 8./5.*a.^2 + 3./5.*a).*d.^3 + (((7.*alpha)./5 + (9.*gamma)./10).*a.^2 + (-(19.*alpha)./10 - (13.*gamma)./10).*a + (11.*alpha)./20 + (2.*gamma)./5).*d.^2 + ((alpha + gamma).*((alpha + gamma./5).*a - (4.*alpha)./5 - gamma./5).*d)./2 + alpha.*(alpha + gamma).^2./20).*(-1 + a).*d.*E.^6 + 15.*sigma.^4.*((a.^2 - 6./5.*a + 4./15).*d.^2 + (((4.*alpha)./5 + (7.*gamma)./15).*a - (3.*alpha)./5 - (2.*gamma)./5).*d + (2.*alpha.*(alpha + gamma))./15).*tau.^2.*(-1 + a).^2.*d.^2.*E.^4 - 6.*sigma.^5.*tau.*((a - 2./3).*d + alpha./3 + gamma./6).*(-1 + a).^3.*d.^3.*E.^2 + d.^4.*sigma.^6.*(-1 + a).^4)./(tau.^4.*E.^12.*kappa.*(alpha.^(3./2).*E.^4.*kappa.*tau + E.^2.*kappa.*d.*(a.*E.^2.*tau - sigma.*(-1 + a)).*sqrt(alpha) - (E.^2.*(a.*d + alpha + gamma).*tau - d.*sigma.*(-1 + a)).*alpha).^3));

% Special cases
GEE = max(0,1-(sigma./(tau.*E.*E)));
R0_I=(tau.*E.^2)/sigma;
R0_D=(beta.*E.^2)./(d.*(1-(1-a).*GEE)+alpha+gamma);

disonly = R0_I<1 & R0_D>1;
infoonly = R0_I>1 & R0_D<1;
neither = R0_I<=1 & R0_D<=1;

w_1(disonly) = -1;
ES(disonly) = -1;

if(a<1)
    w_1(infoonly) = 1;
    ES(infoonly) = 1;
else
    w_1(infoonly) = 0;
    ES(infoonly) = 0;
end

w_1(neither) = 0;
ES(neither) = 0;
```
