## Supplementary material for "Social Information use Shapes the Coevolution of Sociality and Virulence": Source code: fitness_grad_sign_slowinfo.m

```
function [w_1,ES] = fitness_grad_sign_slowinfo(t_max,a,b,E,d,q,alpha,beta,gamma,sigma,tau)

% This function returns the parameterised fitness gradient over E array

eqtol = 1e-2;
init_pop = ((b-d)/(4*q)*ones(1,4));

% Find basic reproductive ratios
R0_I_DF = (tau.*E.^2)/(a*d+sigma);
R0_D_IF = (beta.*E.^2)/(d+alpha+gamma);

SP = (d + alpha + gamma).*(alpha.^2 + (-beta.*E.^2 + d + gamma).*alpha + E.^2.*beta.*(b - d))./(beta.^2.*E.^4.*q);
IP = max(0,(alpha.^2 + (-beta.*E.^2 + d + gamma).*alpha + E.^2.*beta.*(b - d)).*(beta.*E.^2 - alpha - d - gamma)./(beta.^2.*E.^4.*q));
list = SP<=0 | IP<=0;
SP(list) = (b-d)/q;
IP(list) = 0;
N = SP+IP;
R0_I = tau.*E.^2.*(((a.*d + gamma + sigma).*IP + SP.*(a.*d + alpha + gamma + sigma)).*N + E.^2.*IP.*beta.*(IP + SP))./(N.*((a.*d + sigma).*(a.*d + alpha + gamma + sigma).*N + E.^2.*IP.*beta.*(a.*d + alpha + sigma)));
R0_I(IP==0) = R0_I_DF(IP==0);

SP = (a.*d + sigma).*((a.^2 - a).*d.^2 + ((-E.^2.*tau + sigma).*a - sigma).*d + b.*E.^2.*tau)./(E.^4.*q.*tau.^2);
SG = max(0,-(-E.^2.*tau + a.*d + sigma).*((a.^2 - a).*d.^2 + ((-E.^2.*tau + sigma).*a - sigma).*d + b.*E.^2.*tau)./(E.^4.*q.*tau.^2));
list = SP<=0 | SG<=0;
SP(list) = (b-d)/q;
SG(list) = 0;
N = SP+SG;
R0_D = beta.*(((sigma + d + alpha + gamma).*SG + SP.*(a.*d + alpha + gamma + sigma)).*N + E.^2.*SG.*tau.*(SP + SG)).*E.^2./(((d + alpha + gamma).*(a.*d + alpha + gamma + sigma).*N + E.^2.*SG.*tau.*(a.*d + alpha + gamma)).*N);
R0_D(SG==0) = R0_D_IF(SG==0);

% Solve ODEs to find equilibrium
SP = zeros(length(E),1);
SG = zeros(length(E),1);
IP = zeros(length(E),1);
IG = zeros(length(E),1);
N = zeros(length(E),1);
for i=1:length(E)
    [~,x] = sociality_slowinfo_mex(t_max,a,b,E(i),d,q,alpha,beta,gamma,sigma,tau,eqtol,init_pop);
    
    SP(i) = x(end,1);
    SG(i) = x(end,2);
    IP(i) = x(end,3);
    IG(i) = x(end,4);
    N(i) = SP(i)+SG(i)+IP(i)+IG(i);
end

% Assume disease and information present to start
w_1 = sign(N.*E.*(N.*q - b).*((a.*d + alpha + gamma + sigma).^2.*(d.*(SG + IG).*(a - 1).*(d + alpha + gamma).*tau + beta.*alpha.*(IP + IG).*(a.*d + sigma)).*(a.*d + sigma).*(d + alpha + gamma).*N.^4 + 2.*(a.*d + alpha + gamma + sigma).*(d.*(SG + IG).^2.*(a - 1).*(a.*d + alpha + gamma).*tau.^2 + (IP + IG).*((a.^2 - 1).*d.^2 + ((2.*alpha + 2.*gamma + sigma).*a - 2.*gamma - sigma).*d + alpha.*(alpha + gamma + 2.*sigma)).*(SG + IG).*beta.*tau + beta.^2.*alpha.*(IP + IG).^2.*(a.*d + alpha + sigma)).*E.^2.*(a.*d + sigma).*(d + alpha + gamma).*N.^3 + (d.*(SG + IG).^3.*(a - 1).*(a.*d + alpha + gamma).^2.*(a.*d + sigma).*tau.^3 + (IP + IG).*((2.*a.^4 + 2.*a.^3 - 4.*a.^2).*d.^4 + 7.*((alpha + (8.*gamma)./7 + (4.*sigma)./7).*a.^2 + ((2.*alpha)./7 - (4.*gamma)./7 + (2.*sigma)./7).*a - (3.*alpha)./7 - (4.*gamma)./7 - (6.*sigma)./7).*a.*d.^3 + ((9.*alpha.^2 + (15.*gamma + 12.*sigma).*alpha + 6.*gamma.^2 + 12.*gamma.*sigma + 2.*sigma.^2).*a.^2 + (2.*alpha.^2 + (-4.*gamma + 2.*sigma).*alpha - 6.*gamma.^2 - 8.*gamma.*sigma).*a + alpha.^2 + (gamma - 2.*sigma).*alpha - 4.*gamma.*sigma - 2.*sigma.^2).*d.^2 + ((5.*alpha.^3 + (10.*gamma + 14.*sigma).*alpha.^2 + (5.*gamma.^2 + 20.*gamma.*sigma + 5.*sigma.^2).*alpha + 6.*gamma.^2.*sigma + 4.*gamma.*sigma.^2).*a + 2.*alpha.^3 + (4.*gamma + 4.*sigma).*alpha.^2 + (2.*gamma.^2 - 2.*gamma.*sigma + sigma.^2).*alpha - 6.*gamma.^2.*sigma - 4.*gamma.*sigma.^2).*d + alpha.*(alpha + gamma).*(alpha.^2 + (2.*gamma + 6.*sigma).*alpha + gamma.^2 + 6.*gamma.*sigma + 6.*sigma.^2)).*(SG + IG).^2.*beta.*tau.^2 + 2.*(IP + IG).^2.*(a.*(a - 1).*(a.^2 + 4.*a + 1).*d.^4./2 + (((3.*sigma)./2 + (5.*alpha)./2 + 3.*gamma).*a.^3 + ((5.*sigma)./2 + (5.*alpha)./2).*a.^2 + (-(3.*alpha)./2 - 3.*gamma - (7.*sigma)./2).*a - sigma./2 - alpha./2).*d.^3 + (((9.*alpha.^2)./2 + (7.*gamma + 5.*sigma).*alpha + 3.*gamma.^2 + 6.*gamma.*sigma + (3.*sigma.^2)./2).*a.^2 + ((5.*alpha.^2)./2 + (-gamma + 3.*sigma).*alpha + sigma.^2./2 - 3.*gamma.^2 - 3.*gamma.*sigma).*a + (-gamma - 2.*sigma).*alpha - 3.*gamma.*sigma - 2.*sigma.^2).*d.^2 + (((7.*alpha.^3)./2 + (6.*gamma + (15.*sigma)./2).*alpha.^2 + (10.*gamma.*sigma + 5./2.*gamma.^2 + 5./2.*sigma.^2).*alpha + 3.*gamma.^2.*sigma + 3.*gamma.*sigma.^2 + sigma.^3./2).*a + (3.*alpha.^3)./2 + (gamma + (5.*sigma)./2).*alpha.^2 + (-1./2.*gamma.^2 + 1./2.*sigma.^2 - 2.*gamma.*sigma).*alpha - sigma.^3./2 - 3.*gamma.^2.*sigma - 3.*gamma.*sigma.^2).*d + alpha.*(alpha + gamma).*(alpha.^2 + (gamma + 4.*sigma).*alpha + 2.*gamma.*sigma + 3.*sigma.^2)).*(SG + IG).*beta.^2.*tau + alpha.*beta.^3.*(IP + IG).^3.*(a.*d + alpha + sigma).^2.*(d + alpha + gamma)).*E.^4.*N.^2 + 2.*(IP + IG).*(beta.*(IP + IG) + tau.*(SG + IG)).*tau.*E.^6.*(SG + IG).*beta.*((a.^2.*(a - 1).*d.^3 + 2.*a.*((alpha + gamma./2 + sigma./2).*a - gamma./2 - sigma./2).*d.^2 + ((2.*alpha.^2 + (2.*gamma + 2.*sigma).*alpha + gamma.*sigma).*a + alpha.^2 + alpha.*gamma - gamma.*sigma).*d + alpha.*(alpha + gamma).*(alpha + gamma + 2.*sigma)).*(SG + IG).*tau + (IP + IG).*((a.^3 - a).*d.^3 + ((2.*alpha + 2.*gamma + 2.*sigma).*a.^2 + (-2.*gamma - sigma).*a - alpha - sigma).*d.^2 + ((2.*alpha.^2 + (2.*gamma + 2.*sigma).*alpha + 2.*gamma.*sigma + sigma.^2).*a + (-gamma - sigma).*alpha - 2.*gamma.*sigma - sigma.^2).*d + alpha.*(alpha + sigma).*(alpha + gamma)).*beta).*N + (IP + IG).*(beta.*(IP + IG) + tau.*(SG + IG)).^2.*tau.*(alpha.*(SG + IG).*(a.*d + alpha + gamma).*tau + beta.*d.*(IP + IG).*(a - 1).*(a.*d + alpha + sigma)).*E.^8.*(SG + IG).*beta)./(N.^3.*d.*(a.*d + sigma).*(d + alpha + gamma).*(a.*d + alpha + gamma + sigma) + ((2.*a.^2.*d.^2 + a.*((alpha + gamma).*a + 2.*alpha + 2.*gamma + 2.*sigma).*d + (alpha + gamma).*((alpha + gamma + sigma).*a + sigma)).*d.*(SG + IG).*tau + (IP + IG).*beta.*((a.^2 + a).*d.^3 + (alpha.*a.^2 + (2.*alpha + 2.*gamma + 2.*sigma).*a + alpha + sigma).*d.^2 + (alpha.*(alpha + gamma + 2.*sigma).*a + alpha.^2 + (gamma + 2.*sigma).*alpha + 2.*gamma.*sigma + sigma.^2).*d + alpha.*sigma.*(alpha + gamma + sigma))).*E.^2.*N.^2 + (a.*d.*(SG + IG).^2.*(a.*d + alpha + gamma).*tau.^2 + (IP + IG).*((a.^2 + 2.*a).*d.^2 + ((2.*alpha + 2.*gamma + sigma).*a + 2.*alpha + sigma).*d + alpha.*(alpha + gamma + 2.*sigma)).*(SG + IG).*beta.*tau + beta.^2.*(IP + IG).^2.*(alpha + d).*(a.*d + alpha + sigma)).*E.^4.*N + (IP + IG).*beta.*E.^6.*(SG + IG).*tau.*(beta.*(IP + IG) + tau.*(SG + IG)).*(a.*d + alpha)).^2);
ES = sign(-2.*((tau.^2.*a.^2.*(SG + IG).^2.*(a - 1).*d.^5 + (2.*(a - 1).*a.*(SG + IG).^2.*((a + 1./2).*alpha + gamma.*a + gamma./2 + sigma./2).*tau.^2 + 3.*(IP + IG).*alpha.*(a - 1).*(SG + IG).*beta.*(a.^2 + 2./3.*a + 1./3).*tau + beta.^2.*alpha.*a.^3.*(IP + IG).^2).*d.^4 + ((alpha + gamma).*(a - 1).*a.*(SG + IG).^2.*((a + 2).*alpha + gamma.*a + 2.*gamma + 2.*sigma).*tau.^2 + 2.*(IP + IG).*alpha.*(a - 1).*((a + 1).^2.*alpha + gamma.*a.^2 + (2.*gamma + 3.*sigma).*a + gamma + sigma).*(SG + IG).*beta.*tau + (IP + IG).^2.*alpha.*((a + 1).*alpha + gamma + 3.*sigma).*a.^2.*beta.^2).*d.^3 + (a.*(SG + IG).^2.*(alpha + gamma).^2.*(a - 1).*(alpha + gamma + sigma).*tau.^2 + 2.*(IP + IG).*alpha.*(a - 1).*(SG + IG).*((a + 1./2).*alpha.^2 + ((2.*gamma + 2.*sigma).*a + gamma + 2.*sigma).*alpha + gamma.*(gamma + 2.*sigma).*a + ((gamma + 3.*sigma).*(gamma + sigma))./2).*beta.*tau + (IP + IG).^2.*alpha.*(alpha.^2.*a + ((gamma + 3.*sigma).*a + 2.*sigma).*alpha + 2.*gamma.*sigma + 3.*sigma.^2).*a.*beta.^2).*d.^2 + 2.*(IP + IG).*alpha.*((SG + IG).*(a - 1).*(alpha + gamma).*(alpha + gamma + sigma).*tau + (IP + IG).*(alpha.^2.*a + ((gamma + (3.*sigma)./2).*a + sigma./2).*alpha + sigma.*(gamma + sigma)./2).*beta).*beta.*sigma.*d + beta.^2.*alpha.^2.*sigma.^2.*(IP + IG).^2.*(alpha + gamma + sigma)).*(a.*d + alpha + gamma + sigma).^2.*(a.*d + sigma).*(d + alpha + gamma).*N.^6 + 3.*(a.*d + alpha + gamma + sigma).^2.*E.^2.*(a.*d + sigma).*((tau.*a.*(SG + IG) + (IP + IG).*(a + 1).*beta).*tau.^2.*(a - 1).*a.*(SG + IG).^2.*d.^4 + (a.*(SG + IG).^3.*(a - 1).*(a + 1).*(alpha + gamma).*tau.^3 + 2.*(IP + IG).*(a - 1).*((a + 1).^2.*alpha + a.*(a.*gamma + 3.*gamma + sigma)./2).*(SG + IG).^2.*beta.*tau.^2 + beta.^2.*alpha.*(IP + IG).^2.*(a.^3 + 3.*a.^2 - a - 1).*(SG + IG).*tau + beta.^3.*alpha.*a.^2.*(IP + IG).^3).*d.^3 + (a.*(SG + IG).^3.*(alpha + gamma).^2.*(a - 1).*tau.^3 + 3.*(IP + IG).*(a - 1).*(SG + IG).^2.*beta.*((a + 1).*alpha.^2 + (((5.*gamma)./3 + (2.*sigma)./3).*a + gamma + sigma).*alpha + 2.*(gamma + sigma./2).*gamma.*a./3).*tau.^2 + 3.*(IP + IG).^2.*alpha.*((a.^2 + 2./3.*a - 2./3).*alpha + (gamma + (2.*sigma)./3).*a.^2 + (-gamma./3 + (4.*sigma)./3).*a - gamma./3 - (2.*sigma)./3).*(SG + IG).*beta.^2.*tau + (IP + IG).^3.*alpha.*a.*beta.^3.*((a + 1).*alpha + 2.*sigma)).*d.^2 + (IP + IG).*alpha.*((SG + IG).^2.*(a - 1).*(alpha + gamma).*(alpha + gamma + 3.*sigma).*tau.^2 + 2.*(IP + IG).*((a - 1./2).*alpha.^2 + ((gamma + (5.*sigma)./2).*a - gamma./2).*alpha + 3.*((gamma + sigma./3).*a - (2.*gamma)./3 + sigma./3).*sigma./2).*(SG + IG).*beta.*tau + (IP + IG).^2.*(alpha.^2.*a + (2.*a.*sigma + sigma).*alpha + sigma.^2).*beta.^2).*beta.*d + (IP + IG).^2.*((IP + IG).*(alpha + sigma).*beta + tau.*(SG + IG).*(alpha + gamma + 2.*sigma)).*alpha.^2.*beta.^2.*sigma).*(d + alpha + gamma).*N.^5 + 3.*(((SG + IG).^2.*a.^2.*tau.^2 + 2.*(IP + IG).*(a + 3./2).*a.*(SG + IG).*beta.*tau + beta.^2.*(IP + IG).^2.*(a.^2 + 3.*a + 1)).*tau.^2.*(a - 1).*a.*(SG + IG).^2.*d.^4 + (2.*a.^2.*(SG + IG).^4.*(a - 1).*(alpha + gamma).*tau.^4 + 7.*(IP + IG).*(a - 1).*a.*(SG + IG).^3.*beta.*((a + 1).*alpha + (gamma + (2.*sigma)./7).*a + (3.*gamma)./7 + sigma./7).*tau.^3 + 8.*(IP + IG).^2.*((a.^3 + 1./2.*a.^2 - 3./4.*a - 1./8).*alpha + 5.*((gamma + (2.*sigma)./5).*a + gamma + (3.*sigma)./5).*(a - 1).*a./8).*(SG + IG).^2.*beta.^2.*tau.^2 + 3.*(IP + IG).^3.*alpha.*a.*(SG + IG).*beta.^3.*(a.^2 + a - 1./3).*tau + beta.^4.*alpha.*a.^2.*(IP + IG).^4).*d.^3 + (a.*(SG + IG).^4.*(alpha + gamma).^2.*(a - 1).*tau.^4 + 8.*(IP + IG).*(a - 1).*(SG + IG).^3.*beta.*((a + 1./2).*alpha.^2 + (((13.*gamma)./8 + (7.*sigma)./8).*a + gamma./2 + sigma./4).*alpha + (5.*gamma.*a.*(gamma + (3.*sigma)./5))./8).*tau.^3 + 14.*(IP + IG).^2.*((a.^2 - 2./7).*alpha.^2 + (((8.*gamma)./7 + (13.*sigma)./14).*a.^2 + (-(2.*gamma)./7 + sigma./7).*a - gamma./2 - (5.*sigma)./14).*alpha + 5.*(a - 1).*a.*(gamma.^2 + gamma.*sigma + 1./5.*sigma.^2)./14).*(SG + IG).^2.*beta.^2.*tau.^2 + 8.*(IP + IG).^3.*alpha.*((a.^2 + 5./8.*a - 1./8).*alpha + ((5.*gamma)./8 + (3.*sigma)./4).*a.^2 + (-gamma./8 + (5.*sigma)./8).*a - gamma./8 - sigma./8).*(SG + IG).*beta.^3.*tau + (IP + IG).^4.*alpha.*((a + 2).*alpha + 2.*sigma).*a.*beta.^4).*d.^2 + 3.*(IP + IG).*alpha.*((alpha + gamma).*(a - 1).*(alpha + gamma + (5.*sigma)./3).*(SG + IG).^3.*tau.^3 + 8.*(IP + IG).*((a - 1./4).*alpha.^2 + (((13.*gamma)./8 + 2.*sigma).*a - (3.*gamma)./4 - sigma./8).*alpha + (5./8.*gamma.^2 + 7./4.*gamma.*sigma + 5./8.*sigma.^2).*a - ((gamma + (9.*sigma)./4).*gamma)./2).*(SG + IG).^2.*beta.*tau.^2./3 + (7.*(IP + IG).^2.*((a + 2./7).*alpha.^2 + ((gamma + (13.*sigma)./7).*a - (2.*gamma)./7 + (4.*sigma)./7).*alpha + 5.*((gamma + (3.*sigma)./5).*a - (2.*gamma)./5 + (2.*sigma)./5).*sigma./7).*(SG + IG).*beta.^2.*tau)./3 + (2.*(IP + IG).^3.*((a + 1./2).*alpha + sigma./2).*(alpha + sigma).*beta.^3)./3).*beta.*d + (IP + IG).^2.*alpha.^2.*((SG + IG).^2.*(alpha.^2 + (2.*gamma + 5.*sigma).*alpha + gamma.^2 + 5.*gamma.*sigma + 5.*sigma.^2).*tau.^2 + 2.*(IP + IG).*(SG + IG).*beta.*(alpha.^2 + (gamma + (7.*sigma)./2).*alpha + (3.*sigma.*(gamma + (5.*sigma)./3))./2).*tau + beta.^2.*(IP + IG).^2.*(alpha + sigma).^2).*beta.^2).*(a.*d + alpha + gamma + sigma).*E.^4.*(a.*d + sigma).*(d + alpha + gamma).*N.^4 + ((tau.*a.*(SG + IG) + (IP + IG).*(a + 1).*beta).*tau.^2.*(a - 1).*((SG + IG).^2.*a.^2.*tau.^2 + 2.*a.*beta.*(SG + IG).*(IP + IG).*(a + 4).*tau + beta.^2.*(IP + IG).^2.*(a.^2 + 8.*a + 1)).*a.^2.*(SG + IG).^2.*d.^6 + 3.*((a - 1).*(alpha + gamma + sigma./3).*a.^3.*(SG + IG).^5.*tau.^5 + 17.*(IP + IG).*(a - 1).*((a + 28./17).*alpha + ((18.*gamma)./17 + (6.*sigma)./17).*a + (18.*gamma)./17 + (15.*sigma)./17).*a.^2.*(SG + IG).^4.*beta.*tau.^4./3 + (25.*(IP + IG).^2.*((a.^3 + 48./25.*a.^2 - 36./25.*a - 17./25).*alpha + 27.*((gamma + sigma./3).*a.^2 + (2.*gamma + (14.*sigma)./9).*a + gamma./3 + (5.*sigma)./9).*(a - 1)./25).*a.*(SG + IG).^3.*beta.^2.*tau.^3)./3 + (11.*(IP + IG).^3.*(SG + IG).^2.*beta.^3.*((a.^4 + 43./11.*a.^3 - 6./11.*a.^2 - 16./11.*a - 2./11).*alpha + 12.*((gamma + sigma./3).*a.^3 + (3.*gamma + (9.*sigma)./4).*a.^2 + (gamma + (3.*sigma)./2).*a + sigma./12).*(a - 1)./11).*tau.^2)./3 + 3.*(IP + IG).^4.*alpha.*(a + 1./3).*a.^2.*(SG + IG).*beta.^4.*tau + beta.^5.*alpha.*a.^2.*(IP + IG).^5./3).*a.*d.^5 + (3.*a.^3.*(SG + IG).^5.*(a - 1).*(alpha + gamma).*(alpha + gamma + sigma).*tau.^5 + 34.*(IP + IG).*(a - 1).*((a + 29./34).*alpha.^2 + (((61.*gamma)./34 + (16.*sigma)./17).*a + (19.*gamma)./17 + (21.*sigma)./17).*alpha + 27.*(gamma + sigma./9).*(gamma + sigma).*a./34 + (9.*gamma.^2)./34 + (12.*gamma.*sigma)./17 + (3.*sigma.^2)./17).*a.^2.*(SG + IG).^4.*beta.*tau.^4 + 68.*(IP + IG).^2.*((a.^3 + 3./4.*a.^2 - 21./34.*a - 7./68).*alpha.^2 + (((121.*gamma)./68 + (63.*sigma)./68).*a.^3 + ((9.*gamma)./17 + (24.*sigma)./17).*a.^2 + (-(99.*gamma)./68 - (39.*sigma)./34).*a - (2.*gamma)./17 - (21.*sigma)./68).*alpha + 27.*(a - 1).*((gamma + sigma./6).*(gamma + sigma).*a.^2 + (2./3.*gamma.^2 + 5./3.*gamma.*sigma + 5./9.*sigma.^2).*a + ((gamma + (2.*sigma)./3).*sigma)./6)./34).*a.*(SG + IG).^3.*beta.^2.*tau.^3 + 46.*(IP + IG).^3.*((a.^4 + 83./46.*a.^3 - 3./23.*a.^2 - 4./23.*a - 1./46).*alpha.^2 + (((75.*gamma)./46 + (17.*sigma)./23).*a.^4 + ((33.*gamma)./23 + (55.*sigma)./23).*a.^3 + (-(30.*gamma)./23 - (27.*sigma)./46).*a.^2 + (-(15.*gamma)./46 - (13.*sigma)./23).*a - sigma./46).*alpha + 15.*(a - 1).*a.*((gamma + sigma./5).*(gamma + sigma).*a.^2 + (gamma.^2 + 12./5.*gamma.*sigma + 9./10.*sigma.^2).*a + 2.*(gamma + (3.*sigma)./4).*sigma./5)./23).*(SG + IG).^2.*beta.^3.*tau.^2 + 9.*(IP + IG).^4.*alpha.*((a.^2 + 32./9.*a + 10./9).*alpha + gamma.*a.^2 + ((14.*gamma)./9 + 3.*sigma).*a - (2.*gamma)./9 + sigma).*a.^2.*(SG + IG).*beta.^4.*tau + 2.*(IP + IG).^5.*alpha.*a.^2.*beta.^5.*((a + 3./2).*alpha + gamma.*a./2 + (3.*sigma)./2)).*d.^4 + (a.^2.*(SG + IG).^5.*(alpha + gamma).^2.*(a - 1).*(alpha + gamma + 3.*sigma).*tau.^5 + 29.*(IP + IG).*(a - 1).*a.*(SG + IG).^4.*beta.*((a + 10./29).*alpha.^3 + (((70.*gamma)./29 + (57.*sigma)./29).*a + (20.*gamma)./29 + (37.*sigma)./29).*alpha.^2 + ((53./29.*gamma.^2 + 93./29.*gamma.*sigma + 15./29.*sigma.^2).*a + (10.*gamma.^2)./29 + (46.*gamma.*sigma)./29 + (15.*sigma.^2)./29).*alpha + 12.*((gamma.^2 + 3.*gamma.*sigma + sigma.^2).*a + 3.*(gamma + (2.*sigma)./3).*sigma./4).*gamma./29).*tau.^4 + 82.*(IP + IG).^2.*((1./82 - 6./41.*a + a.^3 + 21./82.*a.^2).*alpha.^3 + (((193.*gamma)./82 + (78.*sigma)./41).*a.^3 + (-(6.*gamma)./41 + (45.*sigma)./41).*a.^2 + (-(24.*gamma)./41 - (30.*sigma)./41).*a - (3.*sigma)./41 + gamma./82).*alpha.^2 + ((249./82.*gamma.*sigma + 141./82.*gamma.^2 + 51./82.*sigma.^2).*a.^3 + (15./82.*gamma.*sigma - 63./82.*gamma.^2 + 30./41.*sigma.^2).*a.^2 + (-18./41.*gamma.^2 - 45./82.*sigma.^2 - 135./82.*gamma.*sigma).*a - 9.*(gamma + (2.*sigma)./3).*sigma./82).*alpha + 15.*(a - 1).*a.*((gamma.^3 + 3.*gamma.^2.*sigma + 3./2.*gamma.*sigma.^2 + 1./10.*sigma.^3).*a + 6.*(gamma.^2 + gamma.*sigma + 1./6.*sigma.^2).*sigma./5)./41).*(SG + IG).^3.*beta.^2.*tau.^3 + 82.*(IP + IG).^3.*((3./41.*a + a.^3 + 77./82.*a.^2).*alpha.^3 + (((165.*gamma)./82 + (129.*sigma)./82).*a.^3 + ((63.*gamma)./82 + (90.*sigma)./41).*a.^2 + (-(33.*gamma)./82 - (6.*sigma)./41).*a - (9.*sigma)./82 - (3.*gamma)./82).*alpha.^2 + ((189./82.*gamma.*sigma + 51./41.*gamma.^2 + 18./41.*sigma.^2).*a.^3 + (45./41.*gamma.*sigma - 21./82.*gamma.^2 + 45./41.*sigma.^2).*a.^2 + (-18./41.*gamma.^2 - 27./82.*sigma.^2 - 105./82.*gamma.*sigma).*a - 6.*(gamma + (3.*sigma)./4).*sigma./41).*alpha + 10.*((gamma.^3 + 3.*gamma.^2.*sigma + 9./5.*gamma.*sigma.^2 + 1./5.*sigma.^3).*a + (3.*gamma.^2.*sigma)./2 + (9.*gamma.*sigma.^2)./5 + (9.*sigma.^3)./20).*(a - 1).*a./41).*(SG + IG).^2.*beta.^3.*tau.^2 + 29.*(IP + IG).^4.*alpha.*a.*(SG + IG).*beta.^4.*((a.^2 + 44./29.*a + 11./29).*alpha.^2 + (((40.*gamma)./29 + (27.*sigma)./29).*a.^2 + ((31.*gamma)./29 + (79.*sigma)./29).*a - (2.*gamma)./29 + (20.*sigma)./29).*alpha + 11.*(gamma + (27.*sigma)./11).*gamma.*a.^2./29 + (-1./29.*gamma.^2 + 31./29.*gamma.*sigma + 27./29.*sigma.^2).*a - gamma.^2./29 - (4.*gamma.*sigma)./29 + (9.*sigma.^2)./29).*tau + (IP + IG).^5.*alpha.*((a.^2 + 6.*a + 3).*alpha.^2 + (gamma.*a.^2 + (3.*gamma + 6.*sigma).*a + 6.*sigma).*alpha + 3.*gamma.*sigma.*a + 3.*sigma.^2).*a.*beta.^5).*d.^3 + (sigma.*a.*(SG + IG).^5.*(alpha + gamma).^3.*(a - 1).*tau.^5 + 9.*(IP + IG).*(alpha + gamma).*(a - 1).*(SG + IG).^4.*beta.*(alpha.^3.*a + ((2.*gamma + (40.*sigma)./9).*a + (10.*sigma)./9).*alpha.^2 + ((gamma.^2 + 52./9.*gamma.*sigma + 8./3.*sigma.^2).*a + 10.*(gamma + (4.*sigma)./5).*sigma./9).*alpha + 4.*(gamma + (3.*sigma)./4).*gamma.*a.*sigma./3).*tau.^4 + 46.*(IP + IG).^2.*((3./46 + a.^2 + 3./23.*a).*alpha.^4 + (((129.*gamma)./46 + (165.*sigma)./46).*a.^2 + ((18.*sigma)./23 - (6.*gamma)./23).*a + (3.*gamma)./23 - (9.*sigma)./46).*alpha.^3 + ((15./2.*gamma.*sigma + 60./23.*gamma.^2 + 117./46.*sigma.^2).*a.^2 + (-33./46.*gamma.*sigma - 21./23.*gamma.^2 + 51./46.*sigma.^2).*a + (3.*gamma.^2)./46 - (24.*gamma.*sigma)./23 - (9.*sigma.^2)./23).*alpha.^2 + ((13./46.*sigma.^3 + 105./23.*gamma.^2.*sigma + 78./23.*gamma.*sigma.^2 + 37./46.*gamma.^3).*a.^2 + (-99./46.*gamma.^2.*sigma + 6./23.*sigma.^3 - 12./23.*gamma.*sigma.^2 - 12./23.*gamma.^3).*a - (39.*gamma.^2.*sigma)./46 - (21.*gamma.*sigma.^2)./23 - (5.*sigma.^3)./46).*alpha + 15.*(gamma.^2 + 6./5.*gamma.*sigma + 3./10.*sigma.^2).*(a - 1).*gamma.*a.*sigma./23).*(SG + IG).^3.*beta.^2.*tau.^3 + 68.*(IP + IG).^3.*(SG + IG).^2.*((3./34 + a.^2 + 37./68.*a).*alpha.^4 + (((39.*gamma)./17 + (193.*sigma)./68).*a.^2 + ((67.*sigma)./34 + (9.*gamma)./17).*a + (3.*gamma)./68 + (3.*sigma)./34).*alpha.^3 + ((345./68.*gamma.*sigma + 117./68.*gamma.^2 + 30./17.*sigma.^2).*a.^2 + (18./17.*gamma.*sigma - 21./68.*gamma.^2 + 30./17.*sigma.^2).*a - (3.*gamma.^2)./68 - (39.*gamma.*sigma)./68 - (3.*sigma.^2)./34).*alpha.^2 + ((7./34.*sigma.^3 + 87./34.*gamma.^2.*sigma + 9./4.*gamma.*sigma.^2 + 29./68.*gamma.^3).*a.^2 + (-15./17.*gamma.^2.*sigma + 11./34.*sigma.^3 + 15./68.*gamma.*sigma.^2 - 5./17.*gamma.^3).*a - (33.*gamma.^2.*sigma)./68 - (21.*gamma.*sigma.^2)./34 - (3.*sigma.^3)./34).*alpha + 5.*(gamma.^2 + gamma.*sigma + 1./10.*sigma.^2).*(gamma + sigma./2).*(a - 1).*a.*sigma./17).*beta.^3.*tau.^2 + 34.*(IP + IG).^4.*alpha.*((a.^2 + 14./17.*a + 2./17).*alpha.^3 + (((57.*gamma)./34 + (35.*sigma)./17).*a.^2 + ((12.*gamma)./17 + (81.*sigma)./34).*a + (11.*sigma)./34).*alpha.^2 + ((12./17.*gamma.^2 + 46./17.*gamma.*sigma + 27./34.*sigma.^2).*a.^2 + (-3./34.*gamma.^2 + 24./17.*gamma.*sigma + 31./17.*sigma.^2).*a - sigma.*(gamma - 5.*sigma)./17).*alpha + 11.*(gamma.*(gamma + (27.*sigma)./22).*a.^2 + (-3./22.*gamma.^2 + 10./11.*gamma.*sigma + 9./22.*sigma.^2).*a - ((gamma + 3.*sigma).*(gamma - sigma))./22).*sigma./17).*(SG + IG).*beta.^4.*tau + 3.*(IP + IG).^5.*alpha.*((a.^2 + 2.*a + 1./3).*alpha.^2 + (gamma.*a.^2 + (gamma + 2.*sigma).*a + (2.*sigma)./3).*alpha + sigma.*(gamma.*a + sigma./3)).*(alpha + sigma).*beta.^5).*d.^2 + 9.*(IP + IG).*alpha.*((alpha + gamma).^2.*(a - 1).*(SG + IG).^4.*(alpha + gamma + (11.*sigma)./9).*sigma.*tau.^4 + 11.*(IP + IG).*(SG + IG).^3.*beta.*((3./11 + a).*alpha.^4 + (((75.*sigma)./11 + (34.*gamma)./11).*a + (9.*gamma)./11 + (9.*sigma)./11).*alpha.^3 + ((189./11.*gamma.*sigma + 36./11.*gamma.^2 + 102./11.*sigma.^2).*a + (9.*gamma.^2)./11 - (9.*gamma.*sigma)./11 + (18.*sigma.^2)./11).*alpha.^2 + ((153./11.*gamma.^2.*sigma + 29./11.*sigma.^3 + 174./11.*gamma.*sigma.^2 + 14./11.*gamma.^3).*a + (3.*gamma.^3)./11 - (45.*gamma.^2.*sigma)./11 - (24.*gamma.*sigma.^2)./11 + sigma.^3).*alpha + (((gamma.^3 + 39.*gamma.^2.*sigma + 72.*gamma.*sigma.^2 + 28.*sigma.^3).*a - 27.*gamma.^2.*sigma - 42.*gamma.*sigma.^2 - 8.*sigma.^3).*gamma)./11).*tau.^3./9 + (25.*(IP + IG).^2.*(SG + IG).^2.*beta.^2.*((8./25 + a).*alpha.^4 + (((121.*sigma)./25 + (63.*gamma)./25).*a + (3.*gamma)./5 + (41.*sigma)./25).*alpha.^3 + ((249./25.*gamma.*sigma + 51./25.*gamma.^2 + 141./25.*sigma.^2).*a + (6.*gamma.^2)./25 + (6.*gamma.*sigma)./5 + (57.*sigma.^2)./25).*alpha.^2 + ((156./25.*gamma.^2.*sigma + 37./25.*sigma.^3 + 42./5.*gamma.*sigma.^2 + 13./25.*gamma.^3).*a - gamma.^3./25 - (6.*gamma.^2.*sigma)./5 + (6.*gamma.*sigma.^2)./25 + (23.*sigma.^3)./25).*alpha + 28.*((gamma.^3 + 18./7.*gamma.^2.*sigma + 39./28.*gamma.*sigma.^2 + 1./28.*sigma.^3).*a - (19.*gamma.^3)./28 - (9.*gamma.^2.*sigma)./7 - (9.*gamma.*sigma.^2)./28 - sigma.^3./28).*sigma./25).*tau.^2)./9 + (17.*(IP + IG).^3.*(SG + IG).*beta.^3.*((a + 7./17).*alpha.^4 + (((32.*gamma)./17 + (61.*sigma)./17).*a + (7.*gamma)./17 + (29.*sigma)./17).*alpha.^3 + ((15./17.*gamma.^2 + 93./17.*gamma.*sigma + 53./17.*sigma.^2).*a + 21.*(gamma + (37.*sigma)./21).*sigma./17).*alpha.^2 + 33.*((gamma.^2 + 64./33.*gamma.*sigma + 3./11.*sigma.^2).*a - gamma.^2./11 + (17.*gamma.*sigma)./33 + (5.*sigma.^2)./11).*sigma.*alpha./17 + 11.*((gamma + (9.*sigma)./11).*a - (2.*gamma)./11 + (3.*sigma)./11).*gamma.*sigma.^2./17).*tau)./9 + (IP + IG).^4.*((a + 2./3).*alpha.^2 + (gamma.*a + 1./3.*gamma + 2./3.*sigma).*alpha + gamma.*sigma./3).*(alpha + sigma).^2.*beta.^4./3).*beta.*d + (IP + IG).^2.*((IP + IG).*(alpha + sigma).*beta + tau.*(SG + IG).*(alpha + gamma + 2.*sigma)).*alpha.^2.*(alpha + gamma).*beta.^2.*((alpha.^2 + (2.*gamma + 10.*sigma).*alpha + gamma.^2 + 10.*gamma.*sigma + 10.*sigma.^2).*(SG + IG).^2.*tau.^2 + 2.*(IP + IG).*(alpha.^2 + (gamma + 6.*sigma).*alpha + 4.*gamma.*sigma + 5.*sigma.^2).*(SG + IG).*beta.*tau + beta.^2.*(IP + IG).^2.*(alpha + sigma).^2)).*E.^6.*N.^3 + 3.*(IP + IG).*(beta.*(IP + IG) + tau.*(SG + IG)).*tau.*E.^8.*(SG + IG).*beta.*(((SG + IG).^2.*a.^2.*tau.^2 + 2.*(IP + IG).*(a + 3./2).*a.*(SG + IG).*beta.*tau + beta.^2.*(IP + IG).^2.*(a.^2 + 3.*a + 1)).*tau.*(a - 1).*a.*(SG + IG).*d.^4 + (2.*(a - 1).*(alpha + gamma + sigma./2).*a.^2.*(SG + IG).^3.*tau.^3 + 6.*(IP + IG).*((a.^2 + 1./6.*a - 1./3).*alpha + 7.*(a - 1).*((gamma + (4.*sigma)./7).*a + (3.*gamma)./7 + (4.*sigma)./7)./6).*a.*(SG + IG).^2.*beta.*tau.^2 + 4.*(IP + IG).^2.*((a.^3 + a.^2 - 1./2.*a - 1./4).*alpha + 5.*(a - 1).*((gamma + (3.*sigma)./5).*a.^2 + (gamma + (6.*sigma)./5).*a + sigma./5)./4).*(SG + IG).*beta.^2.*tau + beta.^3.*alpha.*a.^2.*(IP + IG).^3).*d.^3 + (a.*(SG + IG).^3.*(a - 1).*(alpha + gamma).*(alpha + gamma + 2.*sigma).*tau.^3 + 7.*(IP + IG).*(SG + IG).^2.*beta.*((a.^2 + 2./7.*a + 1./7).*alpha.^2 + (((12.*gamma)./7 + (10.*sigma)./7).*a.^2 + (-(3.*gamma)./7 + sigma./7).*a + gamma./7 - sigma./7).*alpha + 5.*((gamma.^2 + 2.*gamma.*sigma + 2./5.*sigma.^2).*a + 3.*(gamma + sigma./3).*sigma./5).*(a - 1)./7).*tau.^2 + 7.*(IP + IG).^2.*((a.^2 + 5./7.*a).*alpha.^2 + (((11.*gamma)./7 + (8.*sigma)./7).*a.^2 + (-gamma./7 + (5.*sigma)./7).*a - (2.*gamma)./7 - (3.*sigma)./7).*alpha + 5.*(a - 1).*((gamma.^2 + 2.*gamma.*sigma + 3./5.*sigma.^2).*a + sigma.*(gamma + (3.*sigma)./5))./7).*(SG + IG).*beta.^2.*tau + (IP + IG).^3.*alpha.*((a + 2).*alpha + gamma.*a + 2.*sigma).*a.*beta.^3).*d.^2 + (sigma.*(SG + IG).^3.*(alpha + gamma).^2.*(a - 1).*tau.^3 + 4.*(IP + IG).*((a + 1./2).*alpha.^3 + ((2.*gamma + (11.*sigma)./4).*a + gamma + sigma).*alpha.^2 + ((gamma.^2 + 4.*gamma.*sigma + sigma.^2).*a + gamma.^2./2 - gamma.*sigma./4 + sigma.^2./4).*alpha + 5.*(a - 1).*gamma.*(gamma + (3.*sigma)./5).*sigma./4).*(SG + IG).^2.*beta.*tau.^2 + 6.*(IP + IG).^2.*((a + 1./2).*alpha.^3 + (((5.*gamma)./3 + 2.*sigma).*a + gamma./3 + (5.*sigma)./6).*alpha.^2 + ((2./3.*gamma.^2 + 8./3.*gamma.*sigma + 2./3.*sigma.^2).*a - gamma.^2./6 - gamma.*sigma./2 + sigma.^2./6).*alpha + 5.*(a - 1).*sigma.*(gamma.^2 + gamma.*sigma + 1./5.*sigma.^2)./6).*(SG + IG).*beta.^2.*tau + 2.*(IP + IG).^3.*alpha.*(alpha + sigma).*beta.^3.*((a + 1./2).*alpha + gamma.*a + sigma./2)).*d + (IP + IG).*alpha.*(alpha + gamma).*((SG + IG).^2.*(alpha.^2 + (2.*gamma + 5.*sigma).*alpha + gamma.^2 + 5.*gamma.*sigma + 5.*sigma.^2).*tau.^2 + 2.*(IP + IG).*(SG + IG).*beta.*(alpha.^2 + (gamma + (7.*sigma)./2).*alpha + (3.*sigma.*(gamma + (5.*sigma)./3))./2).*tau + beta.^2.*(IP + IG).^2.*(alpha + sigma).^2).*beta).*(a.*d + alpha).*N.^2 + 3.*(IP + IG).^2.*(beta.*(IP + IG) + tau.*(SG + IG)).^2.*tau.^2.*E.^10.*(SG + IG).^2.*beta.^2.*((tau.*a.*(SG + IG) + (IP + IG).*(a + 1).*beta).*(a - 1).*a.*d.^3 + (2.*(a.*alpha + ((a - 1).*(gamma + sigma))./2).*a.*(SG + IG).*tau + 2.*(IP + IG).*beta.*((a.^2 - 1./2).*alpha + (a - 1).*((gamma + sigma).*a + sigma./2))).*d.^2 + (2.*((a + 1./2).*alpha.^2 + ((gamma + sigma).*a + gamma./2).*alpha + gamma.*sigma.*(a - 1)./2).*(SG + IG).*tau + 2.*(IP + IG).*beta.*(alpha.^2.*a + (a - 1./2).*(gamma + sigma).*alpha + (gamma + sigma./2).*(a - 1).*sigma)).*d + ((IP + IG).*(alpha + sigma).*beta + tau.*(SG + IG).*(alpha + gamma + 2.*sigma)).*alpha.*(alpha + gamma)).*(a.*d + alpha).*N + (IP + IG).^2.*(beta.*(IP + IG) + tau.*(SG + IG)).^3.*tau.^2.*(a.*beta.*(IP + IG).*(a - 1).*d.^2 + (alpha.*a.*(SG + IG).*tau + beta.*(IP + IG).*(a - 1).*(alpha + sigma)).*d + tau.*alpha.*(SG + IG).*(alpha + gamma)).*E.^12.*(SG + IG).^2.*beta.^2.*(a.*d + alpha)).*N.*E.^2.*(N.*q - b)./(N.^3.*d.*(a.*d + sigma).*(d + alpha + gamma).*(a.*d + alpha + gamma + sigma) + (2.*a.*(((IP + IG).*(a + 1).*beta)./2 + tau.*a.*(SG + IG)).*d.^3 + ((IP + IG).*((a + 1).^2.*alpha + (2.*gamma + 2.*sigma).*a + sigma).*beta + a.*(SG + IG).*tau.*((a + 2).*alpha + gamma.*a + 2.*gamma + 2.*sigma)).*d.^2 + ((IP + IG).*((a + 1).*alpha.^2 + (a + 1).*(gamma + 2.*sigma).*alpha + 2.*gamma.*sigma + sigma.^2).*beta + (a.*alpha + (gamma + sigma).*a + sigma).*(SG + IG).*tau.*(alpha + gamma)).*d + beta.*alpha.*sigma.*(IP + IG).*(alpha + gamma + sigma)).*E.^2.*N.^2 + (a.*((IP + IG).^2.*beta.^2 + tau.*(SG + IG).*(IP + IG).*(a + 2).*beta + tau.^2.*a.*(SG + IG).^2).*d.^2 + ((IP + IG).^2.*((a + 1).*alpha + sigma).*beta.^2 + 2.*(IP + IG).*((a + 1).*alpha + (gamma + sigma./2).*a + sigma./2).*(SG + IG).*tau.*beta + tau.^2.*a.*(SG + IG).^2.*(alpha + gamma)).*d + (IP + IG).*beta.*((IP + IG).*(alpha + sigma).*beta + tau.*(SG + IG).*(alpha + gamma + 2.*sigma)).*alpha).*E.^4.*N + (IP + IG).*beta.*E.^6.*(SG + IG).*tau.*(beta.*(IP + IG) + tau.*(SG + IG)).*(a.*d + alpha)).^3);

% Special cases
disonly = R0_I<1 & R0_D>1;
infoonly = R0_I>1 & R0_D<1;
neither = R0_I<=1 & R0_D<=1;

if(alpha>0)
    w_1(disonly) = -1;
    ES(disonly) = -1;
else
    w_1(disonly) = 0;
    ES(disonly) = 0;
end

if(a<1)
    w_1(infoonly) = 1;
    ES(infoonly) = 1;
else
    w_1(infoonly) = 0;
    ES(infoonly) = 0;
end 

w_1(neither) = 0;
ES(neither) = 0;
```
