## Supplementary material for "Social Information use Shapes the Coevolution of Sociality and Virulence": Source code: singstrat_fastinfo.m

```
function [singstrat,outcome] = singstrat_fastinfo(a,Emin,Emax,d,alpha,beta,gamma,sigma,tau,res1)

%Classification for interior singular strategies:
%	w_1(E<E*)=0, w_1(E>E*)=1: unstable (trait increases when info present): outcome 1
%	w_1(E<E*)=0, w_1(E>E*)=-1: ˜pseudo-stable™ (trait decreases when disease present): outcome 2
%	w_1(E<E*)=-1, w_1(E>E*)=1: convergence unstable, CS=1 (repeller): outcome 3
%	w_1(E<E*)=1, w_1(E>E*)=-1 & ES<0: convergence stable & evolutionarily stable: outcome 4
%	w_1(E<E*)=1, w_1(E>E*)=-1 & ES>0: convergence stable & evolutionarily unstable: outcome 5
%	w_1(E)=1: trait always increasing: outcome 6
%	w_1(E)=-1: trait always decreasing: outcome 7
%	uncertain, potential error: outcome 8
%	uncertain, potential error: outcome 9

%%% First pass
E = linspace(Emin,Emax,res1)';
E(E==0)=[];
[w_1,ES] = fitness_grad_sign_fastinfo(a,E,d,alpha,beta,gamma,sigma,tau);

if(all(isnan(w_1)) || all(w_1==0))
    singstrat = NaN;
    outcome = NaN;
elseif(all(w_1<0)) % All <0 implies E*<=Emin
    singstrat = Emin;
    outcome = 7;
elseif(all(w_1>0)) % All >0 implies E*>=Emax
    singstrat = Emax;
    outcome = 6;
else % Must have at least 1 intermediate SS - narrow down & find CS & ES
    
    % These are the values just before each sign change
    temp = find(diff(w_1));
    
    % Increase resolution to find singstrat again
    singstrat = E(temp);%NaN*zeros(length(temp),1);
    outcome = NaN*zeros(length(temp),1);
    
    for i=1:length(temp)
        
        % This is the value just before the sign change
        temp2 = temp(i);
        if(w_1(temp2-1)==0)
            if(w_1(temp2+1)>0) % unstable (trait increases when info present): outcome 1
                outcome(i) = 1;
            else % ˜pseudo-stable™ (trait decreases when disease present): outcome 2
                outcome(i) = 2;
            end
        elseif(w_1(temp2-1)<0) % convergence unstable, CS=1 (repeller): outcome 3
            outcome(i) = 3;
        else % w_1(E<E*)=1, w_1(E>E*)=-1
            if(ES(temp2)<0) % convergence stable & evolutionarily stable: outcome 4
                outcome(i) = 4;
            else % convergence stable & evolutionarily unstable: outcome 5
                outcome(i) = 5;
            end
        end
    end
end
```
