## Supplementary material for "Social Information use Shapes the Coevolution of Sociality and Virulence": Source code: sociality_fastinfo_mex.c

```
/*******************************************************************************************
 * [t,x,EQFLAG] = sociality_fastinfo_mex(t_max,a,b,E,d,q,alpha,beta,gamma,sigma,tau,eqtol,init_pop)
 ******************************************************************************************/

#include <mex.h>
#include <math.h>

/***********************************
 * Constant parameter values
 ***********************************/
#define MAXSTEPS 1e6 /* Maximum number of steps for ODE solver */
#define INTERVAL 1e2 /* Check if the system is close to equilibrium */
#define EPS 1e-6 /* ODE solver tolerance */
#define TINY 1e-30 /* Constant value for solver */
/* RK solver parameters */
#define b21 0.2
#define b31 3.0/40.0
#define b32 9.0/40.0
#define b41 0.3
#define b42 -0.9
#define b43 1.2
#define b51 -11.0/54.0
#define b52 2.5
#define b53 -70.0/27.0
#define b54 35.0/27.0
#define b61 1631.0/55296
#define b62 175.0/512.0
#define b63 575.0/13824.0
#define b64 44275.0/110592
#define b65 253.0/4096.0
#define c1 37.0/378.0
#define c3 250.0/621.0
#define c4 125.0/594.0
#define c6 512.0/1771.0
#define dc5 -277.00/14336

/*************************************
 * Define structure for model parameters
 *************************************/
struct PARAM{
    double t_max;
    double a;
    double b;
    double E;
    double EE;
    double d;
    double q; 
    double alpha;
    double beta;
    double gamma;
    double sigma;
    double tau;
    double eqtol;
};

/*************************************
 * Function prototypes
 *************************************/
int my_rungkut (double *T, double *x_out, double *EQFLAG, double *init_pop, struct PARAM *p);
void rkqs(double *x, double *dxdt, double *h, double *hnext, double *x_scale, struct PARAM *p);
void rkck(double *x, double *dxdt, double *xout, double *xerr, double h, struct PARAM *p);
void dynamic(double *x, double *dxdt, struct PARAM *p);
double FMAX(double, double);
double FMIN(double, double);

/*************************************
 * Main function
 *************************************/
void mexFunction(int nlhs, mxArray *plhs[], int nrhs, const mxArray *prhs[])
{
    double *T, *x, *t_temp, *x_temp, *EQFLAG, *init_pop, *parameter;
    int i, j, k, colLen, maxsteps;
    struct PARAM p;
    
    /* Allocate inputs */
    if(nrhs!=13){
        mexErrMsgTxt("Incorrect number of input arguments!\n");
    }
    else{                
        parameter= mxGetPr(prhs[0]);
        p.t_max= *parameter;
        parameter= mxGetPr(prhs[1]);
        p.a= *parameter;
        parameter= mxGetPr(prhs[2]);
        p.b= *parameter;
        parameter= mxGetPr(prhs[3]);
        p.E= *parameter;
        p.EE= p.E*p.E;
        parameter= mxGetPr(prhs[4]);
        p.d= *parameter;   
        parameter= mxGetPr(prhs[5]);        
        p.q= *parameter;
        parameter= mxGetPr(prhs[6]);
        p.alpha= *parameter;
        parameter= mxGetPr(prhs[7]);
        p.beta= *parameter;
        parameter= mxGetPr(prhs[8]);
        p.gamma= *parameter;
        parameter= mxGetPr(prhs[9]);
        p.sigma= *parameter;
        parameter= mxGetPr(prhs[10]);
        p.tau= *parameter;
        parameter= mxGetPr(prhs[11]);
        p.eqtol= *parameter;
        init_pop= mxGetPr(prhs[12]);        
    }
    maxsteps = (int)MAXSTEPS;
    
    /* Allocate memory */
    t_temp = malloc(maxsteps*sizeof(double));
    x_temp = malloc(maxsteps*4*sizeof(double));
    
    /* Initialise this output */           
    plhs[2] = mxCreateDoubleMatrix(1, 1, mxREAL);
    EQFLAG = mxGetPr(plhs[2]);
    EQFLAG[0] = 0;
    
    /* Call ODE solver */
    colLen = my_rungkut(t_temp, x_temp, EQFLAG, init_pop, &p);
    
    /* Create outputs */
    plhs[0] = mxCreateDoubleMatrix(colLen, 1, mxREAL);
    plhs[1] = mxCreateDoubleMatrix(colLen, 2, mxREAL);
    T = mxGetPr(plhs[0]);
    x = mxGetPr(plhs[1]);
    
    /* Copy data to outputs */
    for (i=0;i<colLen;i++){
        T[i] = t_temp[i];
        for (j=0;j<2;j++) {
            x[i + j*colLen] = x_temp[i + j*maxsteps];
        }
    }
    
    /* Free memory */
    free(t_temp);
    free(x_temp);
    
    return;
}

/*****************************************
 * ODE solver
 ****************************************/
int my_rungkut (double *T, double *x_out, double *EQFLAG, double *init_pop, struct PARAM *p){
    
    double x[2], dxdt[2], x_scale[2], xmin[2], xmax[2], hnext[1], h[1], t, nextcheck;
    int i, j, k, exitflag, count, maxsteps;
    
    /* Other parameters */
    exitflag = 1;
    count=0;
    k=1;
    h[0] = 1e-3;
    hnext[0] = 1e-3;
    t=0;
    nextcheck = INTERVAL;
    maxsteps = (int)MAXSTEPS;
    
    /* Initialise populations */
    for(i=0;i<2;i++){
        x[i] = init_pop[i];
        xmin[i] = x[i];
        xmax[i] = x[i];
    }
    
    /* Update output */
    T[0]=t;
    for (i=0; i<2; i++) {
        x_out[i*maxsteps] = x[i];
    }
    
    /* Main loop: */
    do{
        /* This ensures the final step lands us on the final time point */
        if(1.1*hnext[0]>(p->t_max-t)){
            hnext[0] = p->t_max-t;
            h[0] = p->t_max-t;
            t=p->t_max;
            exitflag=0;
        }
        else{
            h[0] = hnext[0];
            t+=h[0];
        }
        if(t>=p->t_max) {
            t=p->t_max;
            exitflag=0;
        }
        /* This is where the equations are first solved */
        dynamic(x, dxdt, p);

        /* Adjust the step size to maintain accuracy */
        for (i=0; i<2; i++){
            x[i] = FMAX(x[i],0);
            x_scale[i]=fabs(x[i])+fabs(dxdt[i]*(*h))+TINY;
        }
        
        /* RK solver & adaptive step-size */
        rkqs(x, dxdt, h, hnext, x_scale, p);
        
        /* Make sure nothin has gone negative */
        for (i=0; i<2; i++){
            x[i] = FMAX(x[i],0);
        }
        
        /* Update output */
        count++;
        T[count] = t;
        for (i=0; i<2; i++) {
            x_out[count + i*maxsteps] = x[i];
        }
    
        /* For equilibrium check */
        for (i=0; i<2; i++){
            xmin[i] = FMIN(xmin[i],x[i]);
            xmax[i] = FMAX(xmax[i],x[i]);
        }
        
        /* Check if we're close to equilibrium */
        if(t>nextcheck){
            exitflag = 0;
            for (i=0; i<2; i++){
                if(fabs(xmax[i]-xmin[i])>p->eqtol){
                    exitflag = 1;
                    break;
                }
            }
            /* If close to equilibrium, then break */
            if(exitflag==0){
                t=p->t_max;
                T[count] = t;
                EQFLAG[0] = 1;
                break;
            }
            
            /* If not, then reset min/max values for each class */
            nextcheck+=INTERVAL;
            for (i=0; i<2; i++){
                xmin[i] = x[i];
                xmax[i] = x[i];
            }
        }
    }while(count<(maxsteps-1) && t<=p->t_max && exitflag);
    count++;
    
    return count;
}

/***************************************
 * This generates the adaptive step-size
 **************************************/
void rkqs(double *x, double *dxdt, double *h, double *hnext, double *x_scale, struct PARAM *p)
{
    double x_temp[2], x_err[2], htemp, errmax;
    int i, j, count;
    
    count = 0;
    for(;;)
    {
        rkck(x, dxdt, x_temp, x_err, *h, p);
        
        errmax= 0.0;
        for(i=0;i<2;i++){
            errmax= FMAX(errmax, fabs(x_err[i]/(x_scale[i])));
        }
        errmax/= EPS;
        if(errmax<=1.0) break;
        htemp= 0.9*(*h)*pow(errmax, -0.25);
        *h= (*h>=0.0 ? FMAX(htemp, 0.1*(*h)) : FMIN(htemp, 0.1*(*h)));
        count++;
            
        if(count>1e4){
            printf("%f\n",errmax);
            mexErrMsgTxt("stuck in loop!\n");
            break;
        }
    }    
    if(errmax > 1.89E-4) {
        *hnext= 0.9*(*h)*pow(errmax, -0.2);
    }
    else {
        *hnext= 5.0*(*h);
    }    
    *hnext = FMAX(*hnext, p->t_max/MAXSTEPS);
    
    for(i=0;i<2;i++){
        x[i] = x_temp[i];
    }
}

/**************************************
 * Standard RK solver
 **************************************/
void rkck(double *x, double *dxdt, double *xout, double *xerr, double h, struct PARAM *p){
    int i, j;
    double xk1[2], xk2[2], xk3[2], xk4[2], xk5[2], xk6[2], xtemp[2];
    double dc1=c1-2825.0/27648.0, dc3=c3-18575.0/48384.0, dc4=c4-13525.0/55296.0, 
            dc6=c6-0.25;
    
    for(i=0;i<2;i++){
        xtemp[i] = x[i] + b21*h*dxdt[i];
    }
    dynamic(xtemp, xk2, p);
    
    for(i=0;i<2;i++){
        xtemp[i] = x[i]+h*(b31*dxdt[i]+b32*xk2[i]);
    }    
    dynamic(xtemp, xk3, p);
    
    for(i=0;i<2;i++){
        xtemp[i] = x[i]+h*(b41*dxdt[i]+b42*xk2[i]+b43*xk3[i]);
    }
    dynamic(xtemp, xk4, p);
    
    for(i=0;i<2;i++){
        xtemp[i] = x[i]+h*(b51*dxdt[i]+b52*xk2[i]+b53*xk3[i]+b54*xk4[i]);
    }
    dynamic(xtemp, xk5, p);
    
    for(i=0;i<2;i++){
        xtemp[i] = x[i]+h*(b61*dxdt[i]+b62*xk2[i]+b63*xk3[i]+b64*xk4[i]+b65*xk5[i]);
    }
    dynamic(xtemp, xk6, p);
    
    for(i=0;i<2;i++){
        xout[i]= x[i]+h*(c1*dxdt[i]+c3*xk3[i]+c4*xk4[i]+c6*xk6[i]);
        xerr[i]= h*(dc1*dxdt[i]+dc3*xk3[i]+dc4*xk4[i]+dc5*xk5[i]+dc6*xk6[i]);
    }
}

/**************************************
 * Population and evolutionary dynamics
 **************************************/
void dynamic(double *x, double *dxdt, struct PARAM *p){
    
    double S, I, N, GEE;
    int i, j;
    
    /* Define classes */
    S = x[0];
    I = x[1];
    N = S+I;
    
    GEE = FMAX(0, 1 - (p->sigma/FMAX(TINY,p->tau*p->EE)));
    
    dxdt[0] = (p->b - p->q*N)*N - p->beta*p->EE*S*I/FMAX(TINY,N) - p->d*(1-(1-p->a)*GEE)*S + p->gamma*I;
    dxdt[1] = p->beta*p->EE*S*I/FMAX(TINY,N) - (p->d*(1-(1-p->a)*GEE) + p->alpha + p->gamma)*I;
}

/***************************************
 * Return maximum of two inputs
 ***************************************/
double FMAX(double l, double r)
{
    if(l>r)return l;
    else   return r;
}

/***************************************
 * Return minimum of two inputs
 ***************************************/
double FMIN(double l, double r)
{
    if(l<r)return l;
    else   return r;
}
```
