## Supplementary material for "Social Information use Shapes the Coevolution of Sociality and Virulence": Source code: sociality_slowinfo_competition_mex.c

```
/***********************************************************************************************************
 * [t,SP,SG,IP,IG,EQFLAG] = sociality_slowinfo_competition_mex(t_max,a,b,E,d,q,alpha,beta,gamma,sigma,tau,eqtol,init_pop,strain_total)
 ***********************************************************************************************************/

#include <mex.h>
#include <math.h>

/***********************************
 * Constant parameter values
 ***********************************/
#define MAXSTEPS 1e6 /* Maximum number of steps for ODE solver */
#define INTERVAL 1e2 /* Check if the system is close to equilibrium */
#define EPS 1e-6 /* ODE solver tolerance */
#define TINY 1e-30 /* Constant value for solver */
#define TINY2 1e-30 /* Constant value for solver to avoid tolerance issues */
/* RK solver parameters */
#define b21 0.2
#define b31 3.0/40.0
#define b32 9.0/40.0
#define b41 0.3
#define b42 -0.9
#define b43 1.2
#define b51 -11.0/54.0
#define b52 2.5
#define b53 -70.0/27.0
#define b54 35.0/27.0
#define b61 1631.0/55296
#define b62 175.0/512.0
#define b63 575.0/13824.0
#define b64 44275.0/110592
#define b65 253.0/4096.0
#define c1 37.0/378.0
#define c3 250.0/621.0
#define c4 125.0/594.0
#define c6 512.0/1771.0
#define dc5 -277.00/14336

/*************************************
 * Define structure for model parameters
 *************************************/
struct PARAM{
    double t_max;
    double a;
    double b;
    double d;
    double q; 
    double alpha;
    double beta;
    double gamma;
    double sigma;
    double tau;
    double eqtol;
    int strain_total;
};

/*************************************
 * Function prototypes
 *************************************/
int my_rungkut (double *T, double *SP_out, double *SG_out, double *IP_out, double *IG_out, double *EQFLAG, double *init_pop, double *E, struct PARAM *p);
void rkqs(double *SP, double *SG, double *IP, double *IG,  double *DSPDT, double *DSGDT, double *DIPDT, double *DIGDT, double *h, double *hnext, double *SP_SCALE, double *SG_SCALE, double *IP_SCALE, double *IG_SCALE, double *E, struct PARAM *p);
void rkck(double *SP, double *SG, double *IP, double *IG,  double *DSPDT, double *DSGDT, double *DIPDT, double *DIGDT, double *SPout, double *SGout, double *IPout, double *IGout, double *SPerr, double *SGerr, double *IPerr, double *IGerr, double h, double *E, struct PARAM *p);
void dynamic(double *SP, double *SG, double *IP, double *IG,  double *DSPDT, double *DSGDT, double *DIPDT, double *DIGDT, double *E, struct PARAM *p);
double FMAX(double, double);
double FMIN(double, double);

/*************************************
 * Main function
 *************************************/
void mexFunction(int nlhs, mxArray *plhs[], int nrhs, const mxArray *prhs[])
{
    double *T, *SP, *SG, *IP, *IG, *EQFLAG, *init_pop, *E, *parameter;
    double *t_temp, *SP_temp, *SG_temp, *IP_temp, *IG_temp;
    int i, j, k, colLen, maxsteps;
    struct PARAM p;
    
    /* Allocate inputs */
    if(nrhs!=14){
        mexErrMsgTxt("Incorrect number of input arguments!\n");
    }
    else{
        parameter= mxGetPr(prhs[0]);
        p.t_max= *parameter;
        parameter= mxGetPr(prhs[1]);
        p.a= *parameter;
        parameter= mxGetPr(prhs[2]);
        p.b= *parameter;
        E= mxGetPr(prhs[3]);
        parameter= mxGetPr(prhs[4]);
        p.d= *parameter;   
        parameter= mxGetPr(prhs[5]);        
        p.q= *parameter;
        parameter= mxGetPr(prhs[6]);
        p.alpha= *parameter;
        parameter= mxGetPr(prhs[7]);
        p.beta= *parameter;
        parameter= mxGetPr(prhs[8]);
        p.gamma= *parameter;
        parameter= mxGetPr(prhs[9]);
        p.sigma= *parameter;
        parameter= mxGetPr(prhs[10]);
        p.tau= *parameter;
        parameter= mxGetPr(prhs[11]);
        p.eqtol= *parameter;
        init_pop= mxGetPr(prhs[12]);    
        parameter= mxGetPr(prhs[13]);
        p.strain_total= (int)*parameter;
    }
    maxsteps = (int)MAXSTEPS;
    
    /* Allocate memory */
    t_temp = malloc(maxsteps*sizeof(double));
    SP_temp = malloc(maxsteps*(p.strain_total)*sizeof(double));
    SG_temp = malloc(maxsteps*(p.strain_total)*sizeof(double));
    IP_temp = malloc(maxsteps*(p.strain_total)*sizeof(double));
    IG_temp = malloc(maxsteps*(p.strain_total)*sizeof(double));
    
    /* Initialise this output */           
    plhs[5] = mxCreateDoubleMatrix(1, 1, mxREAL);
    EQFLAG = mxGetPr(plhs[5]);
    EQFLAG[0] = 0;
    
    /* Call ODE solver */
    colLen = my_rungkut(t_temp, SP_temp, SG_temp, IP_temp, IG_temp, EQFLAG, init_pop, E, &p);
    
    /* Create outputs */
    plhs[0] = mxCreateDoubleMatrix(colLen, 1, mxREAL);
    plhs[1] = mxCreateDoubleMatrix(colLen, p.strain_total, mxREAL);
    plhs[2] = mxCreateDoubleMatrix(colLen, p.strain_total, mxREAL);
    plhs[3] = mxCreateDoubleMatrix(colLen, p.strain_total, mxREAL);
    plhs[4] = mxCreateDoubleMatrix(colLen, p.strain_total, mxREAL);
    T = mxGetPr(plhs[0]);
    SP = mxGetPr(plhs[1]);
    SG = mxGetPr(plhs[2]);
    IP = mxGetPr(plhs[3]);
    IG = mxGetPr(plhs[4]);
    
    /* Copy data to outputs */
    for (i=0;i<colLen;i++){
        T[i] = t_temp[i];
        for (j=0;j<p.strain_total;j++) {
            SP[i + j*colLen] = SP_temp[i + j*maxsteps];
            SG[i + j*colLen] = SG_temp[i + j*maxsteps];
            IP[i + j*colLen] = IP_temp[i + j*maxsteps];
            IG[i + j*colLen] = IG_temp[i + j*maxsteps];
        }
    }
    
    /* Free memory */
    free(t_temp);
    free(SP_temp);
    free(SG_temp);
    free(IP_temp);
    free(IG_temp);
    
    return;
}

/*****************************************
 * ODE solver
 ****************************************/
int my_rungkut (double *T, double *SP_out, double *SG_out, double *IP_out, double *IG_out, double *EQFLAG, double *init_pop, double *E, struct PARAM *p){
    
    double *SP, *SG, *IP, *IG, *DSPDT, *DSGDT, *DIPDT, *DIGDT, *SP_SCALE, *SG_SCALE, *IP_SCALE, *IG_SCALE;
    double *SPMIN, *SPMAX, *SGMIN, *SGMAX, *IPMIN, *IPMAX, *IGMIN, *IGMAX, hnext[1], h[1];
    double t, nextcheck;
    int i, j, k, exitflag, count, maxsteps;
    
    /* Allocate memory */
    SP = malloc(p->strain_total*sizeof(double));
    SG = malloc(p->strain_total*sizeof(double));
    IP = malloc(p->strain_total*sizeof(double));
    IG = malloc(p->strain_total*sizeof(double));
    DSPDT = malloc(p->strain_total*sizeof(double));
    DSGDT = malloc(p->strain_total*sizeof(double));
    DIPDT = malloc(p->strain_total*sizeof(double));
    DIGDT = malloc(p->strain_total*sizeof(double));
    SP_SCALE = malloc(p->strain_total*sizeof(double));
    SG_SCALE = malloc(p->strain_total*sizeof(double));
    IP_SCALE = malloc(p->strain_total*sizeof(double));
    IG_SCALE = malloc(p->strain_total*sizeof(double));
    SPMIN = malloc(p->strain_total*sizeof(double));
    SGMIN = malloc(p->strain_total*sizeof(double));
    IPMIN = malloc(p->strain_total*sizeof(double));
    IGMIN = malloc(p->strain_total*sizeof(double));    
    SPMAX = malloc(p->strain_total*sizeof(double));
    SGMAX = malloc(p->strain_total*sizeof(double));
    IPMAX = malloc(p->strain_total*sizeof(double));
    IGMAX = malloc(p->strain_total*sizeof(double));
    
    /* Other parameters */
    exitflag = 1;
    count=0;
    k=1;
    h[0] = 1e-3;
    hnext[0] = 1e-3;
    t=0;
    nextcheck = INTERVAL;
    maxsteps = (int)MAXSTEPS;
    
    /* Initialise populations */
    for(i=0;i<p->strain_total;i++){
        SP[i] = init_pop[i*4];
        SG[i] = init_pop[i*4+1];
        IP[i] = init_pop[i*4+2];
        IG[i] = init_pop[i*4+3];
    }
    
    /* Initialise equilibrium arrays */
    for(i=0;i<p->strain_total;i++){
        SPMIN[i] = SP[i];
        SPMAX[i] = SP[i];
        SGMIN[i] = SG[i];
        SGMAX[i] = SG[i];
        IPMIN[i] = IP[i];
        IPMAX[i] = IP[i];
        IGMIN[i] = IG[i];
        IGMAX[i] = IG[i];
    }
    
    /* Update output */
    T[0]=t;
    for (i=0; i<p->strain_total; i++) {
        SP_out[i*maxsteps] = SP[i];
        SG_out[i*maxsteps] = SG[i];
        IP_out[i*maxsteps] = IP[i];
        IG_out[i*maxsteps] = IG[i];
    }
    
    /* Main loop: */
    do{
        /* This ensures the final step lands us on the final time point */
        if(1.1*hnext[0]>(p->t_max-t)){
            hnext[0] = p->t_max-t;
            h[0] = p->t_max-t;
            t=p->t_max;
            exitflag=0;
        }
        else{
            h[0] = hnext[0];
            t+=h[0];
        }
        if(t>=p->t_max) {
            t=p->t_max;
            exitflag=0;
        }
        /* This is where the equations are first solved */
        dynamic(SP, SG, IP, IG, DSPDT, DSGDT, DIPDT, DIGDT, E, p);

        /* Adjust the step size to maintain accuracy */
        for (i=0; i<p->strain_total; i++){
            SP[i] = FMAX(SP[i],0);
            SG[i] = FMAX(SG[i],0);
            IP[i] = FMAX(IP[i],0);
            IG[i] = FMAX(IG[i],0);
            SP_SCALE[i]=fabs(SP[i])+fabs(DSPDT[i]*(*h))+TINY;
            SG_SCALE[i]=fabs(SG[i])+fabs(DSGDT[i]*(*h))+TINY;
            IP_SCALE[i]=fabs(IP[i])+fabs(DIPDT[i]*(*h))+TINY;
            IG_SCALE[i]=fabs(IG[i])+fabs(DIGDT[i]*(*h))+TINY;
        }
        
        /* RK solver & adaptive step-size */
        rkqs(SP, SG, IP, IG, DSPDT, DSGDT, DIPDT, DIGDT, h, hnext, SP_SCALE, SG_SCALE, IP_SCALE, IG_SCALE, E, p);
        
        /* Make sure nothin has gone negative */
        for (i=0; i<p->strain_total; i++){
            SP[i] = FMAX(SP[i],0);
            SG[i] = FMAX(SG[i],0);
            IP[i] = FMAX(IP[i],0);
            IG[i] = FMAX(IG[i],0);
        }
        
        /* Update output */
        count++;
        T[count] = t;
        for (i=0; i<p->strain_total; i++) {
            SP_out[count + i*maxsteps] = SP[i];
            SG_out[count + i*maxsteps] = SG[i];
            IP_out[count + i*maxsteps] = IP[i];
            IG_out[count + i*maxsteps] = IG[i];
        }
    
        /* For equilibrium check */
        for (i=0; i<p->strain_total; i++){
            SPMIN[i] = FMIN(SPMIN[i],SP[i]);
            SGMIN[i] = FMIN(SGMIN[i],SG[i]);
            SPMAX[i] = FMAX(SPMAX[i],SP[i]);
            SGMAX[i] = FMAX(SGMAX[i],SG[i]);
            IPMIN[i] = FMIN(IPMIN[i],IP[i]);
            IGMIN[i] = FMIN(IGMIN[i],IG[i]);
            IPMAX[i] = FMAX(IPMAX[i],IP[i]);
            IGMAX[i] = FMAX(IGMAX[i],IG[i]);
        }
        
        /* Check if we're close to equilibrium */
        if(t>nextcheck){
            exitflag = 0;
            for (i=0; i<p->strain_total; i++){
                if(fabs(SPMAX[i]-SPMIN[i])>p->eqtol || fabs(SGMAX[i]-SGMIN[i])>p->eqtol || fabs(IPMAX[i]-IPMIN[i])>p->eqtol || fabs(IGMAX[i]-IGMIN[i])>p->eqtol){
                    exitflag = 1;
                    break;
                }
            }
            /* If close to equilibrium, then break */
            if(exitflag==0){
                t=p->t_max;
                T[count] = t;
                EQFLAG[0] = 1;
                break;
            }
            
            /* If not, then reset min/max values for each class */
            nextcheck+=INTERVAL;
            for (i=0; i<p->strain_total; i++){
                SPMIN[i] = SP[i];
                SGMIN[i] = SG[i];
                SPMAX[i] = SP[i];
                SGMAX[i] = SG[i];
                IPMIN[i] = IP[i];
                IGMIN[i] = IG[i];
                IPMAX[i] = IP[i];
                IGMAX[i] = IG[i];
            }
        }
    }while(count<(maxsteps-1) && t<=p->t_max && exitflag);
    count++;
    
    /* Free memory */
    free(SP);
    free(SG);
    free(IP);
    free(IG);
    free(DSPDT);
    free(DSGDT);
    free(DIPDT);
    free(DIGDT);
    free(SP_SCALE);
    free(SG_SCALE);
    free(IP_SCALE);
    free(IG_SCALE);
    free(SPMIN);
    free(SGMIN);
    free(IPMIN);
    free(IGMIN);
    free(SPMAX);
    free(SGMAX);
    free(IPMAX);
    free(IGMAX);
    
    return count;
}

/***************************************
 * This generates the adaptive step-size
 **************************************/
void rkqs(double *SP, double *SG, double *IP, double *IG,  double *DSPDT, double *DSGDT, double *DIPDT, double *DIGDT, double *h, double *hnext, double *SP_SCALE, double *SG_SCALE, double *IP_SCALE, double *IG_SCALE, double *E, struct PARAM *p)
{
    double *SP_temp, *SG_temp, *IP_temp, *IG_temp, *SP_err, *SG_err, *IP_err, *IG_err;
    double htemp, errmax;
    int i, j, count;
    
    /* Allocate memory */
    SP_temp = malloc(p->strain_total*sizeof(double));
    SG_temp = malloc(p->strain_total*sizeof(double));
    IP_temp = malloc(p->strain_total*sizeof(double));
    IG_temp = malloc(p->strain_total*sizeof(double));
    SP_err = malloc(p->strain_total*sizeof(double));
    SG_err = malloc(p->strain_total*sizeof(double));
    IP_err = malloc(p->strain_total*sizeof(double));
    IG_err = malloc(p->strain_total*sizeof(double));
    
    count = 0;
    for(;;)
    {
        rkck(SP, SG, IP, IG, DSPDT, DSGDT, DIPDT, DIGDT, SP_temp, SG_temp, IP_temp, IG_temp, SP_err, SG_err, IP_err, IG_err, *h, E, p);
        
        errmax= 0.0;
        for(i=0;i<p->strain_total;i++){
            errmax= FMAX(errmax, fabs(SP_err[i]/(SP_SCALE[i])));
            errmax= FMAX(errmax, fabs(SG_err[i]/(SG_SCALE[i]))); 
            errmax= FMAX(errmax, fabs(IP_err[i]/(IP_SCALE[i])));
            errmax= FMAX(errmax, fabs(IG_err[i]/(IG_SCALE[i])));   
        }
        errmax/= EPS;
        if(errmax<=1.0) break;
        htemp= 0.9*(*h)*pow(errmax, -0.25);
        *h= (*h>=0.0 ? FMAX(htemp, 0.1*(*h)) : FMIN(htemp, 0.1*(*h)));
        count++;
            
        if(count>1e4){
            printf("%f\n",errmax);
            mexErrMsgTxt("stuck in loop!\n");
            break;
        }
    }    
    if(errmax > 1.89E-4) {
        *hnext= 0.9*(*h)*pow(errmax, -0.2);
    }
    else {
        *hnext= 5.0*(*h);
    }    
    *hnext = FMAX(*hnext, p->t_max/MAXSTEPS);
    
    for(i=0;i<p->strain_total;i++){
        SP[i] = SP_temp[i];
        SG[i] = SG_temp[i];
        IP[i] = IP_temp[i];
        IG[i] = IG_temp[i];
    }
    
    /* Free memory */
    free(SP_temp);
    free(SG_temp);
    free(IP_temp);
    free(IG_temp);
    free(SP_err);
    free(SG_err);
    free(IP_err);
    free(IG_err);
}

/**************************************
 * Standard RK solver
 **************************************/
void rkck(double *SP, double *SG, double *IP, double *IG,  double *DSPDT, double *DSGDT, double *DIPDT, double *DIGDT, double *SPout, double *SGout, double *IPout, double *IGout, double *SPerr, double *SGerr, double *IPerr, double *IGerr, double h, double *E, struct PARAM *p){
    int i, j;
    double *SPk1, *SPk2, *SPk3, *SPk4, *SPk5, *SPk6, *SPtemp;
    double *SGk1, *SGk2, *SGk3, *SGk4, *SGk5, *SGk6, *SGtemp;
    double *IPk1, *IPk2, *IPk3, *IPk4, *IPk5, *IPk6, *IPtemp;
    double *IGk1, *IGk2, *IGk3, *IGk4, *IGk5, *IGk6, *IGtemp;
    double dc1=c1-2825.0/27648.0, dc3=c3-18575.0/48384.0, dc4=c4-13525.0/55296.0, 
            dc6=c6-0.25;
    
    /* Allocate memory */    
    SPk1 = malloc(p->strain_total*sizeof(double));
    SPk2 = malloc(p->strain_total*sizeof(double));
    SPk3 = malloc(p->strain_total*sizeof(double));
    SPk4 = malloc(p->strain_total*sizeof(double));
    SPk5 = malloc(p->strain_total*sizeof(double));
    SPk6 = malloc(p->strain_total*sizeof(double));
    SPtemp = malloc(p->strain_total*sizeof(double));
    
    SGk1 = malloc(p->strain_total*sizeof(double));
    SGk2 = malloc(p->strain_total*sizeof(double));
    SGk3 = malloc(p->strain_total*sizeof(double));
    SGk4 = malloc(p->strain_total*sizeof(double));
    SGk5 = malloc(p->strain_total*sizeof(double));
    SGk6 = malloc(p->strain_total*sizeof(double));
    SGtemp = malloc(p->strain_total*sizeof(double));
    
    IPk1 = malloc(p->strain_total*sizeof(double));
    IPk2 = malloc(p->strain_total*sizeof(double));
    IPk3 = malloc(p->strain_total*sizeof(double));
    IPk4 = malloc(p->strain_total*sizeof(double));
    IPk5 = malloc(p->strain_total*sizeof(double));
    IPk6 = malloc(p->strain_total*sizeof(double));
    IPtemp = malloc(p->strain_total*sizeof(double));
    
    IGk1 = malloc(p->strain_total*sizeof(double));
    IGk2 = malloc(p->strain_total*sizeof(double));
    IGk3 = malloc(p->strain_total*sizeof(double));
    IGk4 = malloc(p->strain_total*sizeof(double));
    IGk5 = malloc(p->strain_total*sizeof(double));
    IGk6 = malloc(p->strain_total*sizeof(double));  
    IGtemp = malloc(p->strain_total*sizeof(double));  
    
    for(i=0;i<p->strain_total;i++){
        SPtemp[i] = SP[i] + b21*h*DSPDT[i];
        SGtemp[i] = SG[i] + b21*h*DSGDT[i];
        IPtemp[i] = IP[i] + b21*h*DIPDT[i];
        IGtemp[i] = IG[i] + b21*h*DIGDT[i];
    }
    dynamic(SPtemp, SGtemp, IPtemp, IGtemp, SPk2, SGk2, IPk2, IGk2, E, p);
    
    for(i=0;i<p->strain_total;i++){
        SPtemp[i] = SP[i]+h*(b31*DSPDT[i]+b32*SPk2[i]);
        SGtemp[i] = SG[i]+h*(b31*DSGDT[i]+b32*SGk2[i]);
        IPtemp[i] = IP[i]+h*(b31*DIPDT[i]+b32*IPk2[i]);
        IGtemp[i] = IG[i]+h*(b31*DIGDT[i]+b32*IGk2[i]);
    }    
    dynamic(SPtemp, SGtemp, IPtemp, IGtemp, SPk3, SGk3, IPk3, IGk3, E, p);
    
    for(i=0;i<p->strain_total;i++){
        SPtemp[i] = SP[i]+h*(b41*DSPDT[i]+b42*SPk2[i]+b43*SPk3[i]);
        SGtemp[i] = SG[i]+h*(b41*DSGDT[i]+b42*SGk2[i]+b43*SGk3[i]);
        IPtemp[i] = IP[i]+h*(b41*DIPDT[i]+b42*IPk2[i]+b43*IPk3[i]);
        IGtemp[i] = IG[i]+h*(b41*DIGDT[i]+b42*IGk2[i]+b43*IGk3[i]);
    }
    dynamic(SPtemp, SGtemp, IPtemp, IGtemp, SPk4, SGk4, IPk4, IGk4, E, p);
    
    for(i=0;i<p->strain_total;i++){
        SPtemp[i] = SP[i]+h*(b51*DSPDT[i]+b52*SPk2[i]+b53*SPk3[i]+b54*SPk4[i]);
        SGtemp[i] = SG[i]+h*(b51*DSGDT[i]+b52*SGk2[i]+b53*SGk3[i]+b54*SGk4[i]);
        IPtemp[i] = IP[i]+h*(b51*DIPDT[i]+b52*IPk2[i]+b53*IPk3[i]+b54*IPk4[i]);
        IGtemp[i] = IG[i]+h*(b51*DIGDT[i]+b52*IGk2[i]+b53*IGk3[i]+b54*IGk4[i]);
    }
    dynamic(SPtemp, SGtemp, IPtemp, IGtemp, SPk5, SGk5, IPk5, IGk5, E, p);
    
    for(i=0;i<p->strain_total;i++){
        SPtemp[i] = SP[i]+h*(b61*DSPDT[i]+b62*SPk2[i]+b63*SPk3[i]+b64*SPk4[i]+b65*SPk5[i]);
        SGtemp[i] = SG[i]+h*(b61*DSGDT[i]+b62*SGk2[i]+b63*SGk3[i]+b64*SGk4[i]+b65*SGk5[i]);        
        IPtemp[i] = IP[i]+h*(b61*DIPDT[i]+b62*IPk2[i]+b63*IPk3[i]+b64*IPk4[i]+b65*IPk5[i]);
        IGtemp[i] = IG[i]+h*(b61*DIGDT[i]+b62*IGk2[i]+b63*IGk3[i]+b64*IGk4[i]+b65*IGk5[i]);
    }
    dynamic(SPtemp, SGtemp, IPtemp, IGtemp, SPk6, SGk6, IPk6, IGk6, E, p);
    
    for(i=0;i<p->strain_total;i++){
        SPout[i]= SP[i]+h*(c1*DSPDT[i]+c3*SPk3[i]+c4*SPk4[i]+c6*SPk6[i]);
        SPerr[i]= h*(dc1*DSPDT[i]+dc3*SPk3[i]+dc4*SPk4[i]+dc5*SPk5[i]+dc6*SPk6[i]);
        SGout[i]= SG[i]+h*(c1*DSGDT[i]+c3*SGk3[i]+c4*SGk4[i]+c6*SGk6[i]);
        SGerr[i]= h*(dc1*DSGDT[i]+dc3*SGk3[i]+dc4*SGk4[i]+dc5*SGk5[i]+dc6*SGk6[i]);
        IPout[i]= IP[i]+h*(c1*DIPDT[i]+c3*IPk3[i]+c4*IPk4[i]+c6*IPk6[i]);
        IPerr[i]= h*(dc1*DIPDT[i]+dc3*IPk3[i]+dc4*IPk4[i]+dc5*IPk5[i]+dc6*IPk6[i]);
        IGout[i]= IG[i]+h*(c1*DIGDT[i]+c3*IGk3[i]+c4*IGk4[i]+c6*IGk6[i]);
        IGerr[i]= h*(dc1*DIGDT[i]+dc3*IGk3[i]+dc4*IGk4[i]+dc5*IGk5[i]+dc6*IGk6[i]);
    }

    /* Free memory */
    free(SPk1);
    free(SPk2);
    free(SPk3);
    free(SPk4);
    free(SPk5);
    free(SPk6);
    free(SGk1);
    free(SGk2);
    free(SGk3);
    free(SGk4);
    free(SGk5);
    free(SGk6);
    free(SPtemp);
    free(SGtemp);
    free(IPk1);
    free(IPk2);
    free(IPk3);
    free(IPk4);
    free(IPk5);
    free(IPk6);
    free(IGk1);
    free(IGk2);
    free(IGk3);
    free(IGk4);
    free(IGk5);
    free(IGk6);
    free(IPtemp);
    free(IGtemp);
}

/**************************************
 * Population and evolutionary dynamics
 **************************************/
void dynamic(double *SP, double *SG, double *IP, double *IG,  double *DSPDT, double *DSGDT, double *DIPDT, double *DIGDT, double *E, struct PARAM *p){
    
    int i, j;
    double *Ni, N, infection_mult, information_mult;
    
    /* Allocate memory */
    Ni = malloc(p->strain_total*sizeof(double));
    
    /* Population sums */
    N = 0;
    for(i=0;i<p->strain_total;i++){
        Ni[i] = SP[i] + SG[i] + IP[i] + IG[i];
        N += Ni[i];
    }
    
    /* Infections and information */
    infection_mult = 0;
    information_mult = 0;
    for(i=0;i<p->strain_total;i++){
        infection_mult += E[i]*(IP[i]+IG[i]);
        information_mult += E[i]*(SG[i]+IG[i]);
    }
    infection_mult = p->beta*FMAX(infection_mult,TINY2)/FMAX(N,TINY2);
    information_mult = p->tau*FMAX(information_mult,TINY2)/FMAX(N,TINY2);
    
    /* ODEs */
    for(i=0;i<p->strain_total;i++){
        DSPDT[i] = (p->b - p->q*N)*Ni[i] - E[i]*SP[i]*(infection_mult + information_mult) - p->d*SP[i] + p->gamma*IP[i] + p->sigma*SG[i];
        DSGDT[i] = E[i]*SP[i]*information_mult - E[i]*SG[i]*infection_mult - p->a*p->d*SG[i] + p->gamma*IG[i] - p->sigma*SG[i];
        DIPDT[i] = E[i]*SP[i]*infection_mult - E[i]*IP[i]*information_mult - (p->d + p->alpha + p->gamma)*IP[i] + p->sigma*IG[i];
        DIGDT[i] = E[i]*SG[i]*infection_mult + E[i]*IP[i]*information_mult - (p->a*p->d + p->alpha + p->gamma + p->sigma)*IG[i];
    }
    
    /* Free memory */
    free(Ni);
}

/***************************************
 * Return maximum of two inputs
 ***************************************/
double FMAX(double l, double r)
{
    if(l>r)return l;
    else   return r;
}

/***************************************
 * Return minimum of two inputs
 ***************************************/
double FMIN(double l, double r)
{
    if(l<r)return l;
    else   return r;
}
```
