## Supplementary material for "Social Information use Shapes the Coevolution of Sociality and Virulence": Source code: sociality_slowinfo_simulation.m

```
function [SOCIPLITY,DISPREV,INFOPREV] = sociality_slowinfo_simulation(t_max,a,b,Emin,Emax,Estart,d,q,alpha,beta,gamma,sigma,tau,res0,nevol,plotflag)

% Note: ensure sociality_slowinfo_competition_mex.c is compiled first

eqtol = 1e-3;
exttol = 1e-5;

E = linspace(Emin,Emax,res0);
SOCIPLITY = zeros(nevol,res0);
DISPREV = zeros(nevol,1);
INFOPREV = zeros(nevol,1);

% initial conditions
initial = find(E>=Estart,1);
E_current = E(initial);
index_current = initial;
strain_total = 1;
init_pop = ((b-d)/(4*q)*ones(1,4));
for ievol=1:nevol
    
    % Run solver
    [~,SP,SG,IP,IG,~] = sociality_slowinfo_competition_mex(t_max,a,b,E_current,d,q,alpha,beta,gamma,sigma,tau,eqtol,init_pop,strain_total);
    SP = SP(end,:);
    SG = SG(end,:);
    IP = IP(end,:);
    IG = IG(end,:);
    N = SP+SG+IP+IG;
    
    % Remove extinct classes
    extinct = (N/sum(N))<exttol;
    strain_total = strain_total-sum(extinct);
    SP(extinct) = [];
    SG(extinct) = [];
    IP(extinct) = [];
    IG(extinct) = [];
    N(extinct) = [];
    index_current(extinct) = [];
    E_current(extinct) = [];
    
    % Update tracker
    SOCIPLITY(ievol,index_current) = N/sum(N);
    DISPREV(ievol) = (sum(IP)+sum(IG))/sum(N);
    INFOPREV(ievol) = (sum(SG)+sum(IG))/sum(N);
    
    % Introduce rare mutant & update init_pop
    weightedprob = N/sum(N);
    cumsum1 = cumsum(weightedprob);
    r1 = rand*cumsum1(end);
    mutator_loc = (find(r1<cumsum1,1));
    mutator = index_current(mutator_loc);
    
    if(mutator==1) % Mutate up
        mutant = mutator+1;
    elseif(mutator==res0) % Mutate down
        mutant = mutator-1;
    else
        if(rand>0.5) % Mutate up
            mutant = mutator+1;
        else % Mutate down
            mutant = mutator-1;
        end
    end
    if(~ismember(mutant,index_current)) % New strain
        strain_total = strain_total+1;
        E_current(end+1) = E(mutant);
        index_current(end+1) = mutant;
                
        SP(end+1) = SP(mutator_loc)/10;
        SG(end+1) = SG(mutator_loc)/10;
        IP(end+1) = IP(mutator_loc)/10;
        IG(end+1) = IG(mutator_loc)/10;
    end
    
    % Update initial population
    init_pop = [];
    for i=1:strain_total
        init_pop = [init_pop,SP(i),SG(i),IP(i),IG(i)];
    end
end
N = SP+SG+IP+IG;

if(plotflag>0)
    figure(plotflag)
    clf
    a0=3;
    SOCIPLITY0=log10(SOCIPLITY);
    SOCIPLITY0(SOCIPLITY0<-a0)=-a0;
    SOCIPLITY0=(SOCIPLITY0+a0)/a0;
    subplot(1,3,1)
    imagesc(SOCIPLITY0);set(gca,'ydir','normal')
    map=colormap('gray');
    map=flipud(map);
    colormap(map);
    ylabel('Evo time')
    xlabel('sociality, c')
    set(gca,'xtick',1:20:res0,'xticklabel',E(1:20:res0))
    subplot(1,3,2)
    plot(DISPREV)
    subplot(1,3,3)
    plot(INFOPREV)
end
```
