## Supplementary figures and images for "Social Information use Shapes the Coevolution of Sociality and Virulence"

### fig1.pdf

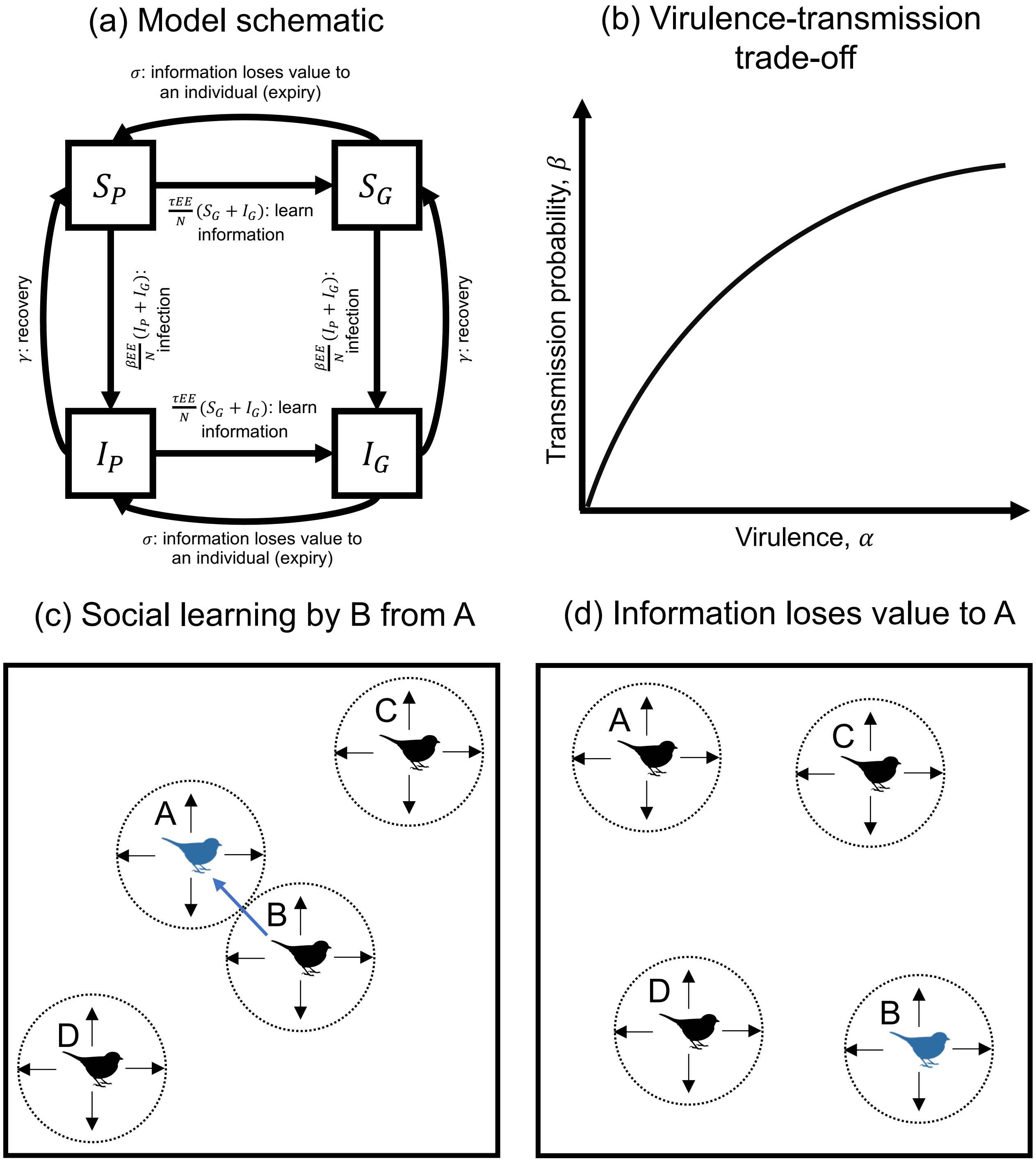

### fig2.pdf

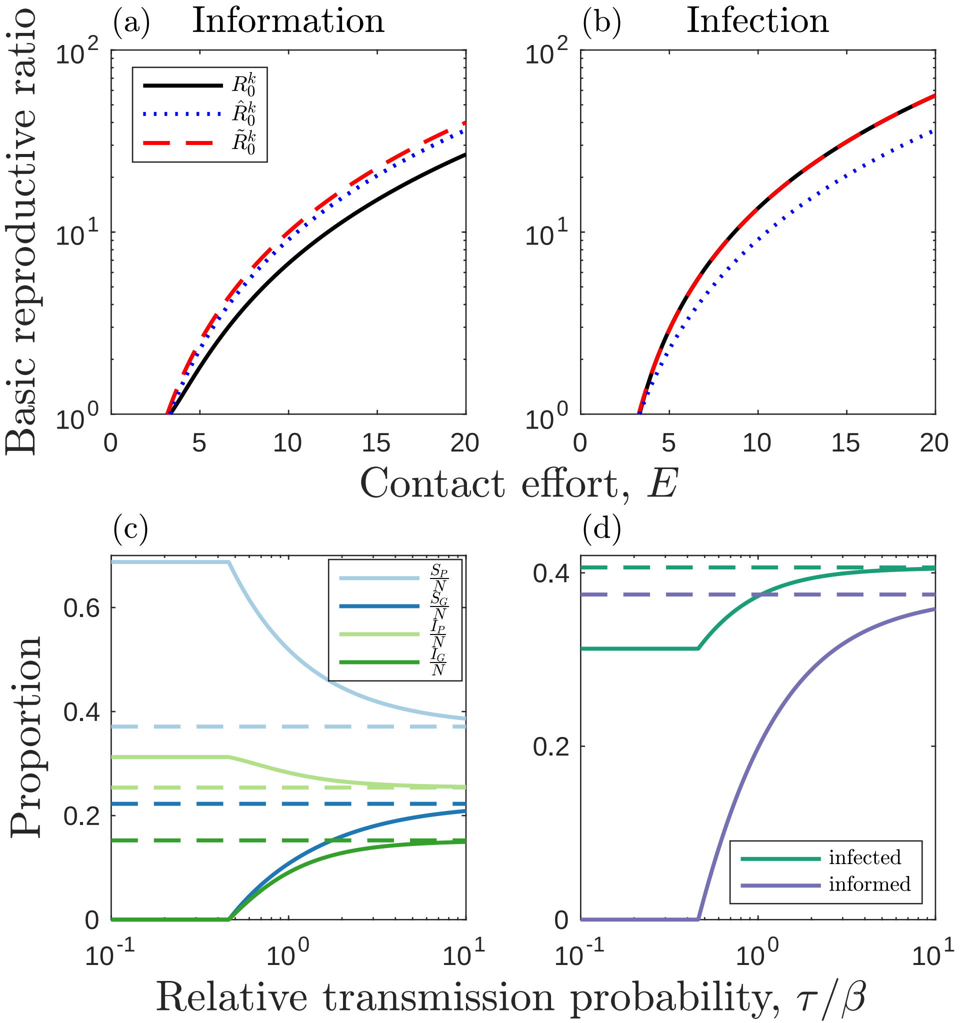

### fig3.pdf

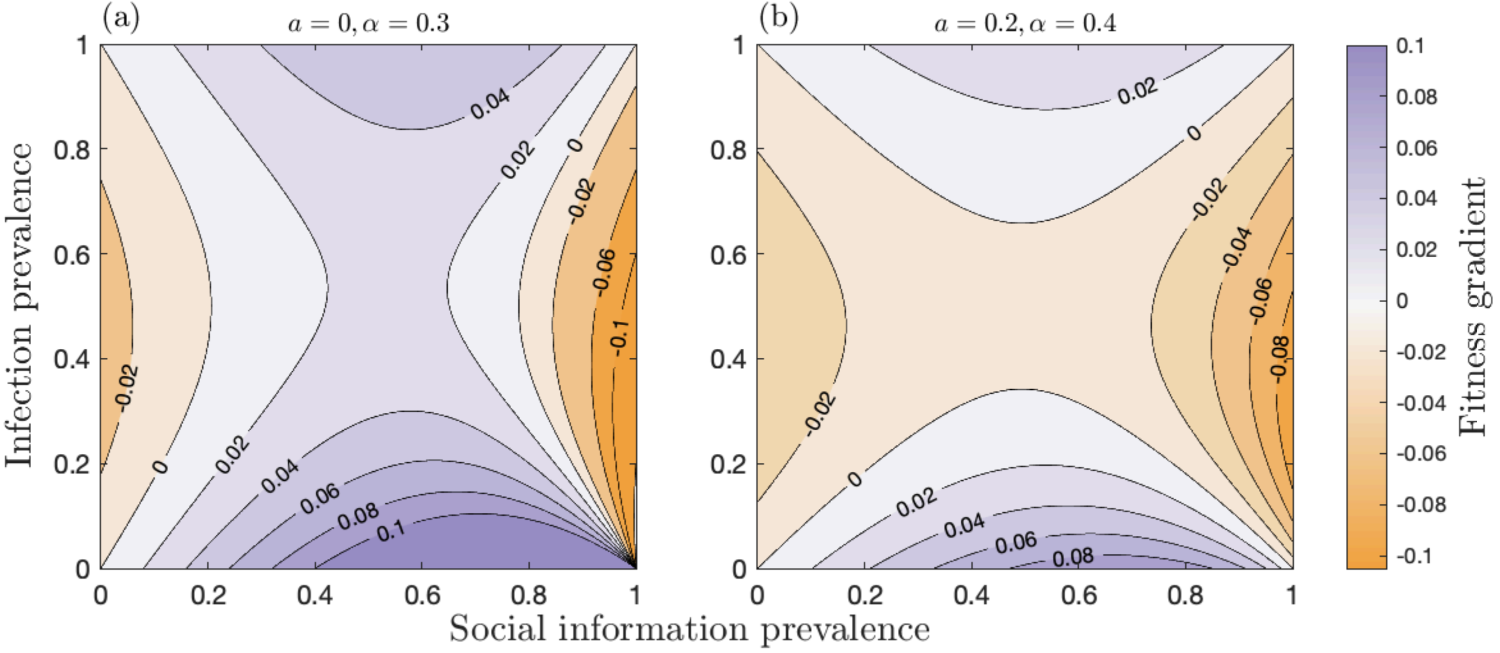

### fig4.pdf

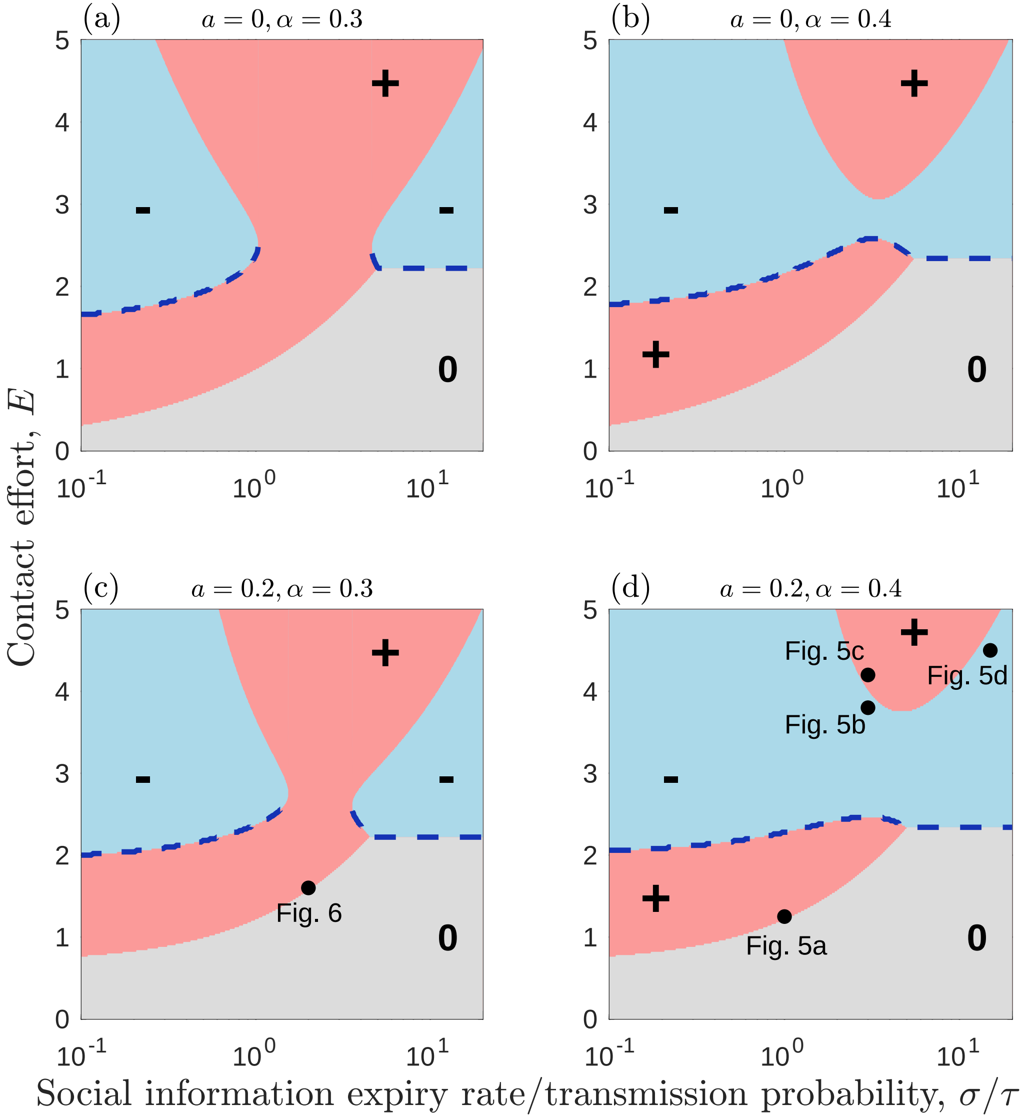

### fig5.pdf

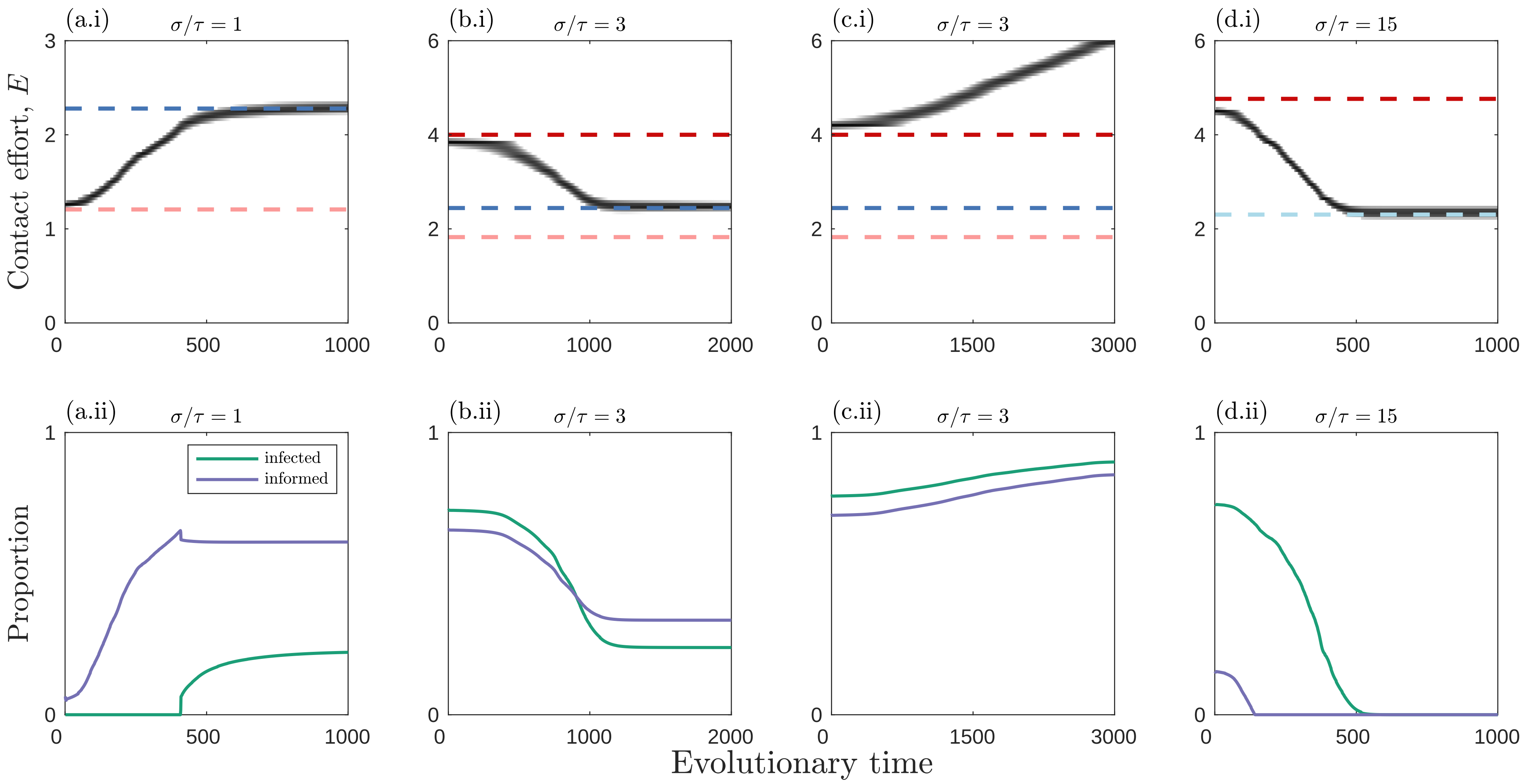

### fig6.pdf

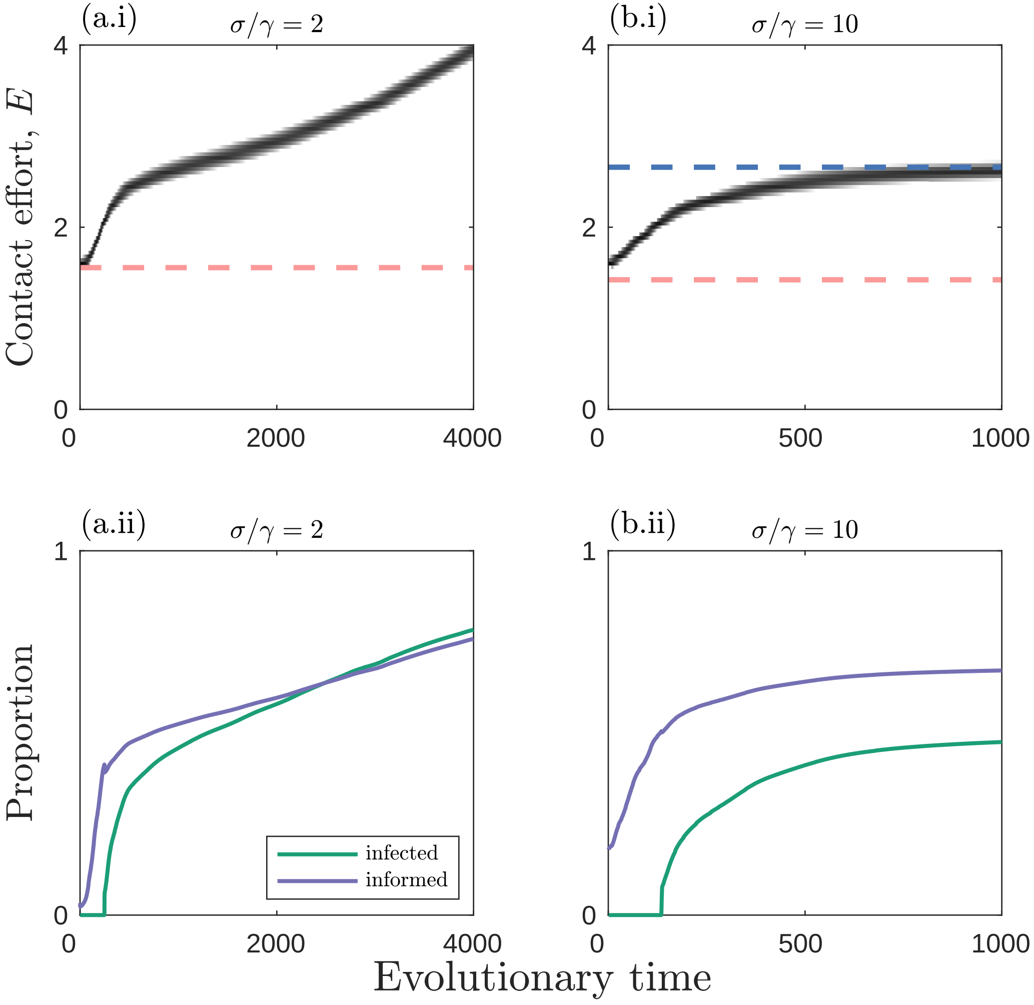

### fig7.pdf

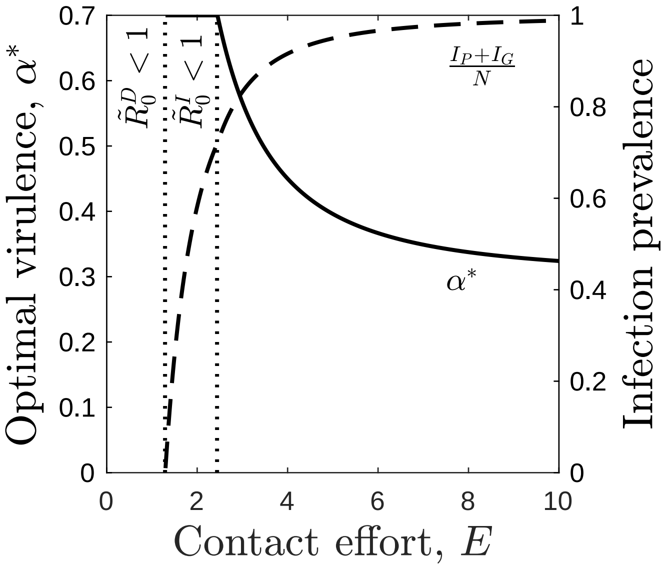

### fig8.pdf

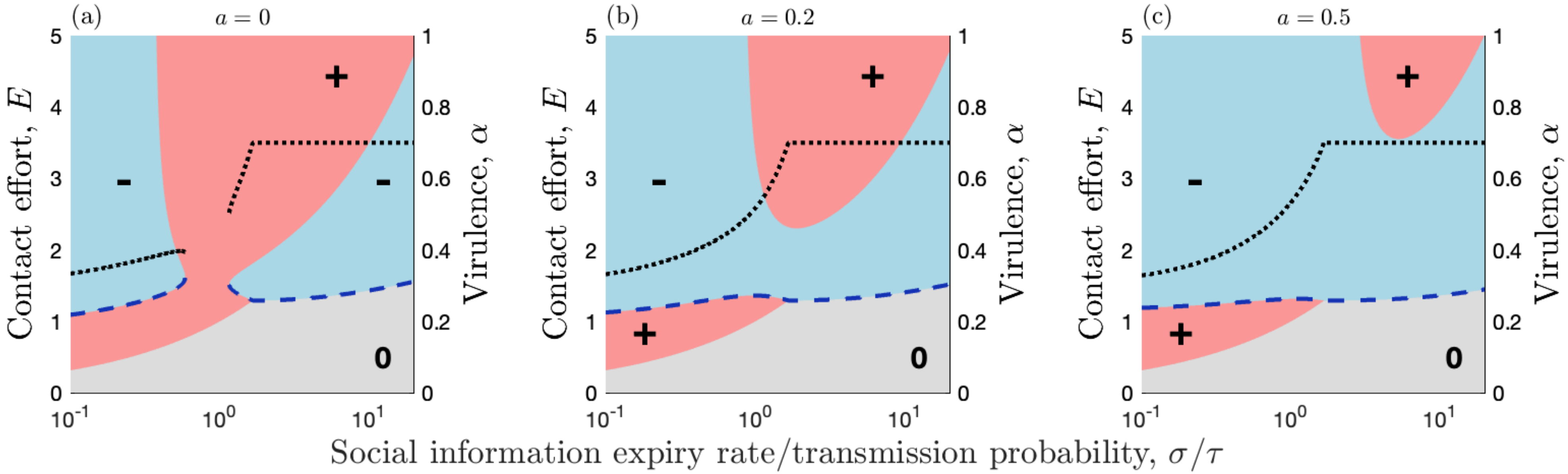

### figS1.pdf

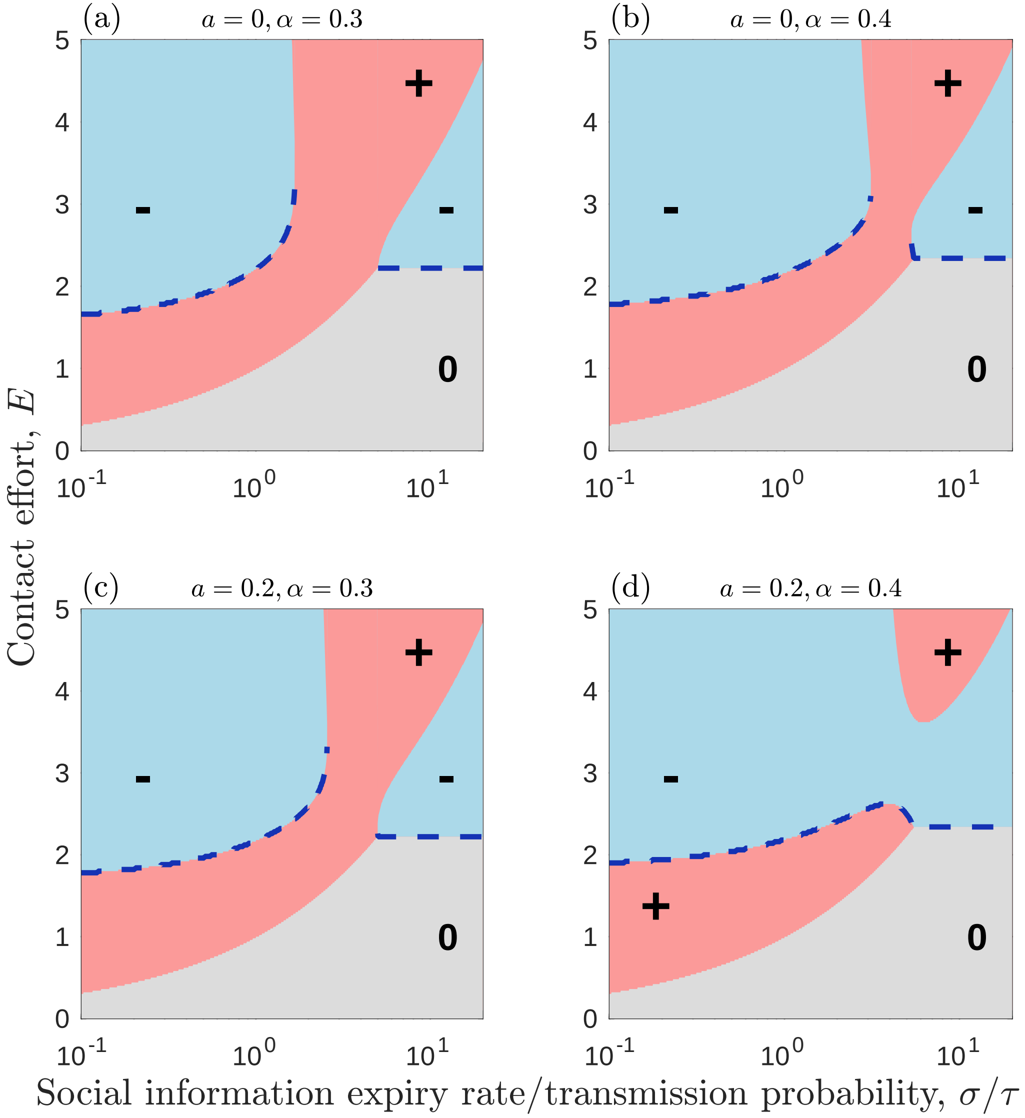
